# Multimodal analysis defines *GNG4* as a distinguishing feature of germinal center-positioned CD4 T follicular helper cells in humans

**DOI:** 10.64898/2025.12.10.693235

**Authors:** Sam Barnett Dubensky, Yutong Zhu, Molly Gallagher, Kingsley Gideon Kumashie, Tianyu Lu, Jonathan Tedesco, Nina De Luna, Katherine Premo, Yi Qi, Suzanna Rachimi, Emylette Cruz Cabrera, Bria Fulmer, Ijeoma C. Meremikwu, Ashley Carter, Sarah E. Henrickson, Neil Romberg, Amy E. Baxter, Derek A. Oldridge, Laura A. Vella

## Abstract

CD4 T follicular helper (Tfh) cells coordinate humoral immune responses within germinal centers (GC) of lymphoid tissue. Despite their critical roles in vaccination and autoimmunity, the gene expression programs that define functionally distinct human Tfh states— and the molecular pathways engaged by Tfh positioned within the GC niche—remain incompletely understood. This gap has limited translational efforts to monitor or therapeutically target specific Tfh states for clinical benefit. Here, we delineate human CD4 T cell heterogeneity in tonsils and peripheral blood using trimodal single-cell sequencing and spectral flow cytometry to define epigenomic, transcriptional, and proteomic features of distinct Tfh states. Tfh with a GC-like phenotype exhibited markedly increased chromatin accessibility and both mRNA and protein expression of G protein subunit gamma 4 (*GNG4)*. In tonsil, single-cell spatial transcriptomics defined *GNG4* expression as a distinguishing feature of activated Tfh states within spatially demarcated GC compartments, with greater specificity than conventionally GC-associated features such as *BCL6, TOX2,* and *S1PR2*. In contrast, *GNG4*^−^ Tfh primarily localized to nonGC regions and exhibited a resting, Th17-polarized phenotype. Together, these data highlight *GNG4* as a central feature of activated, GC-positioned Tfh cell identity in humans.

**One Sentence Summary:** *GNG4* expression defines activated CD4 T follicular helper cells localized to the germinal center of human lymphoid tissue.

## INTRODUCTION

CD4 T follicular helper cells (Tfh) function in secondary lymphoid organs (SLO) where they stimulate proliferation and maturation of B cells within germinal centers (GC) (*1–3*). The GC reaction is tightly coordinated, and the rules that govern human GC output are critical to understand in clinical contexts of vaccination and autoimmunity (*4*). However, translational efforts to target human GC Tfh functions are limited in specificity due to a lack of defined GC-restricted transcriptional programs (*5*). Indeed, many axes of Tfh variation have been reported, including memory versus effector commitment, nonGC versus GC positioning, activation states, and T helper polarization (Tfh1, Tfh2, or Tfh17) (*6–14*). It remains unknown how these axes of variation are integrated into single-cell Tfh states and control of the GC response in humans. Identifying precise Tfh states for clinical intervention therefore requires both deep single-cell and spatial resolution of compartment-dependent gene expression programs in the human system.

Diverse Tfh states have been profiled in human SLOs using single-cell ATAC, RNA, and/or antibody-derived tag (ADT) sequencing (*15–26*). However, because these studies measure at most two modalities at a time, they fall short of providing an integrated view of cell states that unifies chromatin, transcriptional, and protein-level regulation. Recently, TEAseq (simultaneously measuring mRNA Transcripts, surface protein Epitopes, and genome-wide chromatin Accessibility) has enabled deep trimodal molecular analysis at single-cell resolution (*27*). Profiling CD3^+^ peripheral blood mononuclear cells (PBMC) by TEAseq revealed substantial discordance in resolution of non-naive CD4 (nnCD4) T effector memory (Tem) and central memory (Tcm) subsets between protein versus RNA- or ATAC-seq-based definitions. Given that Tfh adopt heterogeneous states including Tem and Tcm-like phenotypes in SLOs and peripheral blood (*9–13, 28–34*), trimodal analysis may thus enhance resolution of functionally significant Tfh states and epigenetic potential in humans.

While Tfh states in blood are conventionally dissected by the same measures used for all CD4 T cells, GC Tfh states are spatially defined by their unique positioning within SLOs (*1–3, 9*). Despite this anatomically grounded definition, dissociated tissue samples are often used to annotate the ‘GC Tfh’ state without spatial context. Flow cytometry-based annotation of GC Tfh in mice using the field-standard definition (CXCR5^hi^ PD1^hi^ Bcl6^+^) was recently found to vastly overestimate counts compared to a histology-based gold standard (*35*). GC-localized Tfh were instead found to be S1PR2^+^ and CD90^-/lo^. However, this dichotomy of ‘GC-like’ versus ‘GC-positioned’ Tfh has not been evaluated in humans. In addition to BCL6, other markers including CD57, TOX2, and ICOS have been used to enrich for ‘GC Tfh’ in humans (*6, 36–40*), but the relative specificity of these features for true GC positioning has not been directly tested. As a result, gene expression programs that are specific to GC-versus nonGC-positioned Tfh states in humans remain undefined.

Here, we dissected Tfh identity from diverse immune cell states in human tonsils and blood. We resolved a group of Tfh states distinct from the greater T cell pool, defined by shared expression of both conventional Tfh features and previously unreported Tfh-associated genes. GC versus nonGC-like states and helper-polarity were linked, with strong Th17-skewing of nonGC-like Tfh in tonsil. Beyond these known axes of Tfh variation, unbiased genome-wide analysis of Tfh variation revealed G protein subunit gamma 4 (Gγ4, encoded by *GNG4*) as a central feature of activated GC-like Tfh states. In spatial analysis, *GNG4* distinguished truly GC-localized Tfh with greater specificity than conventionally GC-associated features such as *BCL6*, *TOX2*, and *S1PR2*. Further, spectral flow cytometry revealed Gγ4 protein enrichment in GC-like Tfh, as well as in Th1- and Th2- versus Th17-polarized subsets of tonsillar Tfh. These data establish *GNG4* as a central feature that distinguishes activated, GC-positioned Tfh cell identity from diverse Tfh states in human lymphoid tissue and peripheral circulation.

## RESULTS

### Epigenomic, transcriptional, and proteomic features variably contribute to immune cell state diversity in circulation and lymphoid organs

To identify multimodal features that distinguish Tfh and CD4 T cell states among greater immune cell diversity across tissues, we performed TEAseq with parallel spectral flow cytometry on tonsil and peripheral blood mononuclear cells from children ages 4-7 years (Fig. 1A, fig. S1, tables S1-6). Live mononuclear cells and CD4 T cells were purified by FACS and labeled with sample barcodes prior to TEAseq (table S2). Dimensionality reduction was performed for all 32,206 cells in each TEAseq modality separately (table S7, Supplementary Methods). Weighted-nearest neighbor analysis (*41*) across all three modalities (3WNN) resolved 15 Level 1 (L1) clusters. These 15 clusters were annotated using lineage-defining features across modalities (fig. S2, data file S1) (*18, 41–46*) and clusters were visualized by Uniform Manifold Approximation and Projection (UMAP) of the 3WNN graph (Fig. 1B). Trimodal clustering resolved expected B and T cell subsets, including a distinct Tfh-like cluster enriched in *CXCR5* [CD185], *PDCD1* [CD279], *BCL6*, *TOX2*, and *IL21* expression (fig. S2). As expected for SLOs, tonsil-derived samples were enriched in Tfh-like cells, germinal center B cells (GCB), and antibody-secreting cells (ASC) relative to peripheral blood (Fig. 1C-D, fig. S3).

**Fig. 1.**
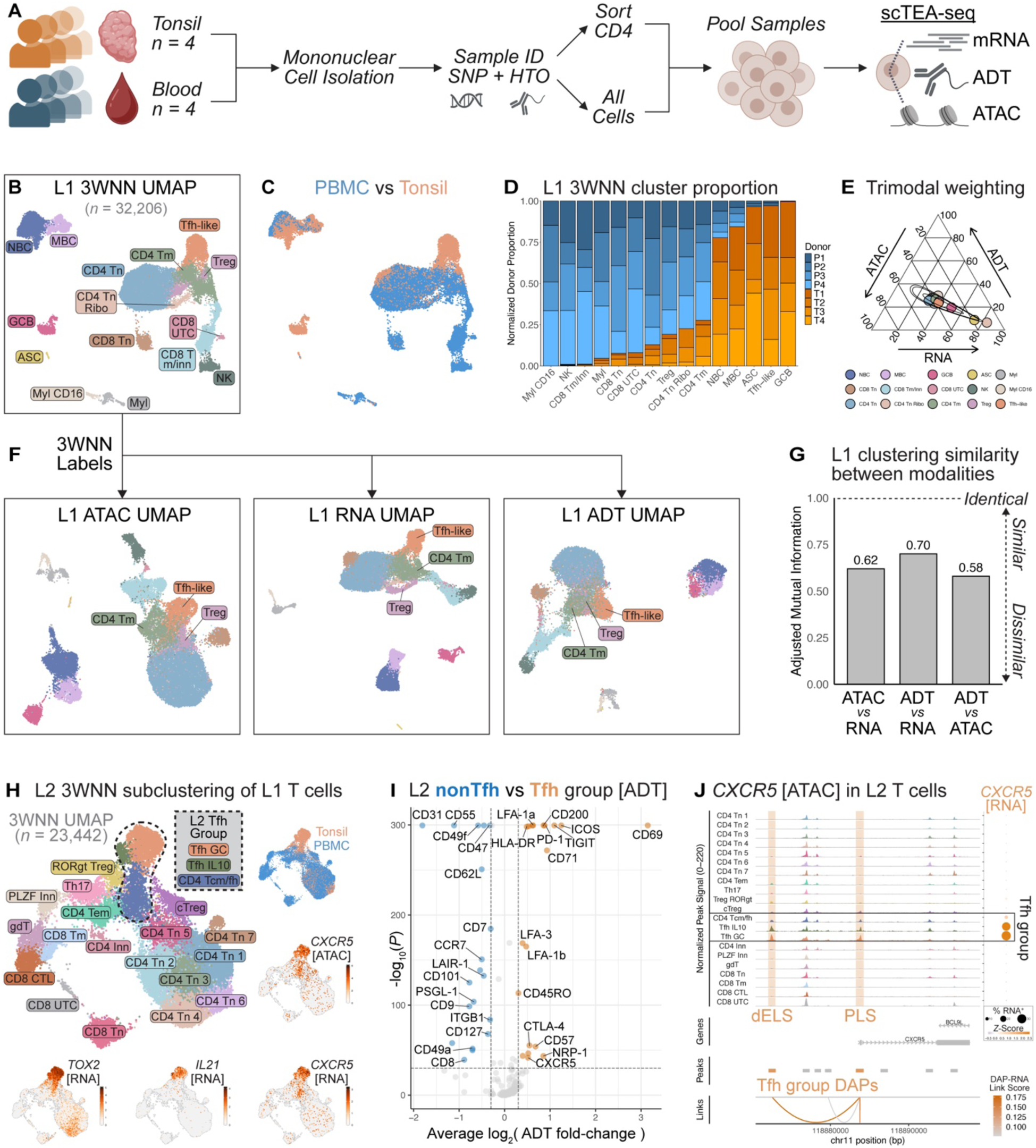
Trimodal analysis resolves distinct Tfh-like states among tonsil and peripheral blood mononuclear cells. **(A)** Experimental schematic for single-cell transcript, epitope, and chromatin accessibility sequencing (TEAseq) of tonsil and peripheral blood mononuclear cells (PBMC). **(B)** Three-way weighted-nearest neighbor (3WNN) UMAP colored by trimodal Level 1 (L1) populations. **(C)** L1 3WNN UMAP colored by tissue type. **(D)** Stacked barplot of unenriched tonsil (T) versus PBMC (P) sample proportions per L1 cluster, normalized to sample cell total. **(E)** Ternary plot of average weights across ATAC, RNA, and ADT modalities for each 3WNN cluster. **(F)** 3WNN Tfh, Treg, and CD4 T memory (Tmem) cluster annotations applied to unimodal UMAP embeddings. **(G)** Barplot of adjusted mutual information (AMI) between clusters resolved by each unimodal analysis. **(H)** 3WNN UMAP embeddings of L2 T cells, with cluster annotations, tissue types, and normalized expression of trimodal features. *CXCR5* [ATAC] represents signal from differentially accessible peak (DAP) containing a promoter-like sequence (PLS). **(I)** Volcano plot of differential ADT expression between grouped L2 Tfh (*n* = 3,657) versus nonTfh (*n* = 19,785) clusters (Wilcoxon rank-sum test *P* < 1e-30, |log_2_FC| > 0.3). **(J)** Normalized *CXCR5* accessibility and RNA expression, pseudobulked per L2 subcluster. DAP enriched in L2 Tfh versus nonTfh clusters included a PLS and a distal enhancer-like sequence (dELS), highlighted by orange bands (Bonferroni-adjusted Wilcoxon rank-sum test *P* < 0.05). Correlation of DAP accessibility with RNA transcript levels is shown by links.

Immune cell identity can be enforced at multiple levels of gene expression (*47–52*), and we reasoned that the 3WNN analysis may differentially weight ATAC, RNA, or protein-based inferences depending on the cell type and tissue origin. We therefore sought to determine the relative contribution of each gene expression modality to immune cell identity in our trimodal data. Most L1 clusters exhibited even weighting between modalities (Fig. 1E & fig. S4), suggesting that all three modalities contributed to lineage identity. However, prior multimodal sequencing studies have demonstrated discrepant T cell states resolved between data types (*41, 53, 54*). To assess agreement across unimodal cluster identities, we first applied 3WNN L1 labels to UMAP embeddings computed using each modality separately (Fig. 1F). Cluster participation was not consistently maintained across modalities. Relative to trimodal analysis, Tfh-like cells in each unimodal embedding exhibited substantial admixture with other CD4 T memory subsets.

To quantify this discrepancy, we resolved and annotated 15 clusters for each modality separately (fig. S5, data file S1) and computed the adjusted mutual information (AMI), which measures cluster assignment similarity per cell regardless of cluster naming (*55–57*). Comparisons with ATAC-based clustering were the most dissimilar, suggesting that immune cell identity defined by chromatin accessibility does not consistently match identity defined by the transcriptome and proteome, even at coarse L1 resolution (Fig. 1G). Given this contrast, we maintained all three modalities for downstream analyses of heterogeneity within the Tfh pool across tissues.

### Tfh exhibit multiple separate yet related gene regulatory states distinct from the greater T cell compartment

Tfh in circulation and tissue can exhibit phenotypes reflecting other major CD4 T cell subsets, including central (Tcm) and effector memory (Tem)-like cells (*1–3*). To resolve Tfh gene expression states distinct from other T cell lineages, trimodal subclustering of L1 T cells was performed as a deeper L2 analysis (Fig. 1H). Among 23,442 T cells, we resolved and manually annotated 21 L2 T cell clusters, including major known CD4 and CD8 T cell subsets (fig. S6, data file S2, Supplementary Methods) (*18, 41, 46*). Even within this broad T cell analysis, we resolved a group of three Tfh-like clusters defined by increased expression of *CXCR5* (Fig. 1H, fig. S7, data file S2). Two of these Tfh-like subsets, ‘Tfh GC’ and ‘Tfh IL10’, were found almost exclusively in tonsil, and both expressed GC-associated features (*TOX2, IL21,* CD57) (Fig. 1H, fig. S7C). ‘Tfh IL10’ were distinguished from ‘Tfh GC’ by features matching an IL10-producing FOXP3^−^ Tfh state (*IL10*, *PRDM1*, CD25; potentially reflecting ‘CD25^hi^ Tfh’ (*22*)) that has been characterized by several groups (*21–25, 58–63*). We also resolved a tonsil-biased (∼80%, fig. S7C) cluster with intermediate expression of both Tfh (*CXCR5, BCL6, TOX2*) and Tcm (*ANXA1, IL7R, ANK3*) signature features (fig. S6, data file S2) (*18, 41*). Given this hybrid phenotype and tissue distribution, we annotated this third Tfh-like 3WNN cluster as ‘CD4 Tcm/fh’. Relative to other nnCD4 clusters, Tcm/fh exhibited decreased *KLF2* [ATAC/RNA] and PSGL-1 [ADT], further suggesting a Tfh-like phenotype (fig. S6, data file S2) (*9, 64, 65*). Relative to nonTfh, all three Tfh-like clusters shared accessibility at 415 peaks near 319 unique genes and expression of 225 transcripts (fig. S7E, data file S2), suggestive of a common gene expression program. Further, L2 Tfh exhibited greater CXCR5 and PD1 protein expression relative to nonTfh (Fig. 1I, data file S2), with decreased PSGL-1, CD62L, and CCR7. Together, these trimodal data indicated that despite substantial heterogeneity, Tfh adopt a shared gene regulatory state distinct from diverse T cell states in circulation and tonsil.

The identification of single-cell Tfh clusters defined by paired epigenomic and transcriptional information allowed for direct nomination of *cis*-regulatory elements (CRE) that may control expression of CXCR5, the signature Tfh feature across diverse Tfh states. Previous studies have revealed enhancer elements of *CXCR5* specific to GC-like Tfh versus nonTfh effector and naive CD4 T cells defined as bulk populations from tonsil (*66, 67*). However, our data enabled fine resolution of CRE that maintain CXCR5 expression across a diverse spectrum of Tfh states, spanning SLO tissue and peripheral circulation, and with a wider array of contrasting cell types.

To identify Tfh-specific CRE that drive *CXCR5* expression, we evaluated distinct ATAC peaks near the *CXCR5* locus across T cell subclusters for correlated accessibility with transcript levels. Among differentially accessible peaks (DAP) enriched in the L2 Tfh group, two resided within distal enhancer (dELS) and promoter-like sequences (PLS) of the *CXCR5* locus (Fig. 1J, data file S2) (*68*). While some regions of *CXCR5* were generally accessible across all T cells, the two Tfh-enriched DAPs directly correlated with *CXCR5* RNA expression. Given that these *CXCR5* dELS are shared by distinct Tfh states, yet relatively inaccessible to other subsets of the diverse T cell pool in human SLOs and circulation, these data suggest Tfh identity may be driven by distinct regulatory elements located far from known promoter regions of the linear genome.

### GC versus nonGC-like Tfh states are skewed in helper-polarity phenotype

Multiple subsets of the human Tfh compartment have been reported, including GC versus nonGC, Tem versus Tcm, and ‘polarized’ (Tfh1 versus Tfh2 and Tfh17) states (*6–9*). However, it is unclear how these independently defined axes of differentiation and helper-polarity intersect in human Tfh. To understand how these major known axes of Tfh variation relate, we first performed 3WNN subclustering analysis of all *CXCR5*-enriched L2 Tfh clusters, paired with flow cytometric profiling of the same samples (Fig. 2A, fig. S1, table S5-6). We annotated nine Level 3 (L3) 3WNN clusters based on differential ATAC, RNA, and ADT feature expression (Fig. 2B, fig. S8, data file S3). Five Tfh clusters exhibited a GC-like phenotype distinguished by expression of *BCL6, TOX2,* and *IL21* [ATAC/RNA] as well as PD1, ICOS, and TIGIT [ADT]. Except for Tfh-IL10, GC-like clusters were sparse in PBMC samples, further suggestive of GC localization (Fig. 2C-D). The remaining four nonGC-like Tfh clusters had increased *KLF2* and CD127 expression, consistent with unstimulated states outside the GC (Fig. 2E, fig. S8) (*1, 12, 24, 25, 32*). Differential TIGIT versus CD127 protein expression across GC versus nonGC-like groups, respectively, resembled TIGIT^+^ GC-committed versus CD127^+^ Tcm-like Tfh states recently described in mice (*13*).

**Fig. 2.**
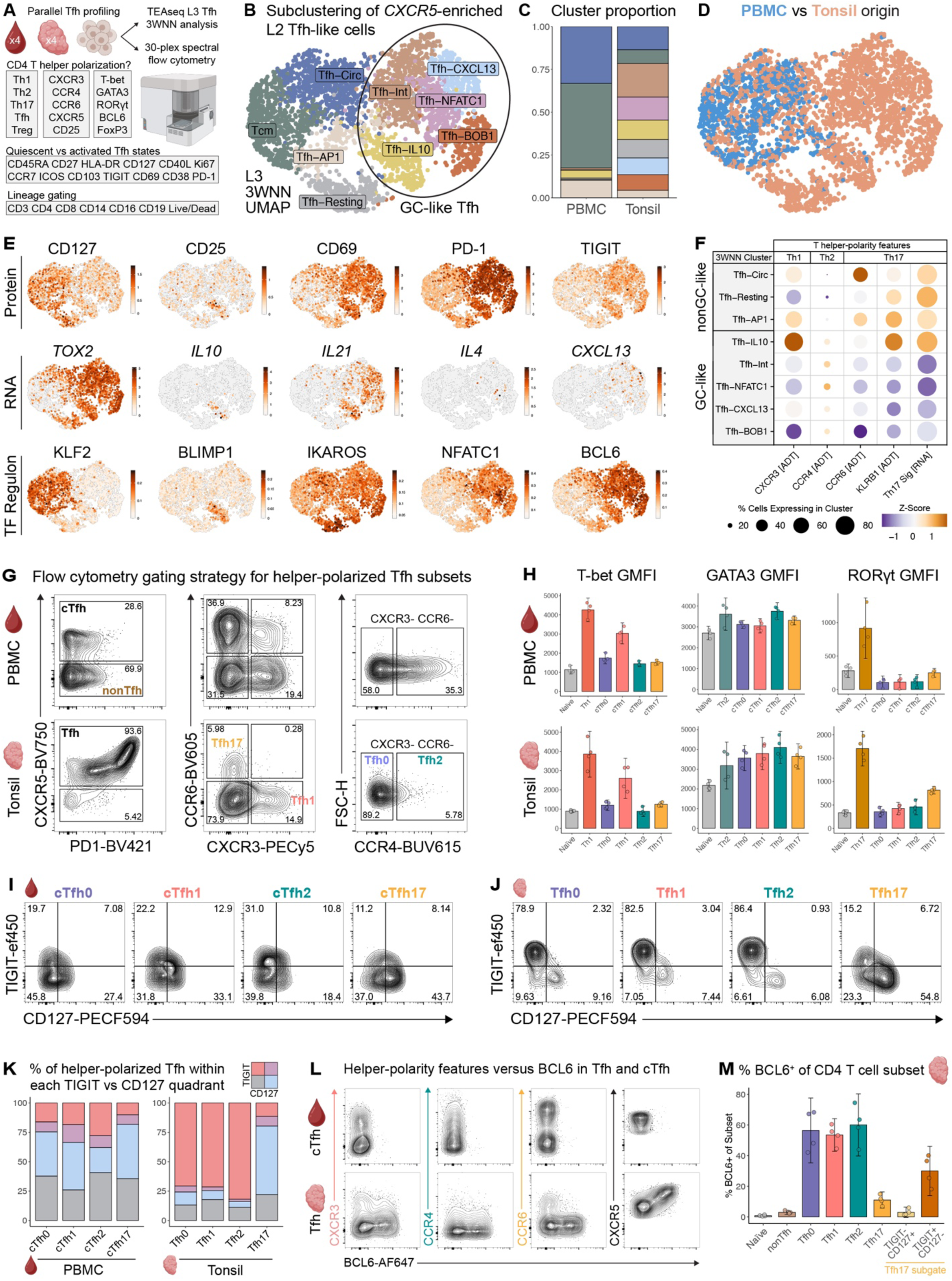
GC versus nonGC-like Tfh states are skewed in helper-polarity phenotype. **(A)** Schematic of parallel spectral flow cytometric and TEAseq Level 3 (L3) three-way weighted-nearest neighbor (3WNN) subclustering analysis of Tfh-like cell states in tonsil and PBMC samples. **(B)** 3WNN UMAP of L3 subclusters from L2 Tfh group. **(C)** Barplot of L3 subcluster proportion per tissue. **(D)** 3WNN UMAP colored by tissue type. **(E)** 3WNN UMAPs of differentially expressed SCENIC regulons, RNA and ADT features. **(F)** Scaled average expression and percent expression of T helper polarity-associated ADT features and Th17 RNA signature across subclusters. **(G)** Gating of helper-polarized tonsil Tfh versus circulating Tfh (cTfh) subsets. **(H)** Barplots of TF GMFI across nnCD4 subsets in tonsil and PBMC (*n* = 4 each, unpaired) with 95% confidence intervals (CI) around subset means. **(I)** Gating of TIGIT^+^ vs CD127^+^ subsets of polarized Tfh in blood and **(J)** tonsil. **(K)** Stacked barplot of average TIGIT vs CD127 subset percentages per helper-polarized Tfh subset. **(L)** Representative expression of BCL6 versus chemokine receptors in Tfh and cTfh. **(M)** Barplot of BCL6^+^ percentage across tonsil CD4 T cell subsets with 95% CIs around subset means. Tfh17 subsets were further gated by TIGIT^−^CD127^+^ and TIGIT^+^CD127^−^ phenotype.

To identify transcription factors (TF) that may drive functional differences between GC and nonGC Tfh states, we first performed SCENIC analysis to resolve TF regulons (Fig. 2E bottom row, fig. S9A, data file S3, Supplementary Methods) (*69*). Regulons including BCL6, NFATC1, and IKAROS were enriched in GC-like Tfh broadly, in contrast to KLF2, an inhibitor of Tfh differentiation in mice (*65*). Further, chromVAR analysis of GC-like Tfh epigenomic states discerned enriched TF motif accessibility of ASCL2, MAF, and several members of the NFAT and POU families (fig. S9B, data file S3), again in contrast to KLF2 (*70*). However, each state was associated with a distinct pattern of TF activity including many unreported potential regulators such as STAT2 in *IL10*^+^ and ELF2 in *CXCL13*^+^ Tfh (fig. S9). Together, these data suggest that GC and nonGC Tfh groups comprise multiple finer gene expression states with distinct networks of coordinated TF activity.

Given the marked diversity within GC and nonGC Tfh compartments, we next asked if Tfh from tonsils exhibited features along the T helper polarity axis. Peripheral blood circulating Tfh (cTfh) can be classified along this axis based on expression of chemokine receptors, with cTfh1 defined as CXCR3^+^CCR6^−^, cTfh2 as CXCR3^−^CCR6^−^ and cTfh17 as CXCR3^−^CCR6^+^ (*7, 11*). Polarized Tfh subsets exhibit distinct, specialized functions, such as enhanced B cell class-switching towards specific isotypes (*15, 71*). To determine how the GC versus nonGC-like programming axis (Fig. 2B) relates to helper-polarization in Tfh, we next leveraged ADT expression of CXCR3, CCR4, and CCR6 in our TEAseq data (Fig. 2F). GC-like Tfh were distinguished by increased CCR4 expression and decreased CXCR3 and CCR6, suggestive of Tfh2 polarity. In contrast, the nonGC-like group of Tfh were enriched in CCR6, suggestive of enriched Tfh17 polarity. Further, KLRB1, a marker of Th17 polarity in humans (*72–77*), was enriched across nonGC-like Tfh states as was a transcriptional signature of human Th17 differentiation (table S8) (*76–87*). Of note, Tfh-IL10 were enriched in both KLRB1 and Th17 signature genes, but not CCR6, suggesting that expression of CCR6 alone may not resolve all Th17-polarized Tfh.

To determine whether the polarity distribution we observed in GC- versus nonGC-like Tfh states reflects the same polarization biology originally defined in peripheral blood (*7*), we used flow cytometry to resolve Tfh polarity across tissues. As expected, cTfh could be further detailed as cTfh1, cTfh17, and cTfh2-enriched subsets (Fig. 2G, fig. S10A-B). In tonsil, Tfh exhibited similarly discrete chemokine receptor expression to cTfh, though CXCR3^+^CCR6^+^ Tfh1/17 were sparse (Fig. 2G, fig. S10C). However, Tfh2 polarization within the CXCR3^−^CCR6^−^ population differed by tissue: only ∼5% of CXCR3^−^CCR6^−^ Tfh expressed the Th2-associated chemokine receptor CCR4 in tonsil, compared to ∼35% of CXCR3^−^CCR6^−^ cTfh in blood (Fig. 2G, fig. S10C). Together, these data indicate that polarity-defining chemokine receptors are differentially expressed between Tfh in SLOs versus peripheral circulation, suggestive of compartment-dependent Tfh polarity programming.

Given the differences in chemokine receptor patterns in tonsil and blood, we next asked whether the polarity of Tfh as determined by lineage-defining transcription factor expression differed by tissue. Prior studies demonstrated that cTfh exhibit discrete RNA expression of polarity-defining TF, including T-bet in cTfh1, GATA3 in cTfh2, and RORγt in cTfh17 (*7*), but simultaneous protein-level staining of these TFs in human Tfh has not been reported. We therefore used TF protein expression to define polarity differences in Tfh between tissues. Th1 and Tfh1 expressed the most T-bet relative to other polarized Tfh subsets (Fig. 2H, fig. S10D). Likewise, RORγt was higher in Th17 and Tfh17 than other subsets, but Tfh17 displayed lower RORγt than their T helper counterparts. In contrast, GATA3 protein was broadly expressed across helper-polarized Tfh subsets in tonsil and peripheral blood, with marginally increased GMFI demonstrated in cTfh2 in blood and Tfh2 in tonsil. Together, these data suggest that Tfh in SLOs and peripheral blood adopt distinct states programmed by differential activity of polarity-defining TF proteins and that these states are preserved across tissues. However, the muted expression of TFs in Tfh compared to conventional T helper subsets suggests that Tfh may be less definitively polarized, potentially with retained and enhanced plasticity.

We next asked if the relationship between polarization and GC- versus nonGC-like Tfh observed in Fig. 2F was maintained in flow cytometric space. Using CD127 and TIGIT to denote nonGC-versus GC-like Tfh, respectively, we observed that cTfh0, cTfh1, and cTfh17 in peripheral blood, as well as Tfh17 in tonsil were most enriched in CD127^+^TIGIT^−^ phenotype (∼38-46% Fig. 2I-K, fig. S10E-F), consistent with Tcm-like, nonGC programming (*13*). In contrast, blood cTfh2 were more enriched in CD127^−^TIGIT^+^ phenotype than other polarized cTfh subsets (∼28%, Fig. 2K, fig. S10E), consistent with enhanced GC effector potential (*13*). Finally, we assessed the co-expression of BCL6 with each polarization state. Tfh17 expressed less BCL6 protein than other polarized Tfh in tonsil tissue, suggesting that Tfh17 primarily localize outside the GC (Fig. 2L-M, fig. S10G). However, TIGIT^+^ Tfh17 cells were more likely to express BCL6 (Fig. 2M, fig. S10G), revealing potential positional or functional heterogeneity within the Tfh17 compartment. Together, these data suggest that GC versus nonGC Tfh programming is skewed by helper-polarity: GC-like Tfh were enriched in Th2-polarity, whereas nonGC-like Tfh primarily adopted a Th17-polarized state.

### *GNG4* (G protein subunit gamma 4) expression distinguishes GC-like Tfh states in tonsil

In mice, relative expression of CD90 versus S1PR2 by Tfh is tightly associated with GC-localization, aiding identification of GC Tfh *ex vivo* (*35*). However, the molecular program that specifically identifies Tfh located within the GC in humans is not well defined. To identify candidate features of bona fide human GC Tfh, we leveraged our multimodal data to compare genome-wide chromatin accessibility and transcription in GC versus nonGC-like Tfh groups (Fig. 3A-C, data file S3). This analysis identified differentially accessible genes (DAG) and differentially expressed genes (DEG) known to be enriched in GC-like Tfh, including *PDCD1, TOX2,* and *B3GAT1* (Fig. 3B-C). We then performed a meta-analysis of loci detected in both RNA and ATAC assays to identify features enriched in GC-like Tfh across modalities (Fig. 3D, data file S3). Strikingly, G protein subunit gamma 4 (Gγ4, encoded by *GNG4*) was among the most distinguishing features of GC-like Tfh, with greater multimodal enrichment relative to nonGC-like states than *TOX2, BCL6*, and *B3GAT1*. G proteins mediate fundamental processes in T cells such as migration and activation by regulating G protein-coupled receptor (GPCR) signaling (*88–98*). We observed that *GNG4* transcripts were largely absent in the nonGC Tfh population, as shown by 3WNN UMAP (Fig. 3E), indicating that *GNG4* expression is highly enriched in the GC Tfh state. *GNG4* expression was also sparse in other L1 mononuclear cell and L2 T cell subsets within our TEAseq dataset (Fig. 3F–G). Together, these findings suggest that *GNG4* may be preferentially engaged in GC Tfh, separating GC from both nonGC Tfh and other lineages in SLOs more broadly.

**Fig. 3.**
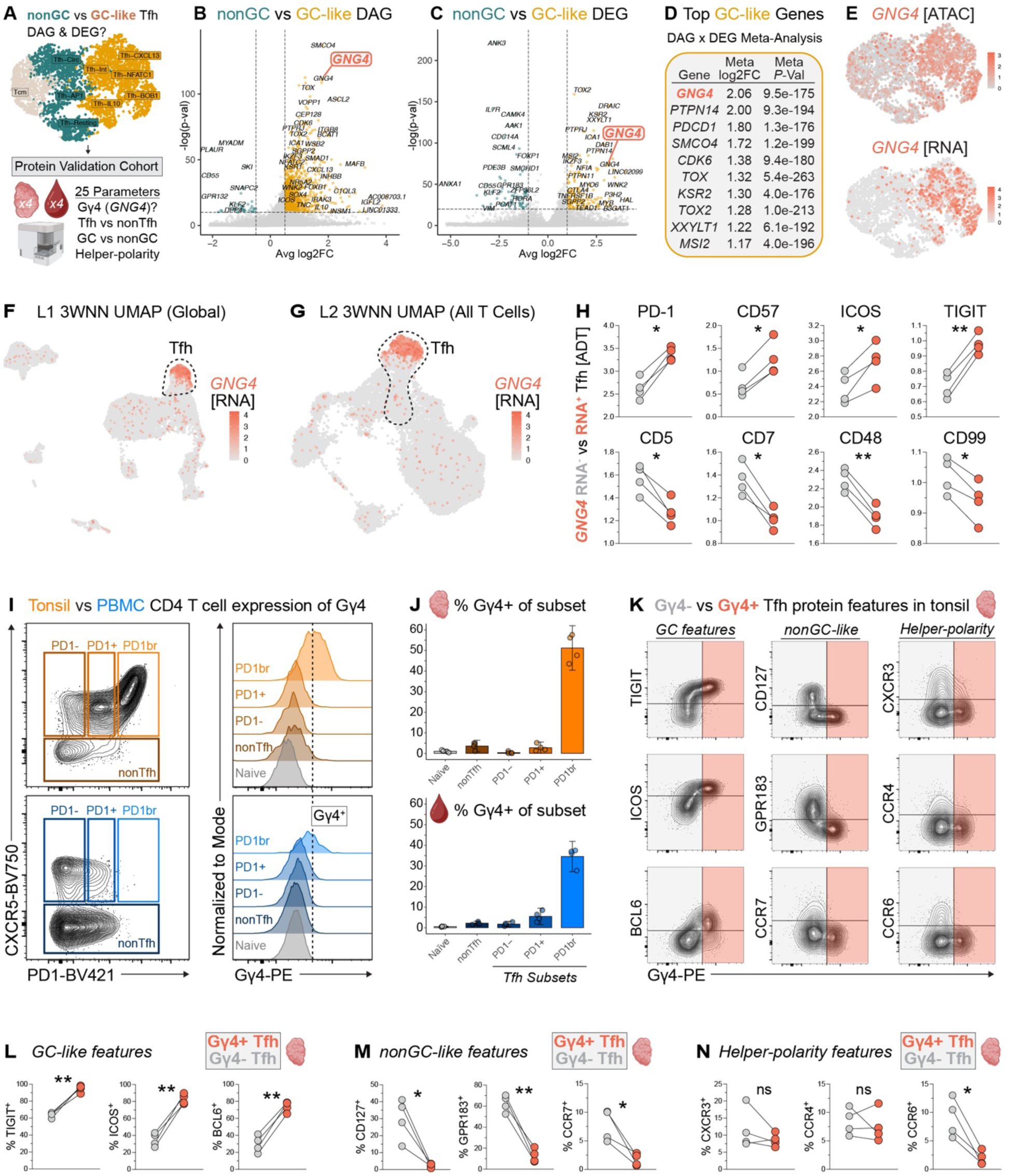
Multimodal *GNG4* expression distinguishes activated GC-like Tfh cell states. **(A)** Schematic of TEAseq and flow cytometry analysis approach to resolve differentially accessible (DAG, from ATAC GeneActivity analysis) and expressed genes (DEG, from normalized RNA data) in GC versus nonGC-like Tfh states. Volcano plots comparing GC- vs nonGC-like Tfh show **(B)** DAGs (Wilcoxon rank-sum test *P* < 1e-10 & |log_2_FC| > 0.5) and **(C)** DEGs (Wilcoxon rank-sum test *P* < 1e-20 & |log_2_FC| > 1). X-axis ranges for panels B and C exclude only non-significant features. **(D)** Top 10 features enriched in the GC-like Tfh group from meta-analysis of all DAGs and DEGs (lowest meta *P-*value genes ranked by meta log_2_FC). **(E)** 3WNN UMAP showing *GNG4* accessibility (ATAC GeneActivity) and normalized RNA expression. **(F)** Normalized *GNG4* RNA expression across all mononuclear cells in L1 TEAseq analysis and **(G)** all T cells in L2 TEAseq subclustering analysis. **(H)** Mean normalized ADT expression in *GNG4* RNA^+^ vs RNA^−^ Tfh per tonsil donor (paired *t*-test with two-stage step-up procedure of Benjamini, Krieger, and Yekutieli, FDR-adjusted \**Q* < 0.1, \*\**Q* < 0.05). **(I)** Representative gating strategy and histogram of Gγ4 expression in Tfh, naive CD4 T cells, and FOXP3^−^CXCR5^−^ nonTfh cells. Vertical line indicates threshold for positive Gγ4 expression defined using peripheral blood naive CD4 T cells as an internal negative control. **(J)** Barplots of % Gγ4^+^ per CD4 T cell subset and tissue. 95% confidence intervals surrounding each subset mean are shown. **(K)** Representative staining of GC-like, nonGC-like, and helper-polarity features plotted against Gγ4 in tonsil Tfh. Shaded boxes indicate Gγ4^+^ (coral) versus Gγ4^−^ (grey) subsets of tonsil Tfh. **(L)** Mean percentage of Gγ4^+^ versus Gγ4^−^ tonsil Tfh expressing GC-like, **(M)** nonGC-like, and **(N)** helper-polarity features (paired *t*-test with Holm-Šídák correction for multiple comparisons, \**P* < 0.05, \*\**P* < 0.01).

Although its role in Tfh biology has been underappreciated to date, *GNG4* expression has appeared in DEG analyses of human T cells, including Tfh, thymic CD8⍺⍺ T cells, cancer-associated subsets, and was listed after mass spectrometry studies of CD4 T cells stimulated *in vitro* (*14, 25, 43, 67, 92, 99–107*). To determine the molecular programs associated with *GNG4*, we compared *GNG4* RNA^+^ versus *GNG4* RNA^−^ Tfh. *GNG4* RNA^+^ Tfh were highly enriched in protein expression of PD1, CD57, TIGIT, and other features associated with GC Tfh (Fig. 3H, data file S3) (*9, 13*). In contrast, *GNG4* RNA^−^ Tfh exhibited greater CD5, CD7, and CD47 protein expression, suggestive of a less activated and undifferentiated Tfh state (*108–113*). Further, *GNG4* RNA^−^ Tfh had increased CD99 expression, a recently identified feature of cTfh and nonGC-like states (*15*). Together, these data suggest that *GNG4* expression marks a highly activated subset of GC-like Tfh.

### Gγ4 protein expression is enriched in activated GC-like Tfh

We next asked if GC vs nonGC Tfh states could be distinguished based on expression of Gγ4 protein. We conjugated human Gγ4-specific antibodies (table S5-6) and analyzed Gγ4 expression in an additional cohort of four pediatric tonsil and four adult PBMC donors (table S1). Naive CD4 T cells were used as an internal biological negative control for Gγ4 protein expression (Fig. 3I-J, fig. S11A-B) given the lack of *GNG4* accessibility and transcription in naive CD4 T cells defined by TEAseq (fig. S8A-B). Gγ4 expression was low or absent in nonTfh as well as in Tfh with absent or intermediate PD1 expression, similar to naive CD4 T cells (Fig. 3J, fig. S11C-D). In contrast, PD1^br^ Tfh from both tonsil and PBMC were highly enriched in Gγ4 expression. Both the PD1^br^ and Gγ4^+^ subsets of Tfh were far more common in tonsil than in blood (fig. S11E-F), consistent with PD1-ADT and *GNG4* RNA-based definitions of Tfh in our TEAseq analysis (fig. S11G-H). Expression of Gγ4 in these rare PD1^br^ cTfh may reflect accessibility and transcription of *GNG4* in PD1^br^ cTfh, a GC-like subset thought to emigrate from SLOs (*9, 14, 114*).

To infer the biological function of Tfh expressing Gγ4 protein, we first compared Gγ4^−^ versus Gγ4^+^ Tfh in tonsil by known features of GC-like, nonGC-like and helper-polarized states (Fig. 3K). GC-associated Tfh features were more frequently expressed in the Gγ4^+^ rather than Gγ4^−^ subset of tonsil Tfh (∼95.0 versus 63.9% TIGIT^+^, 82.8 versus 35.0% ICOS^+^, 73.6 versus 29.7% BCL6^+^, Fig. 3L). In contrast, a lower frequency of Gγ4^+^ Tfh expressed proteins associated with unstimulated and nonGC-like states (∼2.3 versus 29.9% CD127^+^, 12.7 versus 62.5% GPR183^+^, 1.7 versus 7.7% CCR7^+^, Fig. 3M) (*9, 13, 115*). As functionally mature Tfh can differentiate into Tfr and mediate suppressive functions within GCs, we next considered how Gγ4 and FOXP3 protein expression relate (*22, 116–119*). Among all CXCR5^+^ nnCD4 T cells in tonsil, FOXP3 was more frequently expressed in the Gγ4^−^ rather than Gγ4^+^ subset (∼2.92 versus 0.95% FOXP3^+^, fig. S11I). However, the PD1^br^ subset of FOXP3^+^ CD25^+^ Tfr was enriched in Gγ4 expression relative to PD1^−^ and PD1^+^ Tfr, though not to the extent of PD1^br^ Tfh (fig. S11J). Together, these data suggest that Gγ4 can be expressed by both activated, GC-like Tfh and Tfr.

Having shown that Tfh express Gγ4 in GC-like states, as well as features of T helper polarity in tonsil tissue, we therefore asked whether expression of Gγ4 varies between polarized Tfh states. Gγ4^+^ Tfh had decreased expression of the Th17-associated chemokine receptor CCR6 compared to Gγ4^−^ Tfh (∼1.9% versus 9.0%, Fig. 3N), consistent with the prior finding that Tfh17 are enriched in nonGC-like TIGIT^−^CD127^+^ phenotype (Fig. 2K, fig. S10F). In contrast, CXCR3 and CCR4 did not vary with Gγ4 expression. Indeed, only ∼7% of Tfh17 were Gγ4^+^, in contrast to ∼33-38% of Tfh0, Tfh1, and Tfh2 subsets (fig. S11). Together, these data indicate that Gγ4 protein expression delineates GC Tfh identity from diverse nonGC, regulatory, and Th17-polarized states of the Tfh compartment.

### *GNG4* expression is associated with activated Tfh states positioned within the GC light zone

Our single-cell data suggested that *GNG4* marks GC-like Tfh; however, these data lack spatial resolution. To evaluate the fidelity of *GNG4* and other transcripts to identify Tfh within true GCs, we profiled tonsil tissue (*n* = 6 donors, table S1) using the 10x Genomics Xenium Prime (XP) single-cell spatial transcriptomics assay (Fig. 4A). We supplemented the 5001-plex base panel with 100 custom probes (table S9), including *GNG4*, *KLF2*, *GPR183*, *IL7R*, and *RORA* and other transcripts nominated by our trimodal analysis. To identify anatomic GCs, we co-registered H&E and DAPI images for each sample (Fig. 4B-C, fig. S13, data file S4). Canonically GC-associated transcripts such as *BCL6* and genes expressed by light zone (LZ) versus dark zone (DZ) GCB cells—including *LMO2* and *AICDA*, respectively—were enriched in anatomically defined GCs, although *BCL6* was observed outside of the GC and in the DZ as expected (Fig. 4D-E) (*120*). In contrast, *GNG4* expression was more spatially restricted to the GC (Fig. 4F), particularly within the LZ (Fig. 4G).

**Fig. 4.**
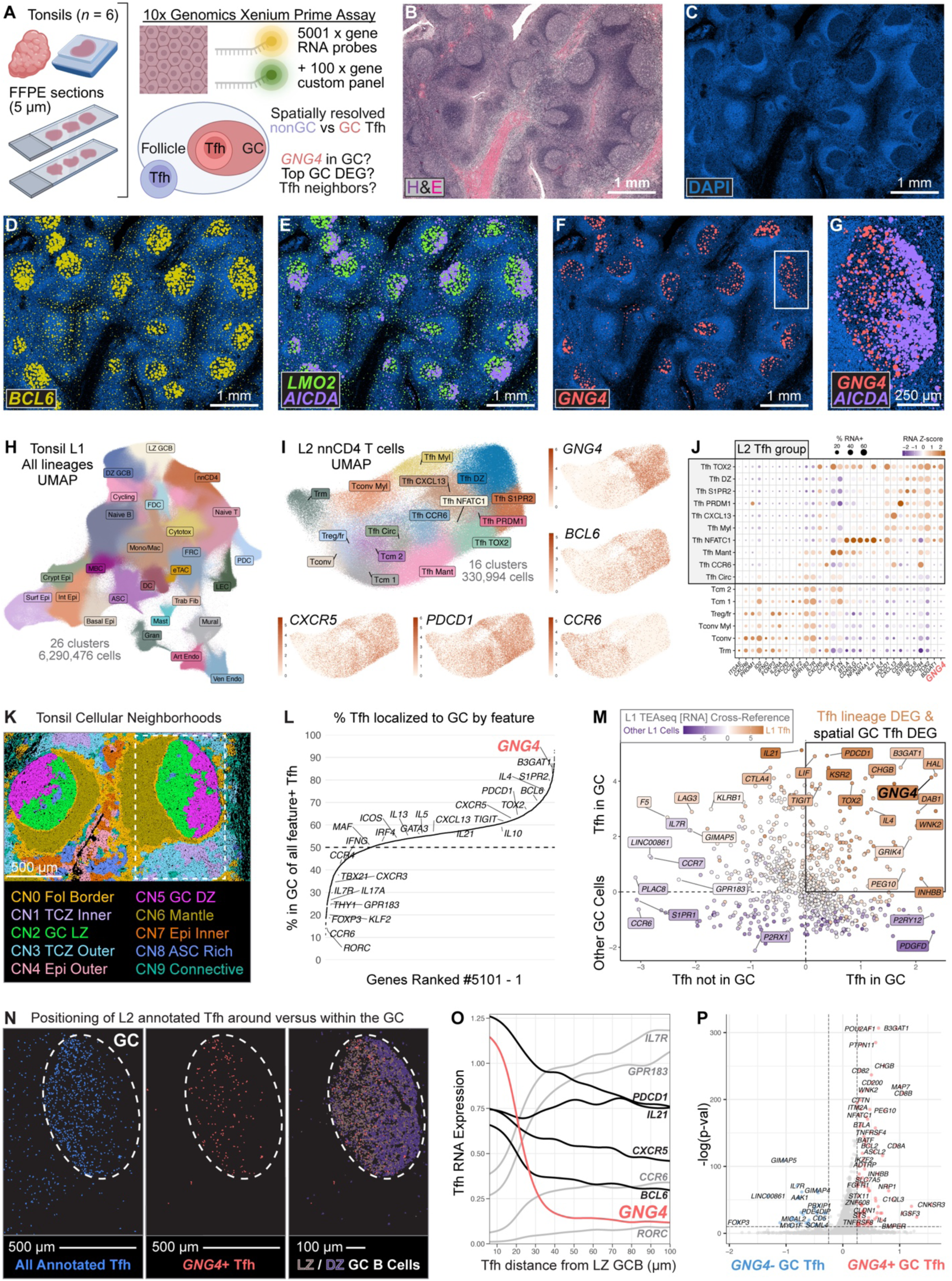
*GNG4* expression distinguishes activated Tfh positioned within the GC light zone. **(A)** Schematic of spatially-resolved tonsil RNA profiling using Xenium Prime (XP). **(B)** Representative images of sample TC653B showing H&E staining, (**C)** DAPI staining, and **(D-F)** DAPI with GC-associated transcripts overlaid. Box indicates **(G)** magnified follicle. **(H)** UMAP of L1 XP clusters (6,290,476 cells). **(I)** UMAP showing L2 subclusters of L1 nnCD4 T cells (*n* = 330,994). **(J)** Scaled average expression and percent expression of signature DEGs across nnCD4 T cell subclusters. **(K)** Representative images of sample TC653B showing cellular neighborhoods (CN) among all L1 cell types. **(L)** Rank plot showing percentage of Tfh localized to the merged GC CN by each feature. **(M)** Scatter plot of the log_2_FC for DEGs in XP L2 Tfh in the GC versus outside the GC (x-axis), XP L2 Tfh versus all other cells within the GC (y-axis), and TEAseq L1 Tfh vs all other clusters (color). Visualized genes were filtered for mutual XP and TEAseq coverage and *P* < 0.05 in each comparison using a Wilcoxon rank-sum test with Bonferroni correction. **(N)** Magnified image of panel K box with all L2 Tfh, *GNG4*^+^ Tfh subset, or L1 LZ/DZ GCB cells shown. **(O)** Kernel-smoothed line plot of L2 Tfh RNA expression versus distance from the nearest L1 LZ GCB cell. **(P)** Volcano plot of DEGs between *GNG4*^+^ vs *GNG4*^−^ Tfh within the merged GC CN (Wilcoxon rank-sum test *P* < 1e-10 & |log_2_FC| > 0.5).

To identify individual cells and determine cell identities within the spatial dataset, we performed cell segmentation followed by dimensionality reduction and clustering analysis to distinguish Tfh from other immune and stromal subsets. In our L1 annotation, we resolved 26 distinct clusters among ∼6.3 million segmented cells across all donors (Fig. 4H, fig. S14, data file S4). We manually annotated T cell, B cell, myeloid, epithelial, endothelial, and fibroblast-like clusters by assessing lineage-defining DEGs (Fig. 4H, fig. S14, data file S4) (*18, 42, 121–127*). To enhance resolution of Tfh from nonTfh, we subclustered L1 ‘nnCD4’ cells and annotated 16 distinct L2 nnCD4 T cell subsets (Fig. 4I, fig. S15, data file S4). Tfh clusters were enriched in *CXCR5*, *PDCD1* and *BCL6*, with decreased *PRDM1* and *KLF2* compared to nonTfh clusters (Fig. 4I-J, fig. S15, data file S4). *GNG4* was highly enriched in GC-like Tfh subclusters defined by increased *B3GAT1*, *TIGIT*, and *PDCD1* expression, consistent with our TEAseq data.

To resolve microanatomic compartments across all six tonsils for downstream quantitative analysis, we computed cellular neighborhoods (CN) as previously described (*128*). This analysis resolved 10 distinct CNs (Fig. 4K), which we annotated based on histologic location and cell type composition (Fig. 4B-C, fig. S16). The ‘GC LZ’ CN2 was enriched in FDC, LZ GCB, and several GC-like Tfh subclusters. Relative to CN2, ‘GC DZ’ CN5 was enriched in DZ Tfh and DZ GCB cells. Tfh clusters were spatially enriched within GC and follicular mantle regions, while nonTfh were found within the T cell zone and epithelial CNs (fig. S16). To define GC regions across all six samples, we first merged CNs annotated as LZ and DZ, then included any interspersed CNs enclosed within these merged GC CNs using ’hole-filling’ image processing techniques (Supplementary Methods). We then categorized Tfh as positive or negative for expression of each gene in the XP panel. Using merged GC CNs as a spatial anchor, we next determined the percentage of Tfh expressing each gene that were positioned within GCs (Fig. 4L, data file S5). Surprisingly, ∼30-50% of Tfh expressing GC-associated features such as *BCL6*, *TIGIT*, and *IL21* did not localize to GCs, indicating an incongruous relationship between transcription of GC-associated features and true, anatomic GC-localization of human Tfh. In contrast, 84% of *GNG4*^+^ Tfh were found within GCs, reflecting that *GNG4* has a high positive-predictive value (PPV) for the GC niche. Indeed, *GNG4* had the 25^th^ highest PPV for true GC positioning among 5,101 assessed genes (99.5^th^ percentile, Fig. 4L, data file S5), with greater fidelity than *BCL6, TOX2*, *S1PR2*, and many other conventionally GC-associated features.

However, several top features predicted to identify GC localization in Tfh by our Fig. 4L analysis appeared to result from contamination of other neighboring GC cell populations, including GC B cells (*SERPINA9, AICDA, LMO2*) (*120, 129*) and FDCs (*DSG2*, *SSTR2*, *SLC1A2*) (*124–126*), transcripts which were not enriched in Tfh from our TEAseq dataset. Indeed, it is known that single-cell analysis in XP and other spatial profiling assays can be affected by multiple sources of cell-to-cell contamination, including transcript diffusion, inaccurate segmentation, and vertical overlap of cells (*130*). To formally integrate the TEAseq and Xenium analyses and account for contaminating signals inherent to spatial profiling, we therefore plotted log fold change for (1) Tfh in the GC versus Tfh not in the GC and (2) Tfh versus all other cell lineages within the GC neighborhood, and points were colored by log fold change of Tfh versus all other cell lineages in our TEAseq RNA dataset. In this analysis, *GNG4* was among the most specific transcripts of GC Tfh (Fig. 4M, data file S5). In contrast, Th17 signature genes (*RORC, RORA*, *IL17A*) were highly enriched in Tfh outside GC (Fig. 4L-M), supporting our finding that Tfh17 primarily adopt Gγ4^−^ nonGC states in tonsil. Together, these data indicate that *GNG4* expression distinguishes tonsillar Tfh transcriptional states in GC from nonGC regions with greater anatomic fidelity than conventionally GC-associated features.

Indeed, *GNG4*^+^ Tfh were primarily positioned near LZ GC B cells (Fig. 4N), supporting our previous inference from the spatial patterning of *GNG4* transcripts (Fig. 4F-G). We hypothesized that proximity of Tfh to B cells in the LZ is associated with *GNG4* expression. Consistent with this hypothesis, *GNG4* expression steeply decreased in Tfh that were more distant from GC LZ B cells (Fig. 4O). In contrast, features conventionally used to define GC Tfh (*BCL6, CXCR5, PDCD1*) maintained greater expression with increasing distance from LZ GCB cells compared to *GNG4*. These data suggested that *GNG4* expression in Tfh was dependent upon spatial proximity to the LZ GCB cell niche, raising the possibility that *GNG4*^+^ Tfh represent a distinct subset with augmented capacity for B cell help.

We next sought to dissect whether *GNG4*^+^ Tfh adopt a transcriptionally distinct state relative to Tfh that lack *GNG4* expression within the GC niche. Indeed, *GNG4*^+^ GC Tfh were enriched in features of helper function (*CHGB, IL4*), cognate stimulation (*CD200*, *NRP1*, *BTLA*), and lineage commitment (*BCL6, TOX2, ASCL2*) compared to *GNG4*^−^ counterparts (Fig. 4P, data file S5) (*64, 131, 132*). In contrast, *GNG4*^−^ Tfh exhibited features consistent with GC egress (*GPR183, SELPLG*), Th17 polarity (*KLRB1*), and Tfr differentiation (*FOXP3, IL32, TNIP3*) (*9, 18, 72, 115, 133*). Given our previous finding that Gγ4 versus FOXP3 protein expression was anticorrelated in Tfh (fig. S11I), we hypothesized that *FOXP3* RNA^+^ Tfh in the GC may represent a precursor induced Tfr subset (*22*) with decreased *GNG4* expression. Indeed, *FOXP3* RNA^+^ GC Tfh were enriched in many signature features of Treg identity (*IL2RA, TNIP3, RTKN2, PBXIP1*) (fig. S18, data file S5) (*133–135*). In contrast, *FOXP3* RNA^−^ GC Tfh were primarily distinguished by increased *GNG4* expression, as well as features suggestive of helper function (*CHGB*, *IL21*, *CD40LG*) (*2, 132*). Together, these contrasting gene signatures indicate that *GNG4* expression is associated with a highly activated, GC-committed subset of Tfh and distinguishes stimulatory from regulatory subsets of the diverse follicular T cell pool in humans.

Our analyses leveraged pediatric tonsils as a model human SLO; however, lymph nodes (LN) are structurally distinct (*136*). To determine if *GNG4* defined GC Tfh in SLOs other than tonsil and from adult subjects, we reanalyzed adult tonsil Visium HD and adult LN Visium v1 data (fig. S19) (*137, 138*). In both datasets, *GNG4* expression was highly enriched in spatially and transcriptionally distinct GC regions (annotated ‘GC’ cluster in each tab of data file S6). In the tonsillar non-immune compartment, *GNG4* transcripts were also enriched in epithelial cells (fig. S13, S14, and S19). Together, these data suggest that *GNG4* expression marks GC-positioned Tfh across a diversity of ages and SLO tissues in humans.

### *GNG4* expression is associated with Tfh activation

Given the enrichment of *GNG4* transcript and enhanced accessibility in activated GC Tfh states (Figure 3E), we hypothesized that distinct *cis-*regulatory elements and TFs may underlie such tightly restricted transcription. To identify gene regulatory mechanisms at the *GNG4* locus in Tfh, we first compared chromatin accessibility between L3 3WNN Tfh-like clusters (Fig. 5A, data file S7). Three DAPs near the *GNG4* transcriptional start site contained promoter-like (PLS), proximal enhancer (pELS), and distal enhancer (dELS)-like sequences (fig. S20A-C) (*68, 139, 140*). Beyond the PLS, the dELS-containing DAP was particularly correlated with *GNG4* RNA expression (Fig. 5A, data file S7). GC-like Tfh exhibited greater accessibility of the dELS region as well as *GNG4* transcripts than the nonGC-like group, suggesting that this dELS region may augment *GNG4* expression in GC Tfh states. Notably, although the amino acid sequence of the Gγ4 protein in mice and humans are nearly identical, the 951 bp region encoding this distal enhancer is not conserved in the orthologous *Gng4* locus of mice (fig. S20C-D). Further, in mouse CD4 T cell scRNAseq data from multiple lymphoid organs, tissues, and health versus disease settings (*141–146*), *Gng4* RNA was either absent or only sparsely expressed (fig. S21, data file S8). Together, these data suggest that *Gng4* may not be a central feature of mouse GC Tfh programming and that an evolutionarily distinct enhancer element may account for this species difference in expression.

**Fig. 5.**
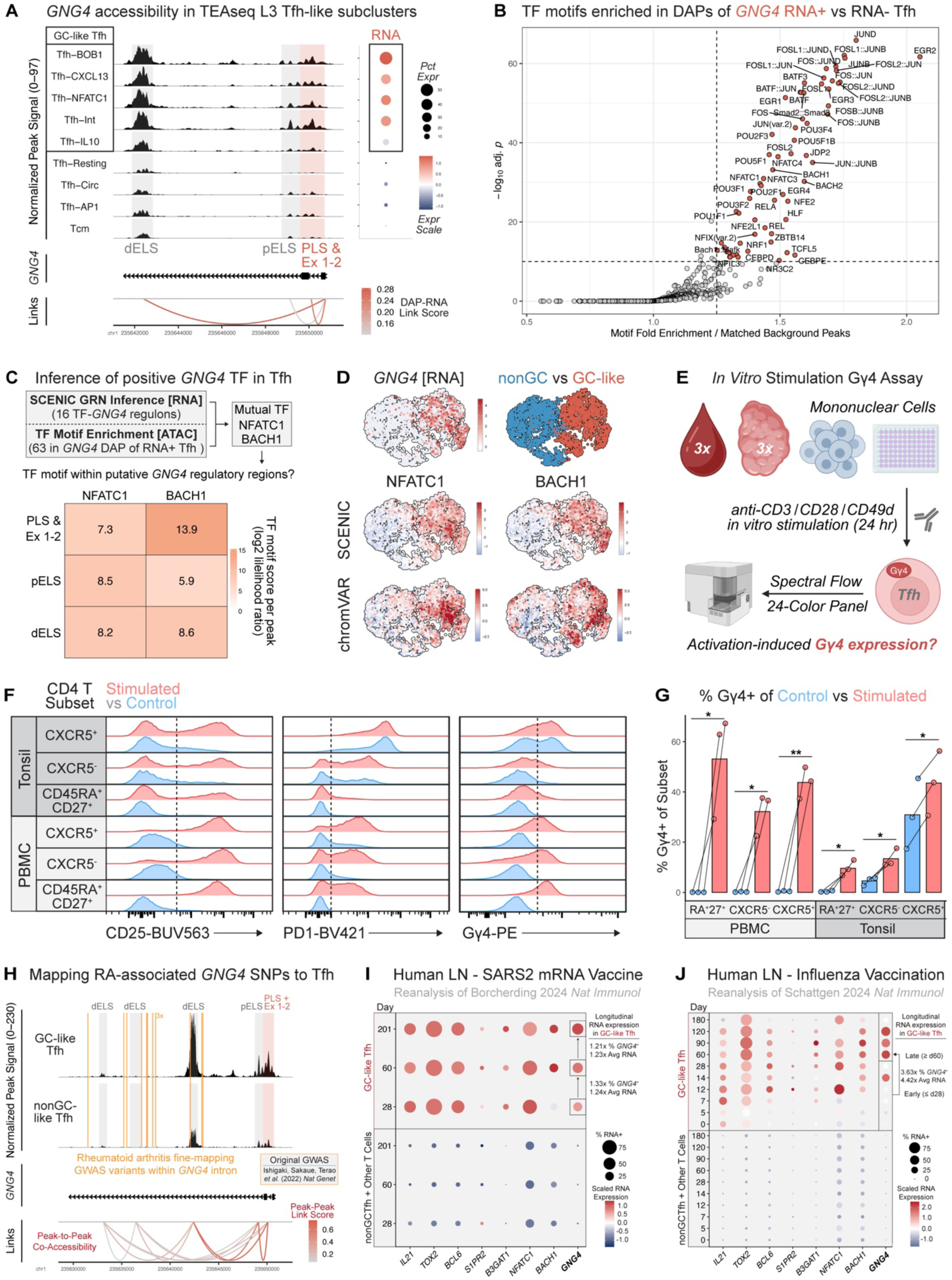
*GNG4* expression in Tfh is associated with activation *in vitro* and *in vivo*. **(A)** Normalized ATAC signal within *GNG4* subregion, pseudobulked per L3 3WNN subcluster. Differentially accessible peak (DAP, shaded) to *GNG4* RNA correlation is shown by links to promoter region. Dot plot shows percent and scaled RNA expression across clusters. DAPs containing proximal enhancer (pELS), distal enhancer (dELS) or promoter (PLS)-like sequences are annotated (*68*). Exons 1-2 (Ex 1-2) are within the PLS-containing DAP. **(B)** TF motifs enriched in DAPs of *GNG4* RNA^+^ versus RNA^−^ L3 3WNN Tfh (Tcm excluded, *p* < 1e-10 by hypergeometric test with Benjamini–Hochberg correction, fold-enrichment > 1.25). **(C)** Schematic illustrating multimodal inference of *GNG4*-associated TF activity. Grid shows log-likelihood of TF motif presence within dELS, pELS, and PLS/Ex1-2 DAPs. **(D)** 3WNN UMAPs of normalized *GNG4* RNA expression, nonGC vs GC-like group identity, and scaled SCENIC regulon AUC (*69*) or chromVAR *Z*-score (*70*) for predicted TFs. **(E)** Schematic of mononuclear cell *in vitro* stimulation experiment. **(F)** Representative histogram of CD25, PD1, and Gγ4 expression in Naive or nnCD4 CXCR5^−^ versus CXCR5^+^ CD4 T subsets paired between conditions. Gating scheme provided in fig. S22. **(G)** Barplot showing mean % Gγ4^+^ per subset between control and stimulated samples (paired *t*-test with two-stage step-up Benjamini, Krieger, and Yekutieli FDR correction for multiple comparisons, \**Q* < 0.05, \*\**Q* < 0.01). **(H)** Accessibility within *GNG4* subregion, pseudobulked by GC vs nonGC-like L3 3WNN Tfh group (Tcm excluded). Vertical lines (gold) indicate coordinates of fine-mapping variants from rheumatoid arthritis GWAS (*157*). Links (red) indicate co-accessibility of ATAC peaks computed by Cicero analysis (*170*). **(I)** Dot plots of percent and scaled RNA expression in human axillary lymph node (LN) CD4 T subsets during longitudinal SARS-CoV-2 (SARS2) BNT162b2 mRNA (*24*) or **(J)** inactivated quadrivalent influenza (Quad Flu) vaccination (*25*).

We next sought to identify transcription factor programs that are associated with the *GNG4^+^*Tfh state. Motif enrichment analysis of *GNG4* RNA^+^ versus RNA^−^ Tfh subsets (Fig. 5B, data file S7) identified 63 TFs, including members of the EGR, POU, and NFAT families that have established roles in Tfh differentiation and activation (*147–152*). These TF data further implicate *GNG4* in activated GC Tfh programming. To identify TFs that may directly regulate *GNG4* expression, we screened our SCENIC data for TFs with both (1) predicted binding motifs within the *GNG4* locus and (2) strongly correlated RNA expression with *GNG4* in Tfh (data file S7) (*69*). Both BACH1 and NFATC1 were identified by SCENIC and motif enrichment analyses (Fig. 5C, data file S7), indicating that these TFs may regulate the *GNG4*⁺ Tfh transcriptional state. Motifs for both BACH1 and NFATC1 were enriched within regions of the *GNG4* locus most accessible in GC-like Tfh, including the putative distal enhancer (Fig. 5C), further implicating these TFs in direct *GNG4* regulation. Moreover, GC-like Tfh exhibited both increased BACH1 and NFATC1 motif accessibility by chromVAR analysis, as well as regulon activity by SCENIC analysis (Fig. 5D, data file S7). Finally, *NFATC1* was among the top DEGs in the *GNG4*^+^ subset of spatially resolved GC Tfh (Fig. 4P), while *BACH1* was not included in our Xenium panel design. Thus, coordinated activity of NFATC1 and BACH1 may drive expression of *GNG4* in human GC Tfh.

NFATC1 is a central TCR-induced TF that is particularly essential for Tfh differentiation during acute viral infection and maintenance of Tfh function during chronic GC responses in mice (*151–153*). BACH1 can modulate sensitivity to ferroptosis, a cell death pathway in TCR-activated Tfh (*154–156*). Provided this context, we hypothesized that potent stimulation of Tfh within GCs may induce *GNG4* expression via NFATC1 and BACH1. We first asked whether Gγ4 protein expression can be induced in CD4 T cells by stimulating total mononuclear cells isolated from pediatric tonsil and adult blood donors (*n* = 3 each, unpaired, table S1) via CD3 ligation with CD28/CD49d costimulation (Fig. 5E, fig. S22). Gγ4 was induced across CD4 T cell subsets from PBMC and tonsil, along with activation associated proteins CD25 and PD1 (Fig. 5F-G). Furthermore, despite high levels of Gγ4 and activation features in GC-like Tfh at baseline, expression of both PD1 and Gγ4 were still moderately increased following stimulation. Together, these data indicated that T cell activation can induce *GNG4*/Gγ4 expression.

### *GNG4* risk alleles for autoimmune arthritis are strongly linked to variants associated with increased expression upon CD4 T cell activation

Given that *GNG4* expression distinguished activated Tfh positioned within GCs, we next asked whether genetic variation in *GNG4* might contribute to human diseases involving GC Tfh-driven pathology. A recent large, multi-ancestry genome-wide association study (GWAS) found that genetic variation at the *GNG4* locus is associated with rheumatoid arthritis (RA) susceptibility (*157*). Fine-mapping methods narrowed the association to 13 candidate causal variants, which were all non-coding and located within an intronic region of *GNG4*, including the lead variant rs61512163. However, it remains unclear whether these non-coding variants alter *GNG4* expression or function, the direction of their allelic effects, or the cell types that are impacted. To address these questions, we first mapped all 13 RA-associated variants to the *GNG4* locus across Tfh in our TEAseq data (Fig. 5H, fig. S23, data file S9). One single-nucleotide variant (rs6429213) resided within the dELS-containing DAP that exhibited maximal accessibility in GC-like Tfh (Fig. 5H). To test the hypothesis that genetically driven *GNG4* expression modulation may be associated with differential RA susceptibility, we reanalyzed data from a recent expression quantitative trait locus (eQTL) study—which relates genetic variation to gene expression levels—in human nnCD4 T cells before and after anti-CD3/CD28 stimulation (data file S9) (*158*). In this dataset, ten variants were significantly associated with altered *GNG4* expression after stimulation. However, these are unlikely to be causal variants themselves due to limited genotyping coverage of the eQTL study, but they may tag variants that are linked to the true causal allele(s), including candidates from the RA GWAS study (*159, 160*). Therefore, we performed linkage disequilibrium (LD) analysis—a measure of how often variants are inherited together—between these ten eQTLs and the lead rs61512163 variant identified by the fine-mapping GWAS (data file S9) (*157, 158*). Notably, alleles associated with higher *GNG4* expression in the eQTL study were tightly linked to the reported RA risk allele (maximum *R*^2^ = 0.94, *P* < 2.2e-16), suggesting that they likely tag a shared haplotype and that elevated *GNG4* expression may be one mechanism through which this haplotype increases RA susceptibility.

A separate expansive GWAS listed an additional *GNG4* variant that was associated with psoriatic arthritis (PsA) (*161*). In contrast to RA, the PsA-associated variant did not have strong linkage to the *GNG4* expression eQTL (maximum *R*^2^ < 0.01, data file S9), suggesting a different underlying causal variant or regulatory mechanism. Notably, haplotype blocks of the *GNG4* locus are relatively insulated from neighboring loci (fig. S23). Therefore, disease-associated variants found within the *GNG4* locus are less likely to be simply tagging causal variants at distal loci, increasing the likelihood that *GNG4* is in fact a causal gene in both RA and PsA studies (*162*). Together, these data indicate that increased *GNG4* expression in CD4 T cells may contribute to autoimmune arthritis pathogenesis in humans.

### *GNG4* expression is specifically increased in GC-like Tfh during vaccination

Our findings suggest that *GNG4* expression is highest in activated, GC-localized Tfh. Therefore, we hypothesized that GC-induction would increase the detection of *GNG4* expression in clinical contexts. To test this hypothesis, we reanalyzed scRNAseq data from human T cells in longitudinally sampled axillary LNs after vaccination with either SARS-CoV-2 (SARS2) BNT162b2 mRNA (Fig. 5I, fig. S24A-C) or inactivated quadrivalent influenza (IIV) (Fig. 5J, fig. S24D-K) (*24, 25*). *GNG4* expression increased in specifically GC-like Tfh clusters from d28 following BNT162b2 vaccination, with sparse expression in nonGC-like Tfh and nonTfh lineages at all timepoints (Fig. 5I, fig. S24C). *NFATC1* and *BACH1* expression was also restricted to GC-like Tfh, further suggesting that these TFs drive *GNG4* expression (Fig. 5A-D). Furthermore, *GNG4*, *NFATC1*, and *BACH1* expression were similarly restricted to GC-like Tfh throughout the IIV response (Fig. 5J, fig. S24D-K). *NFATC1* expression peaked in GC-like Tfh prior to *GNG4* (post-vaccination, d12 versus d14 and after), consistent with a driving role in initiating *GNG4* expression. Together these data demonstrate that vaccination induces *GNG4* expression specifically in Tfh, despite exposure of other CD4 and CD8 T cell subsets within axillary LNs to inflammatory stimuli. Further, these data suggest that *GNG4* expression may be maintained in Tfh during prolonged, vaccine-induced human GC responses.

## DISCUSSION

In this study, we leveraged multimodal approaches to define diverse Tfh cellular programs across peripheral blood and palatine tonsil compartments in humans. We resolved *GNG4/*Gγ4 as a central distinguishing feature of GC-like Tfh states in multimodal, single-cell studies. In spatial analyses, *GNG4* outperformed conventional features—including *BCL6*, *S1PR2*, and *TIGIT*—in identifying Tfh within anatomically defined GCs. We showed that *GNG4*^+^ Tfh positioned within GCs engage a distinct, spatially restricted gene expression program. As a direct application of these findings, a contemporaneous study of idiopathic multicentric Castleman disease—where GCs are present but atretic—found that *GNG4* was among the most dysregulated genes in both Tfh and dysplastic GCs of affected LNs (*163*) (Mumau *et al. bioRxiv*, 2025). These data suggest Gγ4 may mediate context-specific roles in Tfh across several human health and disease states.

The ability to identify GC-positioned Tfh in single-cell analyses based on Gγ4 expression enabled finer dissection of Tfh heterogeneity along multiple functional axes, including helper polarity and memory differentiation states. Tfh17 were less likely to express Gγ4 relative to other subsets, consistent with spatial data suggesting *RORC* expression was enriched outside of the GC. Further, Gγ4^−^ Tfh were more likely to display a central memory CD127^+^TIGIT^−^ phenotype, suggesting enhanced CM function (*13*). These data reflect recent findings in mice that Tfh17 are a CM-like subset with enhanced longevity and effector differentiation potential (*28*). In humans, cTfh17 exhibit superior survival characteristics and have been associated with chronic autoimmune conditions and durable vaccine memory (*6, 7, 164–166*). The role of Tfh17 cells in autoimmune and vaccine-induced memory responses is an ongoing area of investigation.

Resolving Tfh heterogeneity across tissues and gene regulatory states has been historically challenging. By applying TEAseq to SLOs, we found that clusters defined by chromatin accessibility were more disparate than those defined by RNA and protein-based data. These data are consistent with prior work suggesting that T cell identity defined by surface proteins often differs from subsets defined by ATAC or RNA profiles (*54*). This discordance between modalities suggests that separate, unimodal measures of gene expression may yield diverging inferences of B and T cell states that underlie GC biology. Further, features attributed to GC-like Tfh states by single-cell profiling frequently did not map to GC-positioned Tfh. Indeed, approximately 30-50% of Tfh expressing conventionally GC-associated features such as *BCL6*, *TOX2*, or *IL21* were positioned outside GCs, consistent with recent reports in mice (*35*). Together, our findings highlight the importance of using spatially resolved analyses to identify GC Tfh and dissect Tfh heterogeneity in tissues.

Though our data identify *GNG4* as a highly enriched feature of Tfh positioned within the GC, our data also suggest that Gγ4 expression is likely part of a common T cell activation program. Indeed, previous reports of *GNG4* expression in cancer-associated T cell subsets as well as agonist-selected PD1^+^ CD8⍺⍺ T cells in thymus also indicate an activation-induced mechanism (*43, 92, 99–107*). Further, stimulation of nnCD4 T cells from peripheral blood of two human donors for six days was found to induce Gγ4 protein expression by bulk mass spectrometry (*103*). Still, Gγ4 expression may be more tightly regulated in T cells than other features of activation, given (1) marked decrease in transcripts only microns from the GC light zone, (2) a leading meta-analysis rank in GC-like Tfh above common activation features, (3) restricted accessibility of putative *GNG4* CRE to GC-like Tfh states, and (4) the paucity of other nonTfh subsets expressing *GNG4* in the inflammatory environment of tonsils and vaccine-stimulated lymph nodes. Extending the multi-stage model of Tfh differentiation (*2, 64, 167*), we hypothesize that a high threshold of stimulation—one that Tfh may reach within the GC niche *in vivo*—is required for *GNG4* induction.

The specificity of *GNG4* expression to the GC Tfh compartment raises important questions about its potential biological functions. Studies in human cancers of neuronal origin—derived from tissues that physiologically express Gγ4—have linked Gγ4 overexpression with inhibition of CXCR4-driven chemotaxis (*168*), suggesting that similar mechanisms may enforce GC positioning in Tfh. Given that *GNG4*⁺ Tfh exhibit a highly activated, GC-resident state, Gγ4 may help restrict Tfh egress from the GC niche following potent TCR stimulation by cognate GC B cells, a hypothesis that warrants direct investigation. Notably, future mechanistic studies of Gγ4 function in Tfh may require human experimental systems, as Tfh in mice display sparse *Gng4* transcript. Further, the *GNG4* enhancer-like region identified in human Tfh is not conserved in mice. Given the breadth of Gγ4 induction after *in vitro* stimulation of human T cells, future study of this G protein subunit has the potential to identify new mechanisms in T cell activation and regulation that may be used for diagnostic or therapeutic benefit.

## MATERIALS AND METHODS

### Study design

This study aimed to delineate gene expression features of heterogeneous Tfh states across anatomic compartments. Tfh from tonsil versus PBMC donors (*n* = 4 each, unpaired) were first profiled by TEAseq and spectral flow cytometry. From these multimodal data, *GNG4* (encoding Gγ4) emerged as a significant feature of activated GC-like Tfh. Expression of Gγ4 versus other Tfh-associated proteins was assessed in a separate cohort of tonsil and PBMC donors by spectral flow cytometry (*n =* 4 each, unpaired). To assess the specificity of *GNG4* versus other genes for GC-localized Tfh, we profiled tonsils (*n* = 6) by single-cell spatial transcriptomics. Integrated epigenomic and transcriptional analyses were performed to infer gene regulatory features associated with *GNG4* expression in activated, GC-like Tfh states. Gγ4 induction upon T cell activation was evaluated in an additional cohort of tonsil and blood donors (*n* = 3 each, unpaired). Further, we assessed induction of *GNG4* in T cells post-vaccination by reanalyzing published scRNAseq data from longitudinally sampled human LNs. Finally, linkage disequilibrium analysis was performed to assess linkage between *GNG4* variants associated with autoimmune arthritides and increased *GNG4* RNA expression upon CD4 T cell activation.

### Tonsil mononuclear cell suspension processing

Whole tonsils were obtained as discarded surgical tissue from children undergoing tonsillectomy for sleep-related indications at CHOP (table S1). All samples were deidentified before receipt and designated as not human subjects research material by the CHOP IRB. Whole tonsils were washed in RPMI (Gibco, 11875093) supplemented with Penicillin-Streptomycin (PS, [Gibco, 15140122]) before preparing mononuclear cell (TMC) suspensions by mechanical disruption and gradient centrifugation (STEMCELL, 07861). TMC aliquots were cryopreserved until use.

### Tonsil processing for spatial transcriptomics

Discarded tonsillectomy tissue from patients with sleep-related indications was received and washed as above (table S1). 5 mm-thick tonsil samples were fixed in nuclease-free 4% PFA (EMS, 15174) and washed in PBS (Gibco, 10010023). FFPE blocks were prepared by the CHOP Pathology Core.

### PBMC processing

Healthy pediatric donors were recruited under CHOP IRB #15920 (table S1). Healthy adult donors were recruited by the Penn Human Immunology Core under IRB #705906 (table S1). Adult participant and pediatric guardian consent were obtained before study enrollment as well as pediatric subject assent where appropriate. Pediatric PBMC were isolated from heparinized venous blood by SepMate (STEMCELL, 85460) and Lymphoprep gradient centrifugation. Adult PBMC were isolated via leukapheresis and Lymphoprep. Samples were cryopreserved until use.

### Spectral flow cytometry

Samples were thawed, washed in Complete Medium (CM; RPMI Glutamax [Gibco, 61870036] supplemented with 10% FBS [Gemini Bio, 100500500], 1% L-Glutamine [Gibco, 25030081], and 1% PS), counted, and transferred to a 96-well plate. Cells were Fc-receptor blocked (FcR [BD, 564219]) and stained for viability (10 min, ice, Zombie NIR [BioLegend, 423106]), then quenched with FACS Buffer (FB, 2% FBS in PBS). Samples were incubated with chemokine receptor antibodies in Brilliant Stain Buffer (BSB; [BD, 566349]) for 20 minutes at 37°C, then washed twice with FB, before incubating with surface protein antibodies in BSB (15 min, RT), and washed a further three times with FB. Samples were then fixed and permeabilized (30 min, RT, [Invitrogen, 00552300]), washed in Perm/Wash buffer (PW, [Invitrogen, 00552300]), and stained with intracellular protein antibodies in PW (30 min, RT). Samples were washed three times with PW and once with FB, then acquired on an Aurora Spectral Flow Analyzer (Cytek, N700003). Viability controls were heated (5 min, 65°C), shocked (1 min, ice), and stained for viability as above. UltraComp eBeads Plus (Invitrogen, 01-3333-42) were used for single color unmixing controls, except for CD25-BUV563 where TMC were used. Flow cytometric data were gated in FlowJo (Treestar, v10.9.0) and exported to Prism (GraphPad, v10.2.2) and RStudio (Posit, 2024.04.0) for statistical analysis.

### *In vitro* cell stimulation

TMC and PBMC samples (table S1) were thawed and resuspended in CM. A TC-treated flat-bottom 96-well plate (Fisherbrand, FB012931) was coated with 50 μL anti-CD3 (Invitrogen, 16-0037-81) diluted to 2 μg/mL in PBS, then washed three times with PBS. 1e6 cells were stimulated per well in 200 μL CM containing 0.1 μg/mL each of anti-CD28/CD49d (BD, 347690). Matched unstimulated controls were prepared per sample by incubating 1e6 cells in 200 μL CM alone. Samples were incubated at 37°C with 5% CO_2_ for 24 h before flow staining.

### TEAseq

TMC and PBMC samples (table S1) were thawed into CM, then split for parallel flow cytometric and TEAseq processing (fig. S1). TEAseq samples were split for sorting all mononuclear versus CD4^+^ cells, yielding 16 samples for 8 donors. For the CD4-sorting TMC group, CD19^+^ cells were depleted using immunomagnetic beads (STEMCELL, 17854). Samples were then incubated in RPMI with FcR Block and Zombie-NIR viability stain. Samples were next labeled with CD15-PE, CD4-PECy7, CD4-ADT, and 16 different TotalSeqA hashtag-oligonucleotide antibodies (HTO; table S2 & S5). Unstained and single-color controls for CD4-PE, CD4-PECy7, and Zombie-NIR were prepared using TMC as above. Singlet, live, CD15^−^ cells and additionally CD4^+^ SSC^lo^ cells (fig. S1) were isolated by spectral FACS (Cytek, N700094). Equivalent proportions of sorted samples were pooled and counted to verify high viability. Cells were then aliquoted in Cell Staining Buffer (BioLegend, 420201) and processed following the TEAseq protocol (v4 with minor deviations detailed in Supplementary Materials) (*27, 169*).

### Spatial transcriptomics

Data were generated following 10x Genomics Xenium Prime protocols below. Six tonsil FFPE samples (table S1) were first sectioned at 5 μm thickness onto Xenium slides (CG000578_RevE) and maintained in a desiccator until use. To capture *GNG4* and additional transcripts of interest, we supplemented the Human Pan Tissue & Pathways Panel (10x Genomics, 1000671) with a 100-plex custom panel (table S9 [10x Genomics, 1000766]). After production, custom XP reagents were reconstituted (CG000760_RevB). Samples were deparaffinized, rehydrated, decrosslinked, and permeabilized (CG000580_RevE) then processed following the ‘In Situ Gene Expression with optional Cell Segmentation Staining’ protocol (CG000760_RevB). Data were collected using a Xenium Analyzer (CG000584_RevG). Following run completion, slides were stained with H&E (CG000613_RevB). Slide images were obtained using the Aperio VERSA 8 Scanning System (Leica).

### Public data reanalysis

Bioinformatic data reanalysis methods are detailed in the Supplementary Materials.

### Statistical analysis

For flow cytometric data, statistical significance was assessed by one-way ANOVA, unpaired *t*-test, paired *t*-test, and 95% confidence interval approaches, with corrections for multiple comparisons as appropriate. Statistical methods for TEAseq and Xenium Prime analyses are detailed in the Supplementary Materials. Relevant test details and thresholds for statistical significance are defined in the figure legends.

## Supplementary Materials

The PDF file includes:

Supplementary Text

Figs. S1 to S24

Tables S1 to S9

Legends for data files S1 to S10

References (*170–191*)

## Acknowledgments

The authors thank the patients and families who enabled this research. Further, we thank the CHOP Flow Cytometry Core Laboratory (RRID:SCR_009726) for cytometry assistance; the CHOP Pathology Core Laboratory (RRID:SCR_009729) for sample embedding and histology; the University of Pennsylvania Molecular Pathology and Imaging Core Laboratory (RRID: SCR_022420) for tissue sectioning; the CHOP High-Throughput Sequencing Core Laboratory for sequencing assistance; the University of Pennsylvania Human Immunology Core (RRID:SCR_022380) for processing of adult peripheral blood samples; and Samir Sayed of the Sarah E. Henrickson Laboratory for facilitating transfer of pediatric peripheral blood samples. We also thank Saori Sakaue for helpful insights and discussion of *GNG4* variant naming in our reanalysis of the RA fine-mapping GWAS data. Lastly, we would like to acknowledge all laboratory personnel and PhD thesis committee faculty members (Sarah E. Henrickson, David A. Hill, Chris A. Hunter, and Taku Kambayashi, advising SBD) for their suggestions during manuscript preparation.

## Author contributions

Conceptualization: SBD, LAV, DAO, AEB

Methodology: SBD, MG, AEB, DAO, JT, YQ, NDL, SR, EC, AC, BF

Software: SBD, YZ, TL, DAO, ICM

Formal analysis: SBD, YZ, DAO

Investigation: SBD, MG, AEB, DAO, KGK, NDL

Visualization: SBD, DAO

Resources: JT, AC, SR, EC, NR, SEH

Funding acquisition: LAV, DAO, NR, SEH

Project administration: LAV, DAO, AEB

Supervision: LAV, DAO, AEB

Writing – original draft: SBD, LAV, DAO, AEB, MG, YZ

Writing – review & editing: All authors

## Data and materials availability

TEAseq and Xenium Prime datasets are deposited to Gene Expression Omnibus and available through accession number GSE######. Tabulated data underlying the figures are provided in supplementary data files S1-10. All data needed to evaluate the conclusions in the paper are present in the paper or the Supplementary Materials. Related code for all analyses is provided on GitHub: https://github.com/theoldridgelab/TEAseqXeniumPrimeGNG4.

## Supplementary Materials

Please refer to the additional document containing Supplementary Text, Figures, and Tables.

## References and Notes

1. S. Crotty, T Follicular Helper Cell Differentiation, Function, and Roles in Disease. Immunity 41, 529–542 (2014).

2. S. Crotty, T Follicular Helper Cell Biology: A Decade of Discovery and Diseases. Immunity 50, 1132–1148 (2019).

3. C. G. Vinuesa, M. A. Linterman, D. Yu, I. C. M. Maclennan, Follicular Helper T Cells. Annual Review of Immunology 34, 335–368 (2016).

4. L. S. K. Walker, The link between circulating follicular helper T cells and autoimmunity. Nature Reviews Immunology 22, 567–575 (2022).

5. D. Yu, L. S. K. Walker, Z. Liu, M. A. Linterman, Z. Li, Targeting TFH cells in human diseases and vaccination: rationale and practice. Nature Immunology 23, 1157–1168 (2022).

6. H. Ueno, J. Banchereau, C. G. Vinuesa, Pathophysiology of T follicular helper cells in humans and mice. Nat Immunol 16, 142–152 (2015).

7. R. Morita et al., Human Blood CXCR5+CD4+ T Cells Are Counterparts of T Follicular Cells and Contain Specific Subsets that Differentially Support Antibody Secretion. Immunity 34, 108–121 (2011).

8. J. He et al., Circulating Precursor CCR7loPD-1hi CXCR5+ CD4+ T Cells Indicate Tfh Cell Activity and Promote Antibody Responses upon Antigen Reexposure. Immunity 39, 770–781 (2013).

9. W. Song, J. Craft, T Follicular Helper Cell Heterogeneity. Annual Review of Immunology 42, 127–152 (2024).

10. J. S. Hale, R. Ahmed, Memory T Follicular Helper CD4 T Cells. Frontiers in Immunology 6, (2015).

11. N. Schmitt, S.-E. Bentebibel, H. Ueno, Phenotype and functions of memory Tfh cells in human blood. Trends in Immunology 35, 436–442 (2014).

12. H. Feng et al., A novel memory-like Tfh cell subset is precursor to effector Tfh cells in recall immune responses. Journal of Experimental Medicine 221, (2024).

13. F. Zhu et al., Spatiotemporal resolution of germinal center Tfh cell differentiation and divergence from central memory CD4+ T cell fate. Nature Communications 14, (2023).

14. L. A. Vella et al., T follicular helper cells in human efferent lymph retain lymphoid characteristics. Journal of Clinical Investigation 129, 3185–3200 (2019).

15. L. Dalit et al., Divergent cytokine and transcriptional signatures control functional T follicular helper cell heterogeneity. Nature Immunology 26, 1821–1835 (2025).

16. H. Law et al., Human axillary lymph node T follicular helper (Tfh) and Precursor-Tfh cells exhibit functional flexibility following seasonal influenza vaccination. Clinical & Translational Immunology 14, (2025).

17. H. W. King et al., Integrated single-cell transcriptomics and epigenomics reveals strong germinal center–associated etiology of autoimmune risk loci. Science Immunology 6, (2021).

18. R. Massoni-Badosa et al., An atlas of cells in the human tonsil. Immunity 57, 379–399.e318 (2024).

19. J. H. Y. Siu et al., Early lymph node T follicular helper cell signalling hub drives influenza vaccine response in an ancestrally diverse cohort. eBioMedicine 122, (2025).

20. S. Sureshchandra et al., Deep profiling of human T cells defines compartmentalized clones and phenotypic trajectories across blood and tonsils. Immunity, (2025).

21. S. Kumar et al., Developmental bifurcation of human T follicular regulatory cells. Science Immunology 6, eabd8411 (2021).

22. C. Le Coz et al., Human T follicular helper clones seed the germinal center–resident regulatory pool. Science Immunology 8, (2023).

23. S. Fujioka et al., Single-cell multiomic analysis revealed the differentiation, localization, and heterogeneity of IL10+ Foxp3– follicular T cells in humans. International Immunology 37, 475–491 (2025).

24. N. Borcherding et al., CD4+ T cells exhibit distinct transcriptional phenotypes in the lymph nodes and blood following mRNA vaccination in humans. Nature Immunology 25, 1731–1741 (2024).

25. S. A. Schattgen et al., Influenza vaccination stimulates maturation of the human T follicular helper cell response. Nature Immunology 25, 1742–1753 (2024).

26. M. Durand et al., Human lymphoid organ cDC2 and macrophages play complementary roles in T follicular helper responses. J Exp Med 216, 1561–1581 (2019).

27. E. Swanson et al., Simultaneous trimodal single-cell measurement of transcripts, epitopes, and chromatin accessibility using TEA-seq. eLife 10, (2021).

28. X. Gao et al., T follicular helper 17 (Tfh17) cells are superior for immunological memory maintenance. eLife 12, e82217 (2023).

29. C. G. Vinuesa, M. C. Cook, Blood Relatives of Follicular Helper T Cells. Immunity 34, 10–12 (2011).

30. J. S. Hale et al., Distinct Memory CD4+ T Cells with Commitment to T Follicular Helper- and T Helper 1-Cell Lineages Are Generated after Acute Viral Infection. Immunity 38, 805–817 (2013).

31. M. Locci et al., Human Circulating PD-1+CXCR3−CXCR5+ Memory Tfh Cells Are Highly Functional and Correlate with Broadly Neutralizing HIV Antibody Responses. Immunity 39, 758–769 (2013).

32. K. C. Osum et al., A minority of Th1 and Tfh effector cells express survival genes shared by memory cell progeny that require IL-7 or TCR signaling to persist. Cell Reports 44, 115111 (2025).

33. N. Fazilleau et al., Lymphoid reservoirs of antigen-specific memory T helper cells. Nature Immunology 8, 753–761 (2007).

34. A. Asrir, M. Aloulou, M. Gador, C. Pérals, N. Fazilleau, Interconnected subsets of memory follicular helper T cells have different effector functions. Nat Commun 8, 847 (2017).

35. C.-H. Yeh, J. Finney, T. Okada, T. Kurosaki, G. Kelsoe, Primary germinal center-resident T follicular helper cells are a physiologically distinct subset of CXCR5hiPD-1hi T follicular helper cells. Immunity 55, 272–289.e277 (2022).

36. J. R. Kim, H. W. Lim, S. G. Kang, P. Hillsamer, C. H. Kim, Human CD57+ germinal center-T cells are the major helpers for GC-B cells and induce class switch recombination. BMC Immunology 6, 3 (2005).

37. K. Padhan et al., Acquisition of optimal TFH cell function is defined by specific molecular, positional, and TCR dynamic signatures. Proceedings of the National Academy of Sciences 118, e2016855118 (2021).

38. S. Horiuchi et al., Tox2 is required for the maintenance of GC T_FH_ cells and the generation of memory T_FH_ cells. Science Advances 7, (2021).

39. J. P. Weber et al., ICOS maintains the T follicular helper cell phenotype by down-regulating Krüppel-like factor 2. Journal of Experimental Medicine 212, 217–233 (2015).

40. C. H. Kim et al., Subspecialization of CXCR5+ T cells: B helper activity is focused in a germinal center-localized subset of CXCR5+ T cells. J Exp Med 193, 1373–1381 (2001).

41. Y. Hao et al., Integrated analysis of multimodal single-cell data. Cell 184, 3573–3587.e3529 (2021).

42. D. R. Glass et al., An Integrated Multi-omic Single-Cell Atlas of Human B Cell Identity. Immunity 53, 217–232.e215 (2020).

43. L. Billiet et al., Single-cell profiling identifies a novel human polyclonal unconventional T cell lineage. Journal of Experimental Medicine 220, (2023).

44. S. Ricciardi et al., The Translational Machinery of Human CD4+ T Cells Is Poised for Activation and Controls the Switch from Quiescence to Metabolic Remodeling. Cell Metabolism 28, 895–906.e895 (2018).

45. T. Wolf et al., Dynamics in protein translation sustaining T cell preparedness. Nature Immunology 21, 927–937 (2020).

46. M. Terekhova et al., Single-cell atlas of healthy human blood unveils age-related loss of NKG2C(+)GZMB(-)CD8(+) memory T cells and accumulation of type 2 memory T cells. Immunity 56, 2836–2854.e2839 (2023).

47. C. Schmidl, M. Delacher, J. Huehn, M. Feuerer, Epigenetic mechanisms regulating T-cell responses. J Allergy Clin Immunol 142, 728–743 (2018).

48. M. K. Atianand, K. A. Fitzgerald, Long non-coding RNAs and control of gene expression in the immune system. Trends Mol Med 20, 623–631 (2014).

49. C. A. Piccirillo, E. Bjur, I. Topisirovic, N. Sonenberg, O. Larsson, Translational control of immune responses: from transcripts to translatomes. Nat Immunol 15, 503–511 (2014).

50. A. Jay, C. M. Pondevida, G. Vahedi, The epigenetic landscape of fate decisions in T cells. Nat Immunol 26, 544–556 (2025).

51. P. S. Lim, J. Li, A. F. Holloway, S. Rao, Epigenetic regulation of inducible gene expression in the immune system. Immunology 139, 285–293 (2013).

52. A. L. Roy, Transcriptional Regulation in the Immune System: One Cell at a Time. Frontiers in Immunology 10, (2019).

53. J. R. Giles et al., Human epigenetic and transcriptional T cell differentiation atlas for identifying functional T cell-specific enhancers. Immunity 55, 557–574.e557 (2022).

54. Z. Thomson et al., Trimodal single-cell profiling reveals a novel pediatric CD8αα+ T cell subset and broad age-related molecular reprogramming across the T cell compartment. Nature Immunology 24, 1947–1959 (2023).

55. F. Pedregosa et al., Scikit-learn: Machine Learning in Python. Journal of Machine Learning Research, (2011).

56. N. Vinh, J. Epps, J. Bailey, Information theoretic measures for clusterings comparison: Is a correction for chance necessary? , (2009), pp. 135.

57. N. Vinh, J. Epps, J. Bailey, Information Theoretic Measures for Clusterings Comparison: Variants, Properties, Normalization and Correction for Chance. Journal of Machine Learning Research 11, 2837–2854 (2010).

58. E. Marina-Zárate et al., Highly functional and prolonged germinal center T follicular helper cell responses are associated with enhanced neutralizing antibody development. Immunity, (2025).

59. P. F. Cañete et al., Regulatory roles of IL-10–producing human follicular T cells. Journal of Experimental Medicine 216, 1843–1856 (2019).

60. M. Almanan et al., IL-10–producing Tfh cells accumulate with age and link inflammation with age-related immune suppression. Science Advances 6, eabb0806 (2020).

61. H. Li, C. D. Pauza, CD25^+^ Bcl6^low^ T follicular helper cells provide help to maturing B cells in germinal centers of human tonsil. European Journal of Immunology 45, 298–308 (2015).

62. G. Xin et al., Single-cell RNA sequencing unveils an IL-10-producing helper subset that sustains humoral immunity during persistent infection. Nature Communications 9, (2018).

63. M. M. Cavaco, P. Gaspar, R. do Amaral Vieira, F. Ribeiro, L. Graca, Heterogeneity of T follicular regulatory cells: exploring their expanding ontogeny and differentiation pathways. Immunol Cell Biol 103, 622–631 (2025).

64. Y.-J. Kim, J. Choi, Y. S. Choi, Transcriptional regulation of Tfh dynamics and the formation of immunological synapses. Experimental & Molecular Medicine 56, 1365–1372 (2024).

65. J.-Y. Lee et al., The Transcription Factor KLF2 Restrains CD4 + T Follicular Helper Cell Differentiation. Immunity 42, 252–264 (2015).

66. C. Su et al., Mapping effector genes at lupus GWAS loci using promoter Capture-C in follicular helper T cells. Nature Communications 11, (2020).

67. J. S. Weinstein et al., Global transcriptome analysis and enhancer landscape of human primary T follicular helper and T effector lymphocytes. Blood 124, 3719–3729 (2014).

68. ENCODE Project Consortium, An integrated encyclopedia of DNA elements in the human genome. Nature 489, 57–74 (2012).

69. S. Aibar et al., SCENIC: single-cell regulatory network inference and clustering. Nature Methods 14, 1083–1086 (2017).

70. A. N. Schep, B. Wu, J. D. Buenrostro, W. J. Greenleaf, chromVAR: inferring transcription-factor-associated accessibility from single-cell epigenomic data. Nature Methods 14, 975–978 (2017).

71. A. C. Olatunde, J. S. Hale, T. J. Lamb, Cytokine-skewed Tfh cells: functional consequences for B cell help. Trends in Immunology 42, 536–550 (2021).

72. L. Maggi et al., CD161 is a marker of all human IL-17-producing T-cell subsets and is induced by RORC. European Journal of Immunology 40, 2174–2181 (2010).

73. L. Cosmi et al., Human interleukin 17–producing cells originate from a CD161+CD4+ T cell precursor. The Journal of Experimental Medicine 205, 1903–1916 (2008).

74. M. A. Kleinschek et al., Circulating and gut-resident human Th17 cells express CD161 and promote intestinal inflammation. Journal of Experimental Medicine 206, 525–534 (2009).

75. F. Annunziato, S. Romagnani, Do studies in humans better depict Th17 cells? Blood 114, 2213–2219 (2009).

76. S. Q. Crome, A. Y. Wang, M. K. Levings, Translational Mini-Review Series on Th17 Cells: Function and regulation of human T helper 17 cells in health and disease. Clinical and Experimental Immunology 159, 109–119 (2009).

77. M. S. Maddur, P. Miossec, S. V. Kaveri, J. Bayry, Th17 cells: biology, pathogenesis of autoimmune and inflammatory diseases, and therapeutic strategies. The American Journal of Pathology 181, 8–18 (2012).

78. G. Castro et al., RORγt and RORα signature genes in human Th17 cells. PLoS One 12, e0181868 (2017).

79. T. Buchacher et al., PIM kinases regulate early human Th17 cell differentiation. Cell Rep 42, 113469 (2023).

80. F. Annunziato et al., Phenotypic and functional features of human Th17 cells. The Journal of Experimental Medicine 204, 1849–1861 (2007).

81. S. Burgler et al., Differentiation and functional analysis of human TH17 cells. Journal of Allergy and Clinical Immunology 123, 588–595.e587 (2009).

82. P. Daďová et al., A forskolin-mediated increase in cAMP promotes T helper cell differentiation into the Th1 and Th2 subsets rather than into the Th17 subset. Int Immunopharmacol 125, 111166 (2023).

83. E. Cano-Gamez et al., Single-cell transcriptomics identifies an effectorness gradient shaping the response of CD4+ T cells to cytokines. Nature Communications 11, (2020).

84. A. Capone et al., Systems analysis of human T helper17 cell differentiation uncovers distinct time-regulated transcriptional modules. iScience 24, 102492 (2021).

85. N. Manel, D. Unutmaz, D. R. Littman, The differentiation of human TH-17 cells requires transforming growth factor-β and induction of the nuclear receptor RORγt. Nature Immunology 9, 641–649 (2008).

86. N. J. Wilson et al., Development, cytokine profile and function of human interleukin 17–producing helper T cells. Nature Immunology 8, 950–957 (2007).

87. R. Wang et al., Genetic and pharmacological inhibition of the nuclear receptor RORα regulates TH17 driven inflammatory disorders. Nature Communications 12, 76 (2021).

88. A. S. Gukovskaya, G proteins in T cell signal transduction. Immunology Letters 31, 1–9 (1992).

89. C. E. Hörnquist et al., G(alpha)i2-deficient mice with colitis exhibit a local increase in memory CD4+ T cells and proinflammatory Th1-type cytokines. J Immunol 158, 1068–1077 (1997).

90. J. H. Kehrl, Heterotrimeric G Protein Signaling: Roles in Immune Function and Fine-Tuning by RGS Proteins. Immunity 8, 1–10 (1998).

91. R. M. Cinalli et al., T cell homeostasis requires G protein-coupled receptor-mediated access to trophic signals that promote growth and inhibit chemotaxis. European Journal of Immunology 35, 786–795 (2005).

92. E. A. Yost, T. R. Hynes, C. M. Hartle, B. J. Ott, C. H. Berlot, Inhibition of G-Protein βγ Signaling Enhances T Cell Receptor-Stimulated Interleukin 2 Transcription in CD4+ T Helper Cells. PLOS ONE 10, e0116575 (2015).

93. T. R. Hynes, E. A. Yost, C. M. Hartle, B. J. Ott, C. H. Berlot, Inhibition of G-Protein βγ Signaling Decreases Levels of Messenger RNAs Encoding Proinflammatory Cytokines in T Cell Receptor-Stimulated CD4(+) T Helper Cells. J Mol Signal 10, 1 (2015).

94. D. Wang, The essential role of G protein-coupled receptor (GPCR) signaling in regulating T cell immunity. Immunopharmacology and Immunotoxicology 40, 187–192 (2018).

95. M. J. Barnes, J. G. Cyster, Lysophosphatidylserine suppression of T-cell activation via GPR174 requires Gαs proteins. Immunology & Cell Biology 96, 439–445 (2018).

96. E. Lu, J. G. Cyster, G-protein coupled receptors and ligands that organize humoral immune responses. Immunological Reviews 289, 158–172 (2019).

97. D. S. Kuen et al., Critical regulation of follicular helper T cell differentiation and function by Gα(13) signaling. Proc Natl Acad Sci U S A 118, (2021).

98. H. Ham et al., Germline mutations in a G protein identify signaling cross-talk in T cells. Science 385, (2024).

99. L. Duan et al., G-Protein Subunit Gamma 4 as a Potential Biomarker for Predicting the Response of Chemotherapy and Immunotherapy in Bladder Cancer. Genes 13, 693 (2022).

100. A. Magen et al., Intratumoral dendritic cell–CD4+ T helper cell niches enable CD8+ T cell differentiation following PD-1 blockade in hepatocellular carcinoma. Nature Medicine 29, 1389–1399 (2023).

101. X. Lu, S. M. Lofgren, Y. Zhao, P. K. Mazur, Multiplexed transcriptomic profiling of the fate of human CAR T cells in vivo via genetic barcoding with shielded small nucleotides. Nature Biomedical Engineering 7, 1170–1187 (2023).

102. L. Zheng et al., Pan-cancer single-cell landscape of tumor-infiltrating T cells. Science 374, (2021).

103. M. L. Lawton et al., Multiomic profiling of chronically activated CD4+ T cells identifies drivers of exhaustion and metabolic reprogramming. PLOS Biology 22, e3002943 (2024).

104. B. Giotti et al., Single-Cell View of Tumor Microenvironment Gradients in Pleural Mesothelioma. Cancer Discovery 14, 2262–2278 (2024).

105. J.-E. Park et al., A cell atlas of human thymic development defines T cell repertoire formation. Science 367, eaay3224 (2020).

106. M. Heimli et al., Multimodal human thymic profiling reveals trajectories and cellular milieu for T agonist selection. Frontiers in Immunology 13, (2023).

107. Y. Li et al., Unraveling the spatial organization and development of human thymocytes through integration of spatial transcriptomics and single-cell multi-omics profiling. Nature Communications 15, (2024).

108. Y. J. Kim et al., CD5 Expression Dynamically Changes During the Differentiation of Human CD8^+^ T Cells Predicting Clinical Response to Immunotherapy. Immune Network 23, (2023).

109. A. Sood et al., CD5 levels define functionally heterogeneous populations of naïve human CD4^+^ T cells. European Journal of Immunology 51, 1365–1376 (2021).

110. E. M. Aandahl et al., CD7 Is a Differentiation Marker That Identifies Multiple CD8 T Cell Effector Subsets. The Journal of Immunology 170, 2349–2355 (2003).

111. S. Hyslop et al., CD7 regulates the persistence of terminally exhausted CD8+ T cells during chronic infection. Cell Reports 44, 116316 (2025).

112. V. Q. Van et al., CD47high Expression on CD4 Effectors Identifies Functional Long-Lived Memory T Cell Progenitors. The Journal of Immunology 188, 4249–4255 (2012).

113. S. Komori et al., CD47 promotes peripheral T cell survival by preventing dendritic cell–mediated T cell necroptosis. Proceedings of the National Academy of Sciences 120, (2023).

114. D. L. Hill et al., The adjuvant GLA-SE promotes human Tfh cell expansion and emergence of public TCRβ clonotypes. Journal of Experimental Medicine 216, 1857–1873 (2019).

115. D. Suan et al., T Follicular Helper Cells Have Distinct Modes of Migration and Molecular Signatures in Naive and Memory Immune Responses. Immunity 42, 704–718 (2015).

116. I. Sayin et al., Spatial distribution and function of T follicular regulatory cells in human lymph nodes. Journal of Experimental Medicine 215, 1531–1542 (2018).

117. M. A. Linterman et al., Foxp3+ follicular regulatory T cells control the germinal center response. Nature Medicine 17, 975–982 (2011).

118. Y. Chung et al., Follicular regulatory T cells expressing Foxp3 and Bcl-6 suppress germinal center reactions. Nature Medicine 17, 983–988 (2011).

119. J. T. Jacobsen et al., Expression of Foxp3 by T follicular helper cells in end-stage germinal centers. Science 373, eabe5146 (2021).

120. G. D. Victora et al., Identification of human germinal center light and dark zone cells and their relationship to human B-cell lymphomas. Blood 120, 2240–2248 (2012).

121. Y. Abe et al., A single-cell atlas of non-haematopoietic cells in human lymph nodes and lymphoma reveals a landscape of stromal remodelling. Nat Cell Biol 24, 565–578 (2022).

122. M. Xiang et al., A Single-Cell Transcriptional Roadmap of the Mouse and Human Lymph Node Lymphatic Vasculature. Front Cardiovasc Med 7, 52 (2020).

123. E. Trimm, K. Red-Horse, Vascular endothelial cell development and diversity. Nature Reviews Cardiology 20, 197–210 (2023).

124. B. A. Heesters et al., Characterization of human FDCs reveals regulation of T cells and antigen presentation to B cells. J Exp Med 218, (2021).

125. L. L. Tao, Y. H. Huang, Y. L. Chen, G. Y. Yu, W. H. Yin, SSTR2a Is a Useful Diagnostic Marker for Follicular Dendritic Cells and Their Related Tumors. Am J Surg Pathol 43, 374–381 (2019).

126. A. Aguzzi, N. J. Krautler, Characterizing follicular dendritic cells: A progress report. European Journal of Immunology 40, 2134–2138 (2010).

127. G. D. Victora, M. C. Nussenzweig, Germinal Centers. Annual Review of Immunology 40, 413–442 (2022).

128. C. M. Schürch et al., Coordinated Cellular Neighborhoods Orchestrate Antitumoral Immunity at the Colorectal Cancer Invasive Front. Cell 182, 1341–1359.e1319 (2020).

129. S. Montes-Moreno et al., Gcet1 (centerin), a highly restricted marker for a subset of germinal center-derived lymphomas. Blood 111, 351–358 (2008).

130. M. Bilous et al., From Transcripts to Cells: Dissecting Sensitivity, Signal Contamination, and Specificity in Xenium Spatial Transcriptomics. bioRxiv, (2025).

131. A. Renand et al., Neuropilin-1 Expression Characterizes T Follicular Helper (Tfh) Cells Activated during B Cell Differentiation in Human Secondary Lymphoid Organs. PLoS ONE 8, e85589 (2013).

132. I. Papa et al., TFH-derived dopamine accelerates productive synapses in germinal centres. Nature 547, 318–323 (2017).

133. F. Birzele et al., Next-generation insights into regulatory T cells: expression profiling and FoxP3 occupancy in Human. Nucleic Acids Research 39, 7946–7960 (2011).

134. Y. Benita et al., Gene enrichment profiles reveal T-cell development, differentiation, and lineage-specific transcription factors including ZBTB25 as a novel NF-AT repressor. Blood 115, 5376–5384 (2010).

135. R. Bhairavabhotla et al., Transcriptome profiling of human FoxP3+ regulatory T cells. Human Immunology 77, 201–213 (2016).

136. R. Pabst, Plasticity and heterogeneity of lymphoid organs. Immunology Letters 112, 1–8 (2007).

137. 10x Genomics, Inc., Visium HD Spatial Gene Expression Library, Human Tonsil (Fresh Frozen) Dataset. *10x Genomics Publicly Available Datasets*, (2024).

138. 10x Genomics, Inc., Human Lymph Node Visium Spatial Gene Expression Dataset *10x Genomics Publicly Available Datasets*, (2020).

139. F. Abascal et al., Expanded encyclopaedias of DNA elements in the human and mouse genomes. Nature 583, 699–710 (2020).

140. G. Perez et al., The UCSC Genome Browser database: 2025 update. Nucleic Acids Research 53, D1243–D1249 (2025).

141. M. Andreatta et al., Interpretation of T cell states from single-cell transcriptomics data using reference atlases. Nature Communications 12, (2021).

142. M. Andreatta et al., A CD4+ T cell reference map delineates subtype-specific adaptation during acute and chronic viral infections. eLife 11, (2022).

143. A. Cui et al., Dictionary of immune responses to cytokines at single-cell resolution. Nature 625, 377–384 (2024).

144. Y. Gu et al., Immune microniches shape intestinal Treg function. Nature 628, 854–862 (2024).

145. D. Zemmour et al., Single-cell gene expression reveals a landscape of regulatory T cell phenotypes shaped by the TCR. Nature Immunology 19, 291–301 (2018).

146. T. S. P. Heng et al., The Immunological Genome Project: networks of gene expression in immune cells. Nature Immunology 9, 1091–1094 (2008).

147. M. Yanagi et al., Bob1 maintains T follicular helper cells for long-term humoral immunity. Communications Biology 7, (2024).

148. D. Stauss et al., The transcriptional coactivator Bob1 promotes the development of follicular T helper cells via Bcl6. The EMBO Journal 35, 881–898 (2016).

149. X. Zhu et al., Optimal CXCR5 expression during Tfh maturation involves the Bhlhe40-Pou2af1 axis. Cell Reports 44, 116470 (2025).

150. A. Ogbe et al., Early Growth Response Genes 2 and 3 Regulate the Expression of Bcl6 and Differentiation of T Follicular Helper Cells. J Biol Chem 290, 20455–20465 (2015).

151. G. J. Martinez et al., Cutting Edge: NFAT Transcription Factors Promote the Generation of Follicular Helper T Cells in Response to Acute Viral Infection. The Journal of Immunology 196, 2015–2019 (2016).

152. A. Seth et al., AP-1–independent NFAT signaling maintains follicular T cell function in infection and autoimmunity. Journal of Experimental Medicine 220, (2023).

153. J.-U. Lee, L.-K. Kim, J.-M. Choi, Revisiting the Concept of Targeting NFAT to Control T Cell Immunity and Autoimmune Diseases. Frontiers in Immunology 9, (2018).

154. H. Nishizawa, M. Yamanaka, K. Igarashi, Ferroptosis: regulation by competition between NRF2 and BACH1 and propagation of the death signal. The FEBS Journal 290, 1688–1704 (2023).

155. D. Namgaladze, D. C. Fuhrmann, B. Brüne, Interplay of Nrf2 and BACH1 in inducing ferroportin expression and enhancing resistance of human macrophages towards ferroptosis. Cell Death Discovery 8, (2022).

156. Y. Yao et al., Selenium–GPX4 axis protects follicular helper T cells from ferroptosis. Nature Immunology 22, 1127–1139 (2021).

157. K. Ishigaki et al., Multi-ancestry genome-wide association analyses identify novel genetic mechanisms in rheumatoid arthritis. Nature Genetics 54, 1640–1651 (2022).

158. B. J. Schmiedel et al., Single-cell eQTL analysis of activated T cell subsets reveals activation and cell type–dependent effects of disease-risk variants. Science Immunology 7, (2022).

159. V. Tam et al., Benefits and limitations of genome-wide association studies. Nature Reviews Genetics 20, 467–484 (2019).

160. Z. Li, X. Zhou, Towards improved fine-mapping of candidate causal variants. Nature Reviews Genetics 26, 847–861 (2025).

161. A. Verma et al., Diversity and scale: Genetic architecture of 2068 traits in the VA Million Veteran Program. Science 385, (2024).

162. J. D. Wall, J. K. Pritchard, Haplotype blocks and linkage disequilibrium in the human genome. Nature Reviews Genetics 4, 587–597 (2003).

163. M. D. Mumau et al., Dysregulated lymphocyte localization in idiopathic multicentric Castleman disease. bioRxiv, (2025).

164. Y. Che et al., Circulating memory T follicular helper subsets, Tfh2 and Tfh17, participate in the pathogenesis of Guillain-Barré syndrome. Scientific Reports 6, 20963 (2016).

165. P. Zhou, et al., Longitudinal analysis of memory Tfh cells and antibody response following CoronaVac vaccination. JCI Insight 8, (2023).

166. Y. Chen et al., The Third dose of CoronVac vaccination induces broad and potent adaptive immune responses that recognize SARS-CoV-2 Delta and Omicron variants. Emerging Microbes & Infections 11, 1524–1536 (2022).

167. M. A. Podestà et al., Stepwise differentiation of follicular helper T cells reveals distinct developmental and functional states. Nature Communications 14, (2023).

168. J. Pal et al., Epigenetically silenced GNG4 inhibits SDF1α/CXCR4 signaling in mesenchymal glioblastoma. Genes & Cancer 7, 136–147 (2016).

169. E. Swanson, L. Graybuck, P. J. Skene, TEA-seq V.4 Protocol. dx.doi.org/10.17504/protocols.io.bwx17506pfre (2021).

170. H. A. Pliner et al., Cicero Predicts cis-Regulatory DNA Interactions from Single-Cell Chromatin Accessibility Data. Molecular Cell 71, 858–871.e858 (2018).

171. M. Stoeckius et al., Cell Hashing with barcoded antibodies enables multiplexing and doublet detection for single cell genomics. Genome Biology 19, (2018).

172. H. Heaton et al., Souporcell: robust clustering of single-cell RNA-seq data by genotype without reference genotypes. Nature Methods 17, 615–620 (2020).

173. A. Auton et al., A global reference for human genetic variation. Nature 526, 68–74 (2015).

174. M. D. Young, S. Behjati, SoupX removes ambient RNA contamination from droplet-based single-cell RNA sequencing data. GigaScience 9, (2020).

175. T. Stuart, A. Srivastava, S. Madad, C. A. Lareau, R. Satija, Single-cell chromatin state analysis with Signac. Nature Methods 18, 1333–1341 (2021).

176. X. Zhang et al., An immunophenotype-coupled transcriptomic atlas of human hematopoietic progenitors. Nature Immunology 25, 703–715 (2024).

177. K. Ferchen et al., A unified multimodal single-cell framework reveals a discrete state model of hematopoiesis in mice. Nature Immunology, (2025).

178. E. A. K. Depasquale et al., DoubletDecon: Deconvoluting Doublets from Single-Cell RNA-Sequencing Data. Cell Reports 29, 1718–1727.e1718 (2019).

179. P.-L. Germain, A. Lun, C. Garcia Meixide, W. Macnair, M. D. Robinson, Doublet identification in single-cell sequencing data using scDblFinder. F1000Research 10, 979 (2022).

180. L. Heumos et al., Best practices for single-cell analysis across modalities. Nature Reviews Genetics 24, 550–572 (2023).

181. H. M. Amemiya, A. Kundaje, A. P. Boyle, The ENCODE Blacklist: Identification of Problematic Regions of the Genome. Scientific Reports 9, (2019).

182. I. Tirosh et al., Dissecting the multicellular ecosystem of metastatic melanoma by single-cell RNA-seq. Science 352, 189–196 (2016).

183. M. D. Luecken et al., Benchmarking atlas-level data integration in single-cell genomics. Nature Methods 19, 41–50 (2022).

184. I. Korsunsky et al., Fast, sensitive and accurate integration of single-cell data with Harmony. Nature Methods 16, 1289–1296 (2019).

185. L. Waltman, N. J. Van Eck, A smart local moving algorithm for large-scale modularity-based community detection. The European Physical Journal B 86, (2013).

186. O. Fornes et al., JASPAR 2020: update of the open-access database of transcription factor binding profiles. Nucleic Acids Research 48, D87–D92 (2019).

187. S. B. Wells et al., Multimodal profiling reveals tissue-directed signatures of human immune cells altered with age. Nature Immunology 26, 1612–1625 (2025).

188. M. Andreatta, S. J. Carmona, UCell: Robust and scalable single-cell gene signature scoring. Computational and Structural Biotechnology Journal 19, 3796–3798 (2021).

189. J. Cao et al., The single-cell transcriptional landscape of mammalian organogenesis. Nature 566, 496–502 (2019).

190. C. J. Willer, Y. Li, G. R. Abecasis, METAL: fast and efficient meta-analysis of genomewide association scans. Bioinformatics 26, 2190–2191 (2010).

191. A. Bateman et al., UniProt: the Universal Protein Knowledgebase in 2025. Nucleic Acids Research 53, D609–D617 (2025).

