## Supplementary Materials (Text, Figures, and Tables) for "Multimodal analysis defines *GNG4* as a distinguishing feature of germinal center-positioned CD4 T follicular helper cells in humans"

Barnett Dubensky *et al.*

Corresponding authors:
Derek A. Oldridge, and Laura A. Vella,

**The PDF file includes:**
Supplementary Text
Figs. S1-S24
Tables S1-S9
Legends for data files S1-S10
References (*170 – 191*)

**Other Supplementary Material for this manuscript includes the following:**
Data files S1 to S10

### SUPPLEMENTARY TEXT

#### Supplementary Materials and Methods

##### TEAseq Data Generation

TEAseq – ADT incubation

To reduce batch effects in ADT labeling and downstream sequencing, all sixteen hashed samples were pooled before processing by the TEAseq protocol (*169*). Pooled cells were labeled using the TotalSeqA Human Universal Cocktail V1.0 (BioLegend, 399907) containing 155 distinct ADTs and 9 isotypes, supplemented with TotalSeqA CCR7-ADT (Table S2, BioLegend, 353247). The lyophilized Universal Cocktail was equilibrated at room temperature for 5 minutes, then centrifuged at 10,000 × *g* for 30 seconds. The Universal Cocktail was reconstituted with 26.62 µL of cell staining buffer. The CCR7-ADT was centrifuged at 14,000 × *g* for 10 minutes at 4°C before spiking 0.88 µL into the Universal Cocktail. The complete cocktail was vortexed and incubated for 5 minutes at room temperature, vortexed again, centrifuged at 10,000 × *g* for 30 seconds, and transferred to a 1.5 mL low-binding Eppendorf tube. The cocktail was then centrifuged at 14,000 × *g* for 10 minutes at 4°C. After centrifugation, the cocktail was kept on ice until used. The pooled cells were FcR-blocked again before ADT staining. 2.5 µL of Human TruStain FcX blocking reagent (BioLegend, 422304) was added to the pooled cells resuspended in 22.5 µL of cell staining buffer, pipette-mixed, and incubated for 10 minutes on ice. After 10 minutes, 25 µL of the complete ADT cocktail was added to the cells. The cells and antibody cocktail were pipette-mixed and incubated for 30 minutes on ice. After 30 minutes, cells were washed with 3 mL of cell staining buffer. This wash was repeated two additional times.

TEAseq – Cell permeabilization

The cells were centrifuged at 400 × *g* for 5 min at 4°C and the supernatant was discarded. The cell pellet was resuspended in 100 µL of chilled perm buffer (20 mM Tris-HCl pH 7.4, 150 mM NaCl, 3 mM $MgCl_2$, 0.01% Digitonin). Cells were incubated in the perm buffer for 5 minutes on ice. After incubation, cells were washed with 1 mL of chilled wash buffer (20 mM Tris-HCl pH 7.4, 150 mM NaCl, 3 mM $MgCl_2$). 10 µL was used to count the number of cells on the Invitrogen Countess 3 to determine resuspension volume. After the wash, cells were spun at 400 × *g* for 5 minutes at 4°C. The supernatant was discarded, and the cell pellet was resuspended in tagmentation buffer (20 mM Tris-HCl pH 7.4, 150 mM NaCl, 3 mM $MgCl_2$, 1 U/µL RNase Inhibitor [Millipore Sigma, 333539900]).

TEAseq – ATAC and RNA library preparation

10x Genomics RNA + ATAC Multiome: 3' RNA and ATAC libraries were generated using the Chromium Next GEM Single Cell Multiome ATAC and Gene Expression Reagent kit (10x Genomics, 1000283) following the 10x Genomics protocol CG000338 Rev F. The only deviation from the Multiome protocol was the addition of custom 0.2 µM ADT-Rev AMP and Additive HTO Primers (IDT) to the master mix of the Pre-Amplification step (step 4.1), as detailed in the TEAseq v4 protocol. Cells were loaded across 8 wells of a Chromium Next GEM Chip J (10x Genomics, 1000234). RNA libraries were indexed using the Dual Index Kit TT Set A (10x Genomics, 1000215) and amplified using an Eppendorf Mastercycler X50a protocol as follows:

Lid Temperature: 105°C
Reaction Volume: 100 µL
Step 1. 98°C for 45 seconds
Step 2. 98°C for 20 seconds
Step 3. 54°C for 30 seconds
Step 4. 72°C for 20 seconds
Step 5. Go to step 2 for a total of 14 cycles
Step 6. 72°C for 1 minute
Step 7. 4°C hold

ATAC libraries were uniquely indexed using the Single Index Kit N Set A (10x Genomics, 1000212) and incubated using the following protocol:

Lid Temperature: 105°C
Reaction Volume: 100 µL
Step 1. 98°C for 45 seconds
Step 2. 98°C for 20 seconds
Step 3. 67°C for 30 seconds
Step 4. 72°C for 20 seconds
Step 5. Go to step 2 for a total of 9 cycles
Step 6. 72°C for 1 minute
Step 7. 4°C hold

TEAseq – ADT and HTO library preparation

ADT and HTO libraries were generated from the normally discarded cDNA cleanup supernatant in step 6.2 of the 10x Genomics protocol CG000338 Rev F. 180 µL of SPRI beads (Beckman Coulter, B23317) were added to the supernatant and pipette-mixed 10 times. The supernatant and beads were incubated at room temperature for 10 minutes. Samples were then placed on a magnet to allow bead separation until the solution cleared. The beads were washed twice with 80% ethanol. DNA was then eluted from the beads with 90.5 µL of Buffer EB (Qiagen, 19086). Libraries were created with KAPA HiFi HotStart ReadyMix (KAPA, KM2602), SI-PCR Primer (IDT), and custom unique ADT-i7 and HTO-i7 primers (IDT, 100 nmol, DNA Oligo Purification, Standard Desalting, 65 Bases). The ADT libraries were uniquely indexed using 2.5 µL of 10 µM custom ADT-i7 primers and incubated using the following protocol:

Lid Temperature: 105°C
Reaction Volume: 100 µL

Step 1. 95°C for 3 minutes
Step 2. 95°C for 20 seconds
Step 3. 60°C for 30 seconds
Step 4. 72°C for 20 seconds
Step 5. Go to step 2 for a total of 15 cycles
Step 6. 72°C for 5 minutes
Step 7. 4°C hold

The HTO libraries were uniquely indexed using 2.5 µL of 10 µM custom HTO-i7 primers and incubated using the following protocol:

Lid Temperature: 105°C
Reaction Volume: 100 µL
Step 1. 95°C for 3 minutes
Step 2. 95°C for 20 seconds
Step 3. 64°C for 30 seconds
Step 4. 72°C for 20 seconds
Step 5. Go to step 2 for a total of 10 cycles
Step 6. 72°C for 5 minutes
Step 7. 4°C hold

After each ADT and HTO library PCR, a 1.6x bead:sample SPRI bead cleanup was performed.

<u>TEAseq – Library quality control</u>

Final libraries were quantified using Agilent's HSD5000 TapeStation kits (Agilent, 5067-5593 & 5067-5592). The RNA and ATAC libraries were diluted and normalized to 30 nM using the TapeStation results and further quantified via qPCR using the KAPA Library Quantification Kit for Illumina Platforms (KAPA Biosystems, 501965231). Final ADT and HTO libraries were diluted and normalized to 30 nM using the TapeStation results and further quantified using the Qubit dsDNA HS Assay kit (Invitrogen, Q32854). All libraries were then further diluted and normalized to 1.75 nM for sequencing.

<u>TEAseq – Library sequencing</u>

RNA, ADT, and HTO libraries were pooled at different volumes to target 30,000 reads, 12,000 reads, and 2,000 reads per recovered cell respectively. Pooled libraries were denatured and diluted following the Illumina NovaSeq 6000 Denature and Dilute Libraries Guide Protocol A for standard loading and sequenced on an Illumina NovaSeq 6000 using S2 100 and SP 100 reagent kits (Illumina, 20028316 & 20028401) with the following read lengths: Read 1: 28 bp, i7 Index: 10 bp, i5 Index: 10 bp, Read 2: 90 bp. Sequencing of ADT and HTO libraries was repeated to increase read depth using a SP 100 reagent kit with run read lengths of Read 1: 28 bp, i7 Index: 8 bp, Read 2: 15 bp. ATAC libraries were pooled at equal proportions to target 60,000 reads per recovered cell. Pooled ATAC libraries were denatured, diluted and sequenced on an

Illumina NovaSeq 6000 using a S4 200 reagent kit (Illumina, 20028313) with the following read lengths: Read 1: 50 bp i7, Index: 8 bp i5, Index: 24 bp, Read 2: 50 bp. All libraries were sequenced following Illumina Reverse Complement Workflow B using version 1.5 NovaSeq 6000 reagents. Original pooled sequencing of the RNA, ADT, and HTO libraries yielded 4.9e9 reads, repeated sequencing of the ADT and HTO libraries alone yielded 360e6 reads, and the ATAC library sequencing yielded 10.4e9 reads. ADT and HTO reads from the repeated higher-depth library sequencing run rather than original run were used for downstream data analysis.

**TEAseq Data Analysis**

Overview

Analysis of the TEAseq dataset featured twelve preprocessing steps, including quality control, multiplet filtering, and donor demultiplexing of pooled cells across eight GEM wells. After merging preprocessed single-cell data from each GEM well into a combined Seurat object (*41*), trimodal dimensionality reduction and clustering was performed at three levels of resolution – all mononuclear cells (Level 1, L1), T cells (L2), and Tfh-like cells (L3) – with further downstream analyses and data visualization. Key details for each step are summarized below. Related scripts are provided on GitHub: https://github.com/theoldridgelab/TEAseqXeniumPrimeGNG4.

Computing Environment

TEAseq analysis was performed within the Children's Hospital of Philadelphia (CHOP) high-performance computing cluster, primarily using the RStudio (2024.04.0) integrated development environment for R (4.4.0). Select analyses were performed using the HPC Linux command line interface via the Simple Linux Utility for Resource Management (SLURM) workload manager. Packages used throughout our analysis pipeline are listed in table S7. For reproducibility, '26' was specified as the default seed for random number generation in all R-based analyses.

Preprocessing Step 1 – Demultiplexing Illumina raw sequencing reads

As detailed above, ATAC, RNA, and ADT/HTO libraries across multiplexed samples of each GEM well were sequenced on separate Illumina flow cells. Raw BCL reads were demultiplexed into FASTQ files grouped by GEM well using bcl2fastq2 (2.20.0) with library-specific settings guided by Step 88 of the TEAseq v4 protocol (*169*), summarized below:

For ATAC S4 flow cell libraries, four unique 8 bp index sequences were used to demultiplex BCL reads per GEM (table S3) following the Illumina forward strand workflow. Reads were defined as Read 1, Index 1 (i7), Index 2 (i5), or Read 2 using base masks Y50n*, I8n*, n8Y16, and Y50n* respectively (--use-bases-mask). A maximum of one mismatch between detected index reads and specified index sequences was permitted (--barcode-mismatches). Reads containing one of four unique 8 bp index sequences per GEM well library were assigned a suffix 'a' through 'd' and concatenated for downstream analysis.

RNA libraries from the S2 and SP flow cells were demultiplexed per GEM using Dual Index Kit TT Set A sequences (table S3) following the Illumina reverse complement workflow. Reads were defined as Read 1, Index 1 (i7), Index 2 (i5), or Read 2 using base masks Y*, I10, Y10, and Y* respectively (--use-bases-mask). A maximum of one mismatch between detected index reads and specified index sequences was permitted (--barcode-mismatches).

ADT/HTO libraries from the repeated SP flow cell sequencing run were demultiplexed per GEM using custom i7 index sequences (table S3) following the Illumina reverse complement workflow. Reads were defined as Read 1, Index 1 (i7) or Read 2 using base masks Y28n*, I8n*, and Y15n* respectively (--use-bases-mask). Short adapter reads were masked at a threshold of 8 bp (--mask-short-adapter-reads), and any reads shorter than 8 bp were discarded (--minimum-trimmed-read-length). A maximum of one mismatch between detected index reads and specified index sequences was permitted (--barcode-mismatches).

Preprocessing Step 2 – Preparation of ATAC and RNA cell-by-feature count matrices

To obtain cell-by-peak (ATAC) and cell-by-gene (RNA) count matrices with joint cell-barcodes, the cellranger-arc (2.0.2) count function was executed for all eight GEM wells separately. Each of the four indexed 'a' through 'd' ATAC libraries as well as both RNA libraries from the SP and S2 flow cells were provided as input FASTQ files. For read alignment, a contemporaneous Genome Reference Consortium Human Build 38 (hg38) reference genome file supplied by 10x Genomics was used (refdata-cellranger-arc-GRCh38-2020-A-2.0.0).

Preprocessing Step 3 – Preparation of ADT and HTO cell-by-feature count matrices

To obtain cell-by-ADT and cell-by-HTO count matrices per GEM well, the cellranger (8.0.0) multi function was used. RNA libraries from both SP and S2 flow cells were provided as input FASTQ files, as well as ADT and HTO libraries from the repeated SP flow cell sequencing. RNA libraries were only provided as required to execute cellranger multi. Output RNA data were not used in place of the Step 2 cellranger-arc RNA count matrix. For the 'gene-expression' partition of the input configuration spreadsheet, assay chemistry was specified as ARC-v1, and the same hg38 reference genome file as used in Step 2 was provided. For configuration of the 'feature' partition, barcodes for all 16 HTO as well as the ADT cocktail (table S2) were provided. To enhance feature name legibility, 'Hu.' and 'HuMs.' prefixes were removed from the antibody reference and UMI counting CSV file provided by BioLegend, in addition to other minor feature name changes (e.g. punctuation, symbols, and common protein name usage).

Preprocessing Step 4 – SNP-based donor demultiplexing

As an orthogonal approach to HTO-based donor demultiplexing and multiplet detection (*171*), the single-nucleotide polymorphism (SNP) profiles of all eight donors were analyzed by souporcell (2.5) to infer donor of origin per cell (*172*). The core souporcell pipeline (souporcell_pipeline.py) was run per GEM well using the provided singularity (3.11.1-1.e19) container within the CHOP HPC Linux command line interface. The genome.fa FASTA file from the same hg38 reference genome as used in Steps 2-3 was provided for alignment (--f). Common variants filtered to >=2% allele frequency and limited to SNPs for GRCh38 from the 1000 Genomes Project were used for variant calling as recommended (--common_variants) (*173*). Given that four peripheral blood donors and four additional tonsil donors were analyzed, the souporcell pipeline was configured to resolve eight SNP-based clusters (--k). To reduce reference bias and false-positive variant calling, RNA reads were remapped using minimap2 as recommended. As each donor sample was split and hashed for sorting all mononuclear cells as well as $CD4^+$ cells (figure S1), two enriched HTO were expected per donor. In conjunction with HTO-based demultiplexing data (Step 7), these SNP data enhanced downstream multiplet filtering (Step 8) and donor annotation (Step 9).

Preprocessing Step 5 – Assembly of multimodal Seurat objects per GEM well

Following these initial preprocessing steps, Seurat (5.1.0) objects were created per GEM well including data from the ATAC, RNA, ADT, HTO, and SNP modalities (*41*). To mitigate potentially confounding ambient RNA signal, the cellranger-arc filtered cell-by-gene count matrix was corrected using SoupX (1.6.2) as follows (*174*): First, the ambient expression profile of empty droplets as well as the contamination fraction were estimated using cellranger-arc output cell clustering and barcode calling information (autoEstCont). Next, this ambient profile and contamination fraction were used to correct RNA counts per cell. This modified cell-by-RNA count matrix was then used to create the Seurat object to which other assays were appended (CreateSeuratObject). For the ATAC modality, Signac (1.13.0) was used to create a ChromatinAssay from the cellranger-arc output filtered cell-by-peak count matrix and fragments file (*175*). For ADT and HTO modalities, the intersection of barcodes from the unfiltered cellranger multi and filtered cellranger-arc cell-by-feature matrices was found. Next, this common cell barcode list was used to filter the cellranger multi 'Antibody Capture' count matrix. The barcode-filtered ADT and HTO data were then added to the Seurat object as separate assays. Finally, numerical SNP-based cluster assignments (0-7, corresponding to eight unique donors) from Step 4 were added as metadata columns to the multimodal Seurat object.

Preprocessing Step 6 – Initial quality control filtering

As a first-pass quality control (QC) filter, a lower-bound count threshold across modalities was applied to remove low-quality or empty droplets possibly missed by the cellranger-arc barcode calling algorithm (Step 2). We reasoned that first removing such empty droplets would increase the accuracy of downstream preprocessing steps, such as distinguishing multiplet from singlet true cells (Step 8). Further, an upper-bound mitochondrial mRNA enrichment filter (< 30%) was applied to remove low-quality and dying cells that may confound multiplet calling as well as other downstream analyses. Cells with low HTO counts were removed to improve the accuracy of HTO-based donor demultiplexing (Step 7). No filter was applied to the number of unique ADT per cell (nFeature_ADT), as the ADT cocktail (table S2) may be biased towards particular immune cell types, with low coverage of markers for rare subsets. Relatively permissive thresholds were used in this first-pass QC filter of empty droplets before multiplet detection, with a more stringent second-pass QC filter applied across modalities later at Step 10. Specific first-pass QC filters were as follows:

nCount_RNA > 350 (total RNA UMIs)
nFeature_RNA > 250 (total number of unique RNA transcript species detected)
nCount_ATAC > 700 (total number of ATAC peak-overlapping chromatin fragments)
nFeature_ATAC > 350 (unique peaks detected containing at least one fragment count)
nCount_ADT > 175 (total ADT UMIs)
nCount_HTO > 50 (total HTO UMIs)
percent.mt < 30 (percentage of all mRNA UMIs detected with *MT*- gene name prefix)

Preprocessing Step 7 – Determining HTO signal purity per cell

Given that SNP-based demultiplexing could only discern donors (8 total), but not 'bulk' and 'CD4' sorted samples per donor (16 total, figure S1), HTO-based demultiplexing was performed as well. To determine the dominant HTO signal per cell, a manual count percentage approach similar to other highly multiplexed single-cell profiling experiments was employed (*176-178*): First, the percentage of total HTO counts deriving from each HTO per cell was computed as a measure of HTO signal purity. Next, each HTO was ranked per cell by decreasing count percentage, with the top ranked HTO indicating the corresponding original FACS sample. HTO-based sample annotations ('hto.sort') were further grouped by donor ('hto.donor') as well as tissue type ('hto.tissue'), then stored as metadata columns within each GEM well Seurat object. Finally, the percentage of HTO counts deriving from the bottom 15 ranked HTO was computed, wherein a high percentage is suggestive of multiplet cells or poor HTO data quality. HTO count percentage thresholds for singlet versus multiplet and low-quality cells were specified in the following step.

Preprocessing Step 8 – Singlet and multiplet filtering

A 10% multiplet rate was anticipated based on estimates provided by the 10x Genomics Multiome protocol used for TEAseq (CG000338 Rev F, August 26, 2022). In devising our multiplet filtering approach, we anticipated that our TEAseq data may contain both homotypic and heterotypic doublets and multiplets among eight unique donors, sixteen hashed samples, and many distinct cell types (figure S1). Further, we considered that some approaches may only resolve doublets and multiplets between different samples (HTO) or donors (SNP). In contrast, computational approaches leveraging the RNA or ATAC profiles of single-cells such as scDblFinder (*179*) may identify cell type doublets and multiplets from the same sample or donor, though identifying homotypic cell type doublets and multiplets remains challenging. However, we reasoned that the latter may in part be identified by outlier high UMI counts across TEAseq modalities. To enhance removal of homotypic and heterotypic doublets and multiplets from our analysis, we therefore applied five distinct filters to all GEM wells, leveraging each modality of our TEAseq data: HTO, SNP, ATAC, RNA, and an upper UMI threshold across modalities.

For HTO-based filtering, a manual count percentage threshold was used following other highly multiplexed single-cell profiling studies (*176-178*). Cells with a majority of HTO counts from the top rank HTO (> 50%, determined in Step 7) and a low percentage from the second rank HTO (< 15%) were considered singlets. For SNP-based filtering, all cells annotated as doublets by the Souporcell algorithm (Step 4) were removed. scDblFinder (*179*) was used for both ATAC and RNA-based doublet detection, given that some cell types may be readily distinguished by epigenomic but not transcriptional differences, and vice versa. For both modalities, an estimated doublet rate of 10% (dbr = 0.1) was used. As recommended for the ATAC modality (*180*), correlated peaks were aggregated into 25 meta features and normalized before doublet prediction (aggregateFeatures = TRUE, nfeatures = 25, processing = 'normFeatures'). After filtering barcodes classified as non-singlet by any of these four filters, an upper UMI threshold across modalities was applied, as well as an upper unique feature count threshold for ATAC and RNA:

hto.class = 'singlet' (#1 ranked HTO > 50%, #2 HTO < 15%, #2-16 HTO < 50%)
snp.class = 'singlet' (Souporcell, RNA modality) (*172*)
scDblF.RNA.class = 'singlet' (scDblFinder, RNA modality) (*179*)
scDblF.ATAC.class = 'singlet' (scDblFinder, ATAC modality) (*179*)
nCount_HTO < 4000
nCount_ADT < 4500
nCount_RNA < 20000
nFeature_RNA < 5000
nCount_ATAC < 50000
nFeature_ATAC < 15000

Preprocessing Step 9 – Donor annotation from SNP clustering and HTO count percentage data

After removing multiplets, donor and sample labels were assigned per cell using SNP clustering (Step 4) and HTO ranking (Step 7) data. First, HTO counts were normalized per cell by centered log-ratio (CLR) transformation using the NormalizeData function (normalization.method = 'CLR', margin = 2). Next, the normalized HTO signal distribution was inspected across SNP clusters. Given that each donor cell pellet was split into two samples for hashing and sorting (figure S1), each SNP cluster was annotated based on enrichment of the two corresponding HTO. As Souporcell was run on each GEM independently, the specific HTO pairing across the eight numerical SNP clusters varied. HTO-guided SNP annotations for each donor (snp.donor) and tonsil versus peripheral blood tissue type (snp.tissue) were stored as Seurat object metadata.

<u>Preprocessing Step 10 – Second-pass quality control</u>

Before merging data from each GEM well and determining a common ATAC peak set (Step 11), a more stringent second-pass QC filter was applied across modalities. Beyond standard filters for UMI and unique feature counts, the mitochondrial transcript percentage threshold was lowered to 25%, and several additional ATAC-specific filters were applied:

percent.mt < 25 (percentage of all mRNA UMI detected with *MT*- gene name prefix)
nCount_RNA > 600
nCount_RNA < 12500
nFeature_RNA > 450
nFeature_RNA < 4000
nCount_ADT > 175
nCount_ADT < 2500
nCount_HTO > 50
nCount_HTO < 3000
nCount_ATAC < 40000
nCount_ATAC > 700
nFeature_ATAC < 15000
nFeature_ATAC > 350
frac_reads_in_peaks > 0.5 (fraction of ATAC reads in peak regions)
TSS.enrichment > 4 (transcription start site enrichment score defined by ENCODE (*68*), from TSSEnrichment function of Signac (*175*) for ATAC data)
nucleosome_signal < 0.7 (strength of nucleosomal banding pattern, from NucleosomeSignal function of Signac for ATAC data)
blacklist_fraction < 0.005 (ratio of ATAC reads in genomic blacklist regions associated with artefactual signal, provided by ENCODE) (*181*)
atac_mitochondrial_reads < 5000 (number of mitochondrial ATAC fragments)

<u>Preprocessing Step 11 – Unification of overlapping ATAC peak loci and GEM merging</u>

Given that the cellranger-arc pipeline was run on each GEM well separately, the exact coordinates of ATAC peak regions called for each GEM well varied. Thus, a common ATAC peak set across GEM well objects was determined using the UnifyPeaks function of Signac (*175*), merging overlapping peaks (mode = reduce, default). Next, the cell-by-peak matrix for each GEM well object was recomputed by providing the Signac FeatureMatrix function with this common peak set and the original ATAC fragment coordinates of each object. Finally, the merge function was used to combine all eight GEM well objects into a single object, with GEM well number identifiers appended to each cell barcode (‘add.cell.ids’) correspondingly.

<u>Preprocessing Step 12 – Scoring cell-cycle transcriptional signatures</u>

To assess whether transcriptional signatures of cycling cells drive downstream dimensionality reduction and clustering results, the Seurat CellCycleScoring workflow (*41*) was used as follows:

First, RNA count layers across GEM wells were joined. Next, RNA counts were log-normalized using the NormalizeData function. Transcriptional signatures associated with S and G2/M phases of the cell-cycle were extracted from the 'cc.genes.updated.2019' object provided by Seurat (*41, 182*). Using these signatures, the CellCycleScoring function was applied, yielding a 'S.Score' and 'G2M.Score' per cell, as well as an ordinal 'Phase' annotation of G1, G2M, or S (renamed as 'CC.Phase' for clarity). To enable regression of differences among cycling cells while preserving transcriptional differences between cycling and non-cycling states, the difference in S.Score and G2M.Score ('CC.Difference') per cell was computed as recommended. Ultimately, no regression of cycling signatures or other variables was performed during feature scaling for any downstream L1-L3 analyses, though these variables are stored as metadata columns in our provided Seurat objects and available for reanalysis.

<u>Feature normalization, variable feature selection, and scaling</u>

ATAC, RNA, and ADT data were processed for dimensionality reduction and integration following relevant guidelines from the developers of Signac (ATAC analysis) and Seurat (RNA and ADT analyses). Identical workflows were used across L1-L3 objects per modality, with relevant functions and parameters detailed below:

For the ATAC modality, the cell-by-peak count matrix was subjected to term frequency inverse document frequency (TF-IDF) normalization using the RunTFIDF function. To include both rare and common peaks in downstream steps, features for dimensionality reduction were selected using the FindTopFeatures function with 'min.cutoff' set to 'q0'.

For RNA, joined count layers from cell-cycle scoring (Preprocessing Step 12) were retained. Next, the cell-by-gene count matrix was log-normalized using the NormalizeData function. The top 2000 most variable RNA features were selected for scaling and dimensionality reduction using the FindVariableFeatures function. Features were scaled and centered using the ScaleData function without regression of any variables. Default parameters were used across steps.

For ADT, count layers were first split by donor using the split function ('f' = object$hto.donor). Next, count layers were normalized within each cell by centered log-ratio transformation using the NormalizeData function (normalization.method = 'CLR', margin = 2). Given the smaller number of cell surface features labeled relative to the whole transcriptome, all ADT features were scaled and centered using the ScaleData function. No variables were regressed during scaling. To populate the variable feature slot of the Seurat object without limiting features for downstream dimensionality reduction steps, FindVariableFeatures was run subsequently.

<u>Selection of dimensionality reduction components per modality for cross-donor integration</u>

Dimensionality reduction and clustering was next performed using the normalized and scaled ATAC, RNA, and ADT cell-by-feature matrices of each L1-L3 Seurat object. Similar workflows based on Signac and Seurat guidelines were followed for each object, with function parameters tuned to determine ideal feature loadings for downstream data integration across donor samples: For the ATAC modality, the RunSVD function was used to create a latent semantic indexing (LSI) dimensional reduction of the cell-by-peak matrix by truncated singular value decomposition. Next, the standard deviation of each LSI component across cells was inspected by ElbowPlot to determine feature loading for clustering and non-linear dimensionality reduction. A strong positive correlation between the first LSI component and the number of ATAC peaks across cells was confirmed by FeatureScatter, as expected. Following the conventional Signac workflow, the first LSI component was therefore excluded from downstream analyses. Next, the cell-cell shared nearest-neighbor ('snn') graph was computed using the FindNeighbors function, with LSI component loading ('dims') as follows: L1 2:50, L2 2:30, and L3 2:11. Cells were clustered within the snn graph using the FindClusters function, with 'resolution' parameter values as follows: L1 1.0, L2 0.5, and L3 0.4. Next, Uniform Manifold Approximation and Projection (UMAP) of the snn graph was performed using the RunUMAP function with identical LSI component loading ('dims') as used in FindNeighbors. Finally, cell clustering was visualized by DimPlot. Both 'dims' and 'resolution' were tuned for L1-L3 objects to determine ideal LSI component loading before donor integration.

For the RNA modality, linear dimensionality reduction of the cell-by-gene matrix was performed by principal component analysis (PCA) using the RunPCA function. Similar to ATAC data analysis, a standard workflow of FindNeighbors, FindClusters, and RunUMAP was employed for cell clustering and visualization. In contrast to ATAC analysis, RNA PC1 was retained for all L1-L3 analyses. The number of RNA PC 'dims' was tuned as follows: L1 1:50, L2 1:50, and L3 1:35. Clustering 'resolution' was tuned as follows: L1 0.45, L2 1.7, and L3 0.7. Otherwise, default parameters were used throughout. RNA-based cell clustering data before cross-donor integration were not used in formal analyses.

ADT data were processed similar to the RNA modality using a standard workflow of RunPCA, FindNeighbors, FindClusters, and RunUMAP across L1-L3 analyses. The number of ADT PC 'dims' was tuned as follows: L1 1:50, L2 1:50, and L3 1:25. Clustering 'resolution' was tuned as follows: L1 0.84, L2 0.5, and L3 1.0. Otherwise, default parameters were used throughout. ADT-based cell clustering data before cross-donor integration were not used in formal analyses.

<u>Cross-donor modality integration</u>

All cryopreserved peripheral blood and tonsil mononuclear cell samples were thawed together and multiplexed for TEAseq to reduce sequencing batch effects. However, to account for possible technical batch effects between sample collection days and peripheral blood versus tonsil tissue processing protocols, batch correction across all eight donors was applied per modality for L1-L3 analyses. Based on a contemporaneous benchmarking study of multiple single-cell data integration approaches (*183*), Harmony (*184*) was implemented for batch correction of all three TEAseq modalities. Harmony can leverage low-dimensional embeddings from LSI or PCA to project cells from different samples into a new, shared embedding where cells are primarily grouped by biological types rather than technical variables specific to each sample. Harmony implementation varied between modalities and objects as detailed below:

To accommodate the ATAC 'ChromatinAssay' format, Harmony was implemented using the RunHarmony function of the harmony (1.2.0) library, in contrast to the layer-based integration for v5 Seurat assays. For L1-L3 analyses, parameters were specified as follows:

- group.by.vars = 'hto.donor' (integrating across HTO-demultiplexed donors)
- project.dim = FALSE (no additional computation of feature loadings per dimension)
- seed = 26 (for reproducibility)
- dims.use = 2:50 (for L1), 2:30 (for L2), and 2:11 (for L3) based on pre-integration tuning
- reduction.use = 'lsi.atac' (original LSI dimensional reduction)
- reduction.save = 'lsi.atac.harmony' (harmonized ATAC embedding)

For cross-donor integration of RNA and ADT data in each L1-L3 object, v5 assay layers were first split by donor using the split function ('f' = object$hto.donor). Next, Harmony integration was implemented for both RNA and ADT modalities separately using the Seurat IntegrateLayers function (method = 'HarmonyIntegration', seed = 26, orig.reduction = 'pca.rna' or 'pca.adt', new.reduction = 'pca.rna.harmony' or 'pca.adt.harmony').

<u>Selection of integrated embedding components for trimodal cell clustering</u>

Following cross-donor integration, the number of harmonized LSI (ATAC) and PC (RNA and ADT) dimensions as well as clustering resolution were tuned for each L1-L3 object as detailed below. For all data modalities, a standard workflow of ElbowPlot, FindNeighbors, FindClusters, RunUMAP, and DimPlot was used.

For the ATAC modality, all harmonized 'lsi.atac.harmony' components were used for clustering and UMAP visualization ('dims'): L1 1:49, L2 1:29, and L3 1:10. Given that the first pre-integration LSI component was excluded, the total number of projected 'lsi.atac.harmony' components decreased by one. For cell clustering, 'resolution' was tuned as follows: L1 0.45, L2 0.5, and L3 0.4.

For the RNA modality, the number of harmonized 'pca.rna.harmony' components for clustering and UMAP visualization was tuned as follows: L1 1:50, L2 1:50, and L3 1:35. Clustering 'resolution' was tuned as follows: L1 0.45, L2 1.0, and L3 0.7. For visualization purposes only, the UMAP embedding was rotated by 180° by applying a two-dimensional transform to the UMAP coordinate matrix. As rotation is a rigid transform, all pairwise Euclidean distances between cells were retained for visualization without changing the underlying neighbor or clustering data structure. Complete code details for the UMAP coordinate matrix rigid transform are provided in our GitHub repository.

For the ADT modality, the number of harmonized 'pca.adt.harmony' components for clustering and UMAP visualization was tuned as follows: L1 1:50, L2 1:50, and L3 1:25. Clustering 'resolution' was tuned as follows: L1 0.84, L2 0.7, and L3 0.8.

Trimodal clustering and dimensionality reduction

Harmonized embeddings for ATAC, RNA, and ADT were next subjected to three-way weighted nearest-neighbor (3WNN) analysis following Seurat developer guidelines (*41*). WNN analysis provided an unsupervised framework to learn ATAC, RNA, and ADT modality weights per single-cell in our TEAseq dataset. The WNN workflow was implemented for all L1-L3 Seurat objects using the core FindMultiModalNeighbors function, with parameter tuning as below:

- reduction.list = list('lsi.atac.harmony', 'pca.rna.harmony', 'pca.adt.harmony')
- dims.list = list of ATAC, RNA, and ADT components from Harmony, correspondingly:
    - L1 - 1:49, 1:50, 1:50
    - L2 - 1:29, 1:50, 1:50
    - L3 - 1:10, 1:35, 1:25

The output weighted shared nearest neighbor graph ('wsnn') was then used for trimodal cell clustering by calling the FindClusters function, with L1-L3 parameters tuned as detailed below:

- graph.name = 'wsnn'
- algorithm = 3 (smart local moving (*185*) as outlined by the WNN analysis (*41*) developers)
- resolution = 0.2 for L1, 1.08 for L2, and 0.6 for L3 Seurat objects

For visualization of L1-L3 trimodal clusters, the 3WNN structure was used to compute a UMAP embedding by calling the RunUMAP function with 'nn.name' specified as 'weighted.nn'. L1-L3 trimodal UMAP embeddings were visualized using the DimPlot function, with increasing 'pt.size' for L2 and L3 objects given lower cell counts (0.5 for L1, 1.5 for L2, and 2 for L3).

GeneActivity (ATAC Modality)

To infer gene expression 'activity' of each gene based on chromatin accessibility, the Signac GeneActivity workflow (*175*) was leveraged. First, GeneActivity extracts coordinates for each gene, including the upstream 2 kbp region to include promoter region accessibility. Next, ATAC fragments mapping to each extended gene locus per cell are automatically counted by the FeatureMatrix function. GeneActivity was implemented by identical workflows for all L1-L3 objects, with parameters modified to include noncoding and wide gene regions as detailed below:

- biotypes = NULL (default 'protein_coding' modified to include all noncoding genes)
- max.width = NULL (default modified to include all genes regardless of locus width)

- features = rownames(object@assays$RNA) (default NULL filters for protein-coding genes only, which was modified to include all gene names shared with the RNA assay based on the common hg38 cellranger-arc reference genome file used for alignment)

Next, the output cell-by-feature count matrix was used to create a new v5 assay named 'ACT' per L1-L3 object (CreateAssayObject). GeneActivity counts were log-normalized using the NormalizeData function using the median count number as a scaling factor (scale.factor = median(object$nCount_ACT)) as recommended. Finally, all GeneActivity features were centered and scaled using the ScaleData function.

<u>chromVAR (ATAC Modality)</u>

To quantify differences in transcription factor (TF) motif accessibility between single-cell epigenomic states, we performed chromVAR analysis (*70*). chromVAR leverages known TF motif information to identify ATAC peaks containing each motif, then compares accessibility of these regions in each cell to an expected value derived from the accessibility profile of all cells. To correct technical biases, chromVAR compares accessibility of motif-containing peaks to a background peak set with matching GC-content and average accessibility. Finally, the output bias-corrected cell-by-motif 'deviation score' matrix can be used to analyze varying accessibility of a given TF motif across single-cell states.

To implement chromVAR, we first restricted the ATAC feature matrix to regions found only on standard chromosomes in the 'BSgenome.Hsapiens.UCSC.hg38' build. Next, we extracted a position frequency matrix (PFM) model of TF motif binding profiles from the JASPAR2020 database (*186*) using the getMatrixSet function (collection = 'CORE', tax_group = 'vertebrates', all_versions = FALSE). PFM information was added to L1-L3 objects using the AddMotifs function. Finally, the RunChromVAR function was executed, storing the output cell-by-motif deviation score matrix as a new 'chromvar' assay. Scaled chromVAR scores for TFs of interest were visualized by ComplexHeatmap (2.20.0) across 3WNN Tfh-like and Tcm subclusters of the L3 Seurat object, with hierarchical clustering of both rows and columns (fig. S9B).

<u>SCENIC</u>

To infer gene regulatory networks specific to distinct single-cell states within each L1-L3 object, SCENIC (*69*) analysis was performed using the RNA modality. First, SCENIC builds an initial TF-centered co-expression network featuring genes correlated with a given TF. To remove false positive and indirect TF-target gene relationships, these initial modules of highly co-expressed TFs and candidate target genes are filtered based on DNA-binding motif enrichment for each TF. This process yields 'regulons' of TFs and predicted directly regulated genes, as supported by co-expression and motif enrichment data. Finally, regulons are scored per cell to yield a regulon area under the curve (AUC) enrichment score matrix, where higher scores for a given cell and TF indicate that its predicted target genes are among the most highly expressed genes in that cell.

The R implementation of SCENIC (1.3.1) was applied to each L1-L3 TEAseq object separately using similar workflows. First, tables of ranked TF motif scores by gene for both search spaces around gene transcription start sites (TSS) were extracted from v10 of the Stein Aerts Lab TF motif collection (mc_v10_clust):

- hg38_10kbp_up_10kbp_down_full_tx_v10_clust.genes_vs_motifs.rankings.feather
- hg38_500bp_up_100bp_down_full_tx_v10_clust.genes_vs_motifs.rankings.feather

To accommodate formatting expected by the R implementation of SCENIC for both motif tables, the 'motifs' column was renamed as 'features' and moved to be first (GitHub Issue #471). Next, human TF annotations matching the v10 motif collection (motifs-v10nr_clust-nr.hgnc-m0.001-o0.0.tbl) were extracted for downstream motif enrichment steps using RcisTarget (1.23.1). The core scenicOptions object was then initialized including both modified motif ranking tables using the initializeScenic function. Input RNA expression data were filtered using the geneFiltering function based on minimum detection in 1% of cells and total counts equivalent to 3% times the total number of cells (default parameters). The output filtered RNA expression matrix was then log-transformed before GRN modeling using GENIE3 (1.26.0). Finally, a standard workflow using core SCENIC functions was employed to build and score regulons of TFs and putatively targeted genes per cell: runSCENIC_1_coexNetwork2modules, runSCENIC_2_createRegulons, and runSCENIC_3_scoreCells (using AUCell (1.26.0) and the log-transformed filtered RNA count matrix). The output cell-by-regulon AUC score matrix was then appended to each L1-L3 object as a new Seurat v5 assay (CreateAssayObject) for further downstream analysis.

For heatmap visualization of differential regulon activity across clusters of the L3 TEAseq object (fig. S9A), the top nine regulons with 'RelativeActivity' score > 0.5 per cluster were extracted. For TF regulons with both direct and indirect ('extended') motif annotations in the mc_v10_clust database, the direct regulon was selected for visualization. Hierarchical clustering by the ComplexHeatmap function was performed for both cluster rows and regulon columns.

<u>Subclustering and annotation approach for TEAseq L1-L3 cell states</u>

Clusters resolved by 3WNN analysis in each L1-L3 TEAseq object were manually annotated based on differentially accessible ATAC peaks (DAP), expressed gene transcripts (DEG), and expressed surface proteins (DEP). To facilitate cluster annotation based on accessibility data, GeneActivity-based differentially accessible gene (DAG) scores were used. TF activity inferred by ATAC-based chromVAR and RNA-based SCENIC analyses further informed cluster annotation. To determine differential feature expression between all 3WNN clusters within each L1-L3 object, the Seurat FindAllMarkers function was implemented per modality as follows:

- DAP (ATAC), DAG (ATAC), DEG (RNA), and DEP (ADT) features between clusters were determined using default FindAllMarkers parameters
    - Coordinates for each DAP were converted to a GRanges object and supplied to the Signac ClosestFeature function to determine the nearest annotated gene

- chromVAR (ATAC)
  - mean.fxn = rowMeans (In contrast to log-normalized modalities, chromVAR returns TF motif *Z*-scores. As recommended by the Signac developers, the rowMeans function was used to determine the average difference in *Z*-scores between clusters rather than computing $log_2FC$)
  - Output 'pct.1' and 'pct.2' columns were renamed for clarity as non-zero chromVAR *Z*-scores do not connote 'positive' expression as for other modalities
  - Common TF names for each JASPAR2020 PFM ID were appended for legibility
- SCENIC (RNA)
  - mean.fxn = rowMeans (SCENIC regulon AUC scores are not log-transformed. As above, the average difference in regulon score was computed rather than $log_2FC$).
  - min.pct = 0, logfc.threshold = 0 (default thresholds were lowered as the mean difference in regulon AUC score between clusters can often be less than 0.1)
- FindAllMarkers results for each modality are provided as supplementary data files (data file S1 for L1 TEAseq 3WNN clusters, data file S2 for L2, and data file S3 for L3)

Statistical significance for all differentially expressed features was defined as $P < 0.05$ using a Wilcoxon rank-sum test (test.use = 'wilcox' by default) including a Bonferroni correction for multiple comparisons. Differentially expressed features from each modality were interpreted in context of relevant human literature (*18, 41-43, 46, 187*), as well as fundamental studies performed in mice, referenced in cluster lists grouped by L1-L3 object below. Clusters were primarily annotated using common names of major immune cell subsets, and secondarily based on features enriched in clusters belonging to the same major subset. Preferential enrichment of a given cluster in tonsil versus peripheral blood further influenced annotation.

TEAseq 3WNN L1 cluster annotation (primary immune cell lineages)

The L1 object comprised 15 distinct clusters based on 3WNN analysis of all 32,206 peripheral blood and tonsil mononuclear cells following data preprocessing, as detailed above. Multimodal cluster features are provided in fig. S2 and data file S1. In the B cell compartment, major subsets resolved included naive (NBC), memory (MBC), germinal center (GCB), and antibody-secreting cells (ASC). In the myeloid compartment, two clusters were resolved, one of which was enriched in CD16 expression across modalities (Myl vs Myl CD16). A single NK cell cluster was annotated based on enrichment in features such as CD16, CD56, and NKp46 proteins. Three clusters were resolved in the CD8 T cell compartment, including naive (CD8 Tn), memory or innate-like (CD8 Tm/Inn), and CD8 unconventional T cells (CD8 UTC). In contrast to CD8 Tn cells, CD8 Tm/Inn exhibited features of both conventional non-naive CD8 T cells (such as *PRF1* and *GZMB* transcripts), as well as innate-like subsets (such as TCR chains Vδ2 and Vα7.2). The remaining CD8 T cell cluster was annotated as CD8 UTC based on enrichment of features such as *GNG4* and *XCL1* accessibility, *ZNF683* and *IKZF2* transcripts, and several innate-like features including NKG2D protein (*43, 105-107*). Among CD4 T cells, five clusters were annotated:

naive (CD4 Tn), naive ribosomal protein gene-enriched (CD4 Tn Ribo), memory (CD4 Tm), Treg, and Tfh-like cells. CD4 Tn Ribo cells were enriched in ribosomal protein genes suggestive of a previously characterized poised naive state (*44, 45*). Treg cells were enriched in classic features such as *FOXP3* RNA and CD25 protein. Tfh-like cells were enriched in classic features such as *BCL6* and *TOX2* RNA, as well as CXCR5, PD1, and CD57 proteins. CD4 Tm cells lacked expression of CD45RA protein, Tfh, and Treg features, but were enriched in T helper cell features such as *ICOS*, *GATA3*, and *RORA* transcripts.

<u>TEAseq 3WNN L2 cluster annotation (T cells only)</u>

All CD4 and CD8 T cell 3WNN clusters were extracted from the L1 object for L2 3WNN subclustering analysis, with the exception of CD4 Tn Ribo as ribosomal protein genes were not a feature of interest for downstream comparisons of T cell states. 3WNN analysis of the remaining 23,442 T cells resolved 21 distinct clusters. Multimodal features of each 3WNN L2 T cell cluster are provided in fig. S6 and data file S2.

In the CD8 T cell compartment, six subclusters were resolved, including naive (CD8 Tn), memory (CD8 Tm), cytotoxic (CTL), gamma delta (gdT), PLZF-expressing innate-like (PLZF Inn), and unconventional T cells (CD8 UTC). CD8 Tn, CD8 Tm, and CD8 UTC clusters were annotated using similar markers as resolved in the L1 object. The CTL cluster was annotated based on enriched cytotoxicity-related features such as *GNLY* and *GZMH* transcripts and NKp46 protein. PLZF Inn cells were annotated based on high expression of *ZBTB16* and features of multiple innate-like subsets such as TCR V$\delta$2, TCR V$\alpha$7.2, and KLRB1 surface proteins as well as *CEBPD* and *RORC* transcripts. In contrast to PLZF Inn, gdT cells lacked TCR V$\alpha$7.2 protein expression and exhibited greater expression of both *TRGC1* and *TRDC* transcripts.

In the CD4 T cell compartment, 15 subclusters were resolved, including seven naive states (CD4 Tn 1-7), three Tfh-like states, and five nonTfh-like non-naive CD4 (nnCD4) T cell states. nonTfh nnCD4 T cell clusters included Th17, circulating Treg (cTreg), *RORC*-expressing Treg (*RORC* Treg), innate-like (CD4 Inn), and effector memory-like (CD4 Tem) cells. Th17 cells were annotated based on enriched expression of *RORC* and *CCR6* RNA as well as KLRB1 surface protein. Both *RORC* Treg and cTreg clusters were enriched in *FOXP3* RNA and CD25 surface protein, while *RORC* Treg exhibited greater *RORC* and *CCR6* RNA. In contrast, cTreg were enriched in circulatory features such as CCR7 and CD62L surface proteins as well as *KLF2* transcripts. CD4 Inn were annotated based on enriched expression of CD16, KLRB1, and CD35 (Complement Receptor 1) surface proteins as well as *ZBTB16* transcripts. CD4 Tem cells were annotated based on enriched expression of *BHLHE40*, *KLF2*, *IL2RA* and *ITGB1* (CD29) RNA in contrast to a cluster exhibiting signature features of both Tcm and Tfh-like cells (CD4 Tcm/fh). Relative to CD4 Tem, CD4 Tcm/fh were enriched in CD27 protein and *CCR7* RNA, as well as Tfh signature features such as *CXCR5*, *ICOS*, *BCL6*, and *TOX2* RNA. The remaining Tfh GC and Tfh IL10 clusters were annotated based on high expression of these signature Tfh features, as well as *IL10* transcripts in the latter.

TEAseq 3WNN L3 Cluster Annotation (Tfh-like and Tcm cells only)
To resolve finer Tfh gene expression states, all three Tfh-like clusters were extracted from the L2 T cell object for further 3WNN subclustering analysis. Among 3,657 cells, nine distinct L3 subclusters were resolved, including five GC-like versus three nonGC-like Tfh states, as well as one Tcm-like cluster. Multimodal features of each 3WNN L3 Tfh-like and Tcm cluster are provided in fig. S8 and data file S3.

The GC-like Tfh group was annotated based on enriched expression of features including PD1, ICOS, TIGIT, and OX40 surface proteins as well as *TOX2* RNA. Among GC-like Tfh, four clusters were annotated based on maximally enriched transcripts of interest, including *POU2AF1* (Tfh-BOB1), *CXCL13* (Tfh-CXCL13), *IL10* (Tfh-IL10), and *NFATC1* (Tfh-NFATC1). The remaining GC-like Tfh cluster exhibited absent or intermediate expression of these signature transcripts, and was correspondingly annotated as Tfh-Int. Among nonGC-like Tfh, one cluster was annotated based on maximal expression of AP1-family TF genes including *JUN* and *FOSB* (Tfh-AP1). A nonGC-like Tfh cluster enriched in circulatory features such as CD62L and CCR7 were annotated as Tfh-Circ. Tfh-Resting cells were annotated based on high accessibility of *CXCR5* and *IKZF1* (IKAROS) relative to other nonGC-like Tfh, in the absence of GC-like surface protein features such as PD1, ICOS, and TIGIT. Finally, Tcm were annotated based on minimal expression of Tfh signature features, maximal expression of Tcm signature features including *KLF2*, *CCR7*, and *ANXA1* RNA, as well as maximal enrichment in peripheral blood samples relative to the eight Tfh states.

TEAseq 3WNN L4 Object (Tfh-like cells only)
For downstream TEAseq analyses involving Tfh states only, all eight Tfh subclusters were extracted from the L3 Seurat object to create a L4 object that excluded Tcm. No further dimensionality reduction or subclustering of the resulting 2,996 Tfh cells was performed.

Determining donor and tissue composition per 3WNN cluster
Enrichment of 3WNN clusters in peripheral blood versus tonsil samples was determined using unenriched samples only (fig. S1). To correct for unequal cell numbers between donors, cluster cell counts were converted to proportions of the total cell count per donor. Next, these within-donor proportions were normalized across all donors such that donor representation per cluster summed to 1. Finally, normalized donor proportions per cluster were ordered by relative tonsil enrichment and visualized by stacked barplot, as in Fig. 1D for L1 3WNN clusters and fig. S7C for L2 3WNN subclusters. To compare the overall distribution of L3 3WNN subclusters within tonsil versus peripheral blood samples, both unenriched and CD4-sorted samples were used without normalization based on sample cell counts.

TEAseq L1 object cross-modality comparisons

To determine the relative contribution of each TEAseq modality to 3WNN L1 cluster identity, modality weights computed by the FindMultiModalNeighbors function were extracted. Weights per cell for each modality were visualized by ternary plot, grouped by L1 cluster (fig. S4) created using the ggtern function of the ggtern package (3.5.0). Next, modality weights were averaged across cells per cluster, and visualized for comparison on the same ternary plot (Fig. 1E). To visualize differences in dimensionally reduced embeddings between modalities, all fifteen 3WNN L1 cell identities were projected onto UMAP embeddings computed using each ATAC, RNA, and ADT modality separately.

To quantify differences in clustering between modalities, fifteen L1 clusters were resolved per modality as detailed above (section titled 'Selection of dimensionality reduction components per modality for cross-donor integration'). Next, the adjusted mutual information (AMI) between each set of unimodal cluster identities was determined as a metric of clustering similarity corrected for chance (*55-57*). AMI was computed using the sklearn$adjusted_mutual_info_score function of the scikit-learn (1.6.1) package in a reticulate (1.37.0) interface for python (3.9) within the same RStudio environment used throughout TEAseq analyses. AMI scores for each comparison were visualized by barplot (Fig. 1G), wherein a maximum score of 1 indicates complete agreement between cluster labels from each modality.

While AMI is agnostic of cluster naming, all fifteen clusters per L1 unimodal analysis were annotated based on differential feature expression (data file S1, refer to section above titled 'Subclustering and annotation approach for TEAseq L1-L3 cell states'), as visualized by the DotPlot function of Seurat (fig. S5C). Clusters were annotated and named similar to the L1 3WNN analysis, with the exception of the unimodal ADT analysis, in which clusters of mucosal-associated invariant T cells (MAIT), CD14 versus CD16 surface protein-expressing monocytes (CD14 Mono and CD16 Mono), and dendritic cells (DC) were resolved. Further, a single antigen-experienced B cell (Ag Exp B) cluster was annotated in the unimodal ADT analysis, in contrast to separate MBC, GCB, and ASC clusters for unimodal ATAC and RNA analyses.

Comparisons of L2 Tfh versus nonTfh Groups

To identify genome-wide features enriched in all Tfh clusters relative to nonTfh, a new annotation metadata column was appended to the L2 Seurat object wherein Tfh GC, Tfh IL10, and CD4 Tcm/fh identities were retained while all nonTfh cluster identities were merged. Differential accessibility of ATAC peaks and expression of RNA transcripts were then compared between all Tfh clusters and the nonTfh group using the Seurat FindAllMarkers, with lowered fold-change and percent detection thresholds for initial data exploration (logfc.threshold = 0, min.pct = 0). The resulting DAP and DEG tables were then filtered based on average $\log_2$FC > 0 and $P < 0.05$ by Wilcoxon rank-sum test with Bonferroni correction for multiple comparisons. The closest annotated gene for each DAP was determined as detailed above before extracting all unique enriched feature names per cluster. Next, enriched features of Tfh GC, Tfh IL10, and CD4 Tcm/fh clusters but not the nonTfh group were determined for both ATAC and RNA modalities. Features specific to the Tfh group in both ATAC and RNA analyses were considered multimodal 'core Tfh features' (data file S2). Overlap between ATAC and RNA feature sets was visualized by Venn diagram (fig. S7E). Finally, ADT expression between L2 Tfh and nonTfh groups was compared using the FindMarkers function, with enrichment defined as $|\log_2 FC| > 0.3$ and $P < 1e\text{-}30$ by Wilcoxon rank-sum test (Fig. 1I, data file S2).

Determining Th17 transcriptional signature enrichment across Tfh states

To assess Th17-polarization across Tfh states, a gene signature was curated from relevant human literature detailing enriched features in the Th17 subset (*72-77*) (Table S8). This Th17 gene signature was scored across all cells in the L3 Seurat object using the AddModuleScore_UCell function of the UCell package (2.8.0) (*188*). Th17 signature UCell scores were then compared between each L3 3WNN Tfh-like cluster using the Seurat DotPlot function (Fig. 2F).

Annotation of candidate *cis*-regulatory elements contained within DAP regions

Genomic regions for each DAP of interest were explored in the UCSC Genome Browser (*140*). To determine whether each DAP may contain candidate *cis*-regulatory elements (cCRE), the 'ENCODE cCRE' track was referenced (*68*). cCRE included promoter-like (PLS), proximal enhancer-like (pELS), and distal enhancer-like (dELS) sequences as defined by the ENCODE Project based on cCRE proximity to the TSS of a given gene as well as ChIPseq data in humans:

- PLS exhibit high DNase and H3K4me3 signals and are < 200 bp from the TSS
- pELS exhibit high DNase and H3K27ac signals and are < 2 kbp from TSS, as well as low H3K4me3 signal if < 200 bp from the TSS
- dELS exhibit high DNase and H3K27ac signals and are > 2 kbp from the TSS

Following this approach, several DAPs of interest in the TEAseq dataset were annotated:

- *CXCR5* PLS and dELS (Fig. 1J and fig. S7)
- *ICOS* dELS (fig. S7)
- *GNG4* PLS, pELS, and multiple dELS (Fig. 5A, C, H, and fig. S20)

Correlation of ATAC Peak Accessibility with RNA Expression

Inference of CRE activity in peaks of interest was further assessed by correlating accessibility of each peak with RNA expression using the Signac LinkPeaks function (*175*). LinkPeaks was implemented using default parameters, yielding Pearson correlation coefficients for each peak within 500 kb of gene TSS with RNA expression of that gene. For each peak an expected correlation coefficient was computed based on GC content, accessibility and length. The observed and expected correlation coefficients were then compared, yielding a *Z*-score and associated *P*-value. All output 'Links' were filtered by $P < 0.05$ and visualized by the Signac CoveragePlot function, wherein the 'Links' track displays arcs between each peak and the TSS region colored by the Pearson correlation coefficient. Identical LinkPeaks workflows were performed for both *CXCR5* (Fig. 1J) in L2 and *GNG4* (Fig. 5A, data file S7) in L3 objects.

Determining co-accessibility of *GNG4* ATAC peaks in Tfh

To infer networks of co-accessible CRE for *GNG4*, cicero (1.3.9) analysis was performed (*170*). First, the L4 Tfh subset Seurat object was converted to a Monocle 3 (1.3.7) (*189*) cell_data_set object (as.cell_data_set) before preparing the required input object for Cicero (make_cicero_cds) including 'reduced_coordinates' from the Tfh 3WNN UMAP embedding. Finally, a standard Cicero analysis workflow was employed, including the core run_cicero, generate_ccans, and ConnectionsToLinks functions. The output co-accessibility 'Links' were filtered for *GNG4* (data file S7) then visualized by CoveragePlot in L4 GC versus nonGC-like Tfh groups (Fig. 5H).

Comparison of GC versus nonGC-like Tfh across TEAseq modalities

To identify differentially accessible and transcribed genes in GC versus nonGC-like Tfh, the Seurat FindMarkers function was executed for both GeneActivity (ACT) and RNA assays of the L4 Tfh subset Seurat object. To retain all ACT and RNA features for visualization using volcano plot (Fig. 3B-C) and downstream genome-wide meta-analysis, default filters of the FindMarkers function were lowered (min.pct = 0, logfc.threshold = 0, min.cells.feature = 0, min.cells.group = 0). For the GeneActivity assay, DAG were defined by $|\log_2 FC| > 0.5$ and Wilcoxon rank-sum test $P < 1e\text{-}10$ (Fig. 3B, data file S3). For RNA, DEGs were defined by $|\log_2 FC| > 1$ and Wilcoxon rank-sum test $P < 1e\text{-}20$ (Fig. 3C, data file S3). In both volcano plots, the displayed $\log_2$FC range was limited to exclude only non-statistically significant features to improve visualization. To relate DAG and DEG results and identify multimodal GC Tfh-enriched features, meta-analysis leveraging METAL (*190*) was performed. Features detected only in the ACT or RNA assay were excluded, yielding 20,338 total multimodal features for meta-analysis. For each gene, $\log_2$FC and Wilcoxon rank-sum *P*-values were extracted from the unfiltered FindMarkers differential expression results above. Using data from both ACT and RNA modalities, meta-analysis was performed (*190*), yielding a multimodal 'Meta $\log_2$FC' and 'Meta *P*-value' per gene where greater fold change indicates enrichment in the GC-like Tfh group (data file S3). To define multimodal features that were most enriched in GC-like Tfh, the lowest 10 Meta *P*-value genes were ranked by Meta $\log_2$FC (Fig. 3D).

GC versus nonGC-like Tfh epigenomic states were further compared at the ATAC peak level in the L4 Tfh subset object using the FindMarkers function, with lowered detection thresholds for initial data exploration (min.pct = 0, logfc.threshold = 0). DAPs were filtered based on $P < 0.05$ by a Wilcoxon rank-sum test with Bonferroni correction, and annotated using the Signac ClosestFeature function as described above (data file S7). DAPs found within the *GNG4* locus and enriched in GC-like Tfh were highlighted by CoveragePlot visualization (Fig. 5A).

<u>Comparison of ADT features between *GNG4* RNA$^+$ versus RNA$^-$ Tfh states</u>

To identify surface protein features of *GNG4*$^+$ Tfh in tonsil, the L4 Tfh subset object was filtered based on 'hto.tissue' identities as determined in Preprocessing Step 7. Next, tonsil Tfh were stratified as *GNG4* RNA$^+$ versus RNA$^-$ based on a normalized RNA count threshold of 1. For each paired *GNG4* RNA Tfh group per tonsil donor, CLR-normalized counts of each ADT feature were averaged. Mean expression of each ADT was then compared between the *GNG4* RNA$^+$ versus RNA$^-$ Tfh groups of each tonsil donor using a paired *t*-test with the two-stage step-up FDR procedure of Benjamini, Krieger, and Yekutieli in Prism (10.2.2) (data file S3). FDR-based 'Discoveries' of differentially expressed ADT features between Tfh groups were defined based on *Q*-value thresholds of $*Q < 0.1$ $**Q < 0.05$, then visualized using connected paired dot plots in Prism (Fig. 3H).

<u>Inference of putative regulatory TFs for *GNG4* expression</u>

Inferring putative TFs of the *GNG4*$^+$ Tfh state leveraged both ATAC-based motif enrichment and RNA-based SCENIC analyses. First, DAPs between *GNG4* RNA$^+$ vs RNA$^-$ cells in the L4 Tfh subset object were identified using the FindMarkers function with a significant threshold of $P < 0.05$ by Wilcoxon rank-sum test with a Bonferroni correction (data file S7). DAPs with $\log_2$FC > 0 were extracted to identify chromatin regions with enriched accessibility in *GNG4* RNA$^+$ Tfh. TF motifs within these DAP regions were identified using the Signac FindMotifs function relative to a background peak set with matching GC content and accessibility in Tfh. Enriched TF motifs in *GNG4* RNA$^+$ Tfh were defined based on fold-enrichment > 1.25 and $P <$ 1e-10 using a hypergeometric test with Benjamini-Hochberg FDR correction (data file S7).

TF regulon activity was assessed across L3 Tfh transcriptional states using SCENIC as detailed above. TFs inferred to regulate *GNG4* were extracted from the output 'regulonTargetsInfo' file (data file S7). Next, putative TF lists from ATAC-based motif enrichment versus SCENIC analyses were compared, taking the mutually identified TFs (BACH1 and NFATC1) for further analysis. To determine whether BACH1 and NFATC1 motifs were specifically present within the PLS, pELS, and dELS DAP regions of *GNG4* detailed above, motifmatchr (1.26.0) analysis was performed. First, position frequency matrix information for NFATC1 and BACH1 motifs were extracted from the JASPAR2020 database (collection = 'CORE', tax_group = 'vertebrates', all_versions = FALSE). Next, the matchMotifs function was called to return the maximum position weight matrix match score for NFATC1 and BACH1 motifs found within the *GNG4* PLS, pELS, and dELS regions that passed a *P*-value threshold of 0.05 (out = 'scores', p.cutoff = 0.05). Finally, motif match log-odds scores for NFATC1 and BACH1 within the PLS, pELS, and dELS regions of *GNG4* were visualized as a grid using the ComplexHeatmap package (Fig. 5C).

<u>Species conservation of *GNG4* putative CRE DNA sequence and Gγ4 amino acid sequence</u>

Conservation of the *GNG4* PLS, pELS, and dELS regions defined in Fig. 5A, as well as the entire *GNG4* locus was visualized in the UCSC Genome Browser using the 'Hiller Lab 470

Mammals' track ('470 mammalian genomes aligned with Multiz by Michael Hiller's Group') including PhyloP conservation scores (fig. S20). Conservation of the Gγ4 75-amino acid polypeptide chain was assessed using the UniProt 'Align' webtool between human (entry P50150) and mice (entry P50153).

**Xenium Prime Data Analysis**

Overview

Analysis of the Xenium Prime (XP) dataset featured several data preprocessing steps per tonsil sample followed by Sketch-based dimensionality reduction and clustering at two levels of resolution – all immune, stromal, epithelial, and endothelial subsets (L1) and non-naive CD4 T cells alone (L2). L1 and L2 objects were then used for analyses visualized in Figure 4 and supplementary figures S13 through S18. Related code to reproduce analyses is provided in our GitHub repository: https://github.com/theoldridgelab/TEAseqXeniumPrimeGNG4.

Computing Environment

XP analysis was performed within the CHOP high-performance computing cluster, primarily using the RStudio (2024.04.0) integrated development environment for R (4.4.0). For reproducibility, '26' was specified as the default seed for random number generation in all R-based analyses. Whole-slide image registration and cellular neighborhood analyses were performed in Python. Key packages used throughout analysis of XP data are listed in table S7.

XP data preprocessing

Preprocessed data from the Xenium Onboard Analysis (XOA) pipeline were imported for all six tonsil samples using a custom 'ReadXenium' function due to file formatting incompatibility between XOA (3.2.1.2) and the Seurat (5.1.0) version used at the time of our XP data analysis (GitHub Issue #9060). Imported XOA data were then used to create a Seurat object per sample, with donor demographic and sample technical information stored as metadata columns. The distribution of detected RNA probe counts (nCount_RNA) and unique features (nFeature_RNA) was inspected across cells and samples to determine thresholds for QC filtering. In addition to QC filtering performed by XOA, high-quality cells for each sample were further defined by nCount_RNA >= 50 and nFeature_RNA >= 45 thresholds. Following quality control filtering, all six samples were merged into a single Seurat object.

L1 XP object dimensionality reduction and clustering

To enable dimensionality reduction and clustering of all 6,290,476 cells in the L1 Seurat object, a standard Sketch-based downsampling procedure was performed. First, RNA probe counts were log-normalized using the NormalizeData function. Highly-variable features were selected using the FindVariableFeatures function for Sketch analysis. Finally, the SketchData function was used to create a downsampled 'sketch' assay of 10,000 cells per sample, for a total of 60,000 cells (method = 'LeverageScore'). Dimensionality reduction and clustering analysis was performed using the downsampled data. First, variable features were determined by FindVariableFeatures. To mitigate downstream clustering of distinct cell types based on probe counts (nCount_RNA) and unique features detected (nFeature_RNA), both technical variables were regressed during data centering and scaling using the ScaleData function (vars.to.regress). The RunPCA function was used for linear dimensionality reduction of downsampled sketch data. 30 PCs and clustering resolution of 1.0 were selected for downstream L1 XP analyses based on inspection of the ElbowPlot as well as an iterative process of clustering and UMAP visualization.

Clustering and UMAP dimensionality reduction of the L1 XP Seurat object was performed using a standard workflow of FindNeighbors, FindClusters, and RunUMAP. FindClusters 'resolution' was varied from 0.2 to 1.0 to determine an ideal parameter value for resolving major expected cell types. Differential feature expression between clusters resolved at each level of resolution was determined using the FindAllMarkers function. Finally, clustering, PCA, and UMAP data computed using 30 PCs and 1.0 clustering resolution were projected from the downsampled 'sketch' assay to the entire dataset using the ProjectData function.

<u>L1 XP object cluster annotation</u>

All 26 clusters of the L1 XP object were manually annotated based on DEGs (fig. S14, data file S4), spatial positioning, and published human reference atlases detailing immune, stromal, epithelial, and endothelial cell types (*18, 42, 121-124*). Among endothelial cells, arterial (Art Endo), venous (Ven Endo), and lymphatic (LEC) subsets were annotated. Mural cells (Mural) were annotated based on DEGs and proximity to endothelial cell types. In the epithelial compartment, surface (Surf Epi), intermediate (Int Epi), and basal (Basal Epi) clusters were annotated based on DEGs and relative positioning between clusters. 'Crypt Epi' cells were annotated based on epithelial DEGs and enriched positioning within crypt regions. In the stromal compartment, follicular dendritic cells (FDC), fibroblastic reticular cells (FRC), and trabecular fibroblasts (Trab Fib) were annotated by signature DEGs as well as biased positioning within B cell follicles versus lining B cell follicles or in connective tissue bands, respectively.

In the immune compartment, 15 distinct L1 clusters were annotated, including major B, T, granulocyte, and dendritic cell subsets. Among B cells, naive (NBC), memory (MBC), antibody-secreting (ASC), light zone germinal center (LZ GCB), and dark zone germinal center (DZ GCB) were annotated based on signature DEGs. In the T cell compartment naive (Naive T), cytotoxic (Cytotox), and non-naive CD4 (nnCD4) clusters were annotated. Annotated myeloid subsets included monocytes and macrophages (Mono/Mac), conventional dendritic cells (DC), plasmacytoid dendritic cells (PDC), extrathymic *AIRE*-expressing cells (eTAC), neutrophil-like granulocytes (Gran), and mast cells (Mast). A cluster enriched in cell-cycle-related genes and features of several immune cell lineages was annotated as 'Cycling'.

L2 XP object dimensionality reduction and clustering

To identify distinct Tfh versus nonTfh transcriptional states, subclustering analysis of all 330,994 L1 'nnCD4' T cells was performed. First, a downsampled 'sketch' assay including 60,000 cells from the L2 object was generated using a similar Sketch-based workflow as implemented for the L1 object, including NormalizeData, FindVariableFeatures, and SketchData functions. Highly variable features of the L2 sketch assay were determined by FindVariableFeatures, followed by regression of nCount_RNA and nFeature_RNA during data scaling and centering by ScaleData. Following dimensionality reduction by RunPCA, 15 PCs were selected for clustering and UMAP visualization. As above, the 'resolution' parameter of FindClusters was varied from 0.2 to 1.0 to determine an ideal clustering resolution. Clustering, PCA, and UMAP data computed using 15 PCs and 1.0 resolution were projected to the complete L2 dataset using the ProjectData function.

L2 XP object cluster annotation

16 L2 nnCD4 T cell subclusters were manually annotated based on DEGs (fig. S15, data file S4), spatial positioning, and relevant human literature (*18, 24, 25, 41, 46*). 10 Tfh versus 6 nonTfh clusters were defined by FindMarkers comparison between groups, with a DEG statistical significance threshold of $P < 0.05$ by Wilcoxon rank-sum test with Bonferroni correction. DEGs used to define the Tfh group included enriched *CXCR5*, *BCL6*, *PDCD1*, *IL21*, and *CXCL13* versus decreased *PRDM1*, *KLF2*, and *BHLHE40* relative to nonTfh (fig. S15, data file S4).

Six Tfh clusters were annotated based on maximal expression of particular genes of interest, such as *TOX2*, *S1PR2*, *PRDM1*, *CXCL13*, *NFATC1* and *CCR6* (Tfh TOX2, Tfh S1PR2, Tfh PRDM1, Tfh CXCL13, Tfh NFATC1, and Tfh CCR6). Two Tfh clusters were annotated based on preferential positioning in the germinal center dark zone (Tfh DZ) and mantle region (Tfh Mant). Tfh with a circulatory phenotype (Tfh Circ) were annotated based on positioning beyond the follicle and enrichment in migratory features such as *CCR7* and *SELL*.

Among nonTfh, resident memory-like cells (Trm) were annotated by enriched positioning in epithelial regions and *ITGAE* expression. The cluster with maximal expression of *FOXP3* and other Treg signature genes was annotated as 'Treg/fr'. Two central memory-like (Tcm 1, Tcm 2) clusters were annotated by positioning outside follicles and enrichment in migratory features. Relative to Tcm, conventional effector-like cells (Tconv) were annotated based on enrichment in TF, cytokine, and chemokine receptor features of T helper cells. Finally, Tconv and Tfh-like clusters exhibiting features of myeloid lineage cells (*CSF1R*, *CD68*, *ITGAX*) were annotated as 'Tconv Myl' and 'Tfh Myl'. As detailed below, additional analyses were performed to resolve Tfh-specific features relative to other lineages due to potentially contaminating signals from cell-to-cell transcript diffusion, inaccurate segmentation, and vertical overlap of cells (*130*).

Registration of Xenium Multi-Tissue Stain Mix and H&E whole-slide images

H&E-stained images were acquired for each tonsil sample following the XP assay to facilitate spatial analyses of single-cell transcriptional states. Registration between Xenium and H&E images was executed using wsireg, an open-source, graph-based multimodal whole-slide image registration toolkit designed for histopathological data. The wsireg package was employed to align H&E and Xenium Multi-Tissue Stain Mix images (including nuclear, membrane, and cell interior stains) by affine transformation and optimizing spatial transformations. Registration quality per tonsil sample was first evaluated by visual inspection of DAPI staining versus the deconvolved hematoxylin signal of H&E images in QuPath. Quantitative validation of image registration employed normalized mutual information, Dice coefficient, and Jaccard index metrics calculated on corresponding nuclear polygons delineated in the DAPI and H&E images.

XP tonsil cellular neighborhood analysis

Cellular neighborhood (CN) analysis was performed by identifying k = 40 nearest neighbors for each cell. Following image data preprocessing and annotation of L1 cell types, a machine-learning-based, k-nearest neighbor (k-NN) approach was leveraged to classify all tonsillar cells across six donors into one of 10 distinct CNs (*128*). Each CN was annotated by visual alignment with DAPI and H&E images in Xenium Explorer (3.2.0) as well as varying representation of L1 and L2 tonsil cell types resolved by RNA-based clustering analyses described above.

To facilitate comparisons of GC versus nonGC Tfh transcriptional states, CNs originally annotated as GC LZ (CN2) and GC DZ (CN5) were merged into a single GC mask (CN2). Cell centroids were rasterized into a fixed grid, followed by GC seed map cleaning using scikit-image morphology tools (closing, hole filling, small-object removal). Cells were reassigned to the merged GC CN if found within the cleaned GC mask and were at least a set interior margin from the GC boundary as well as a mantle-exclusion margin from the mantle region (CN6) (KD-tree). Otherwise, original CN labels of cells positioned outside the GC were retained.

Xenium Explorer visualization

Xenium Explorer (3.2.0) was used to visualize whole-slide images, transcripts, cellular neighborhoods, and cell types of interest across tonsil samples (Fig. 4, fig. S13, fig. S16, fig. S18). Whole-slide images were visualized with values for Opacity = 100, Brightness = 50, and Contrast = 10. Transcripts were viewed as Points (Circle), with size scale = 12 and Opacity = 100. Low-quality transcripts were filtered by XOA and not visualized. Cells were viewed as both Filled and Outlined, with Opacity = 100. Cells that passed XOA QC filtering but not second-pass QC filtering in our analysis were not assigned annotations and therefore were not visualized.

Defining features enriched in GC-localized Tfh

To determine features with high positive-predictive value (PPV) for GC-localization in Tfh, L2 XP object Tfh were defined as positive or negative for all 5,101 features based on a normalized signal threshold of 1. Using merged GC CNs as a spatial anchor (refer to section titled ‘XP tonsil cellular neighborhood analysis’), the percentage of feature-positive Tfh across all samples that localized to the GC was determined for all 5,101 features (data file S5). The PPV for all features was then ranked from highest to lowest (data file S5), and visualized by plotting GC percentage versus feature rank (Fig. 4L). To determine the relative % GC PPV of *GNG4* versus a panel of other conventionally GC-associated features, PPV was computed per donor, then compared between *GNG4* and each feature. A threshold of $Q < 0.05$ by paired Wilcoxon signed-rank test with Benjamini-Hochberg FDR correction was used to classify ‘discoveries’.

To enhance resolution of features restricted to Tfh residing within the merged GC CN, as well as the Tfh lineage generally, three comparisons were made using the FindMarkers function:

1) XP Tfh within the GC versus Tfh positioned outside the GC [RNA]
2) XP Tfh versus all other cells within the GC [RNA]
3) TEAseq Tfh versus all other 3WNN clusters of the L1 object [RNA]

All comparisons were initially made with lowered detection thresholds to allow complete exploration of the data (min.pct = 0, logfc.threshold = 0) (data file S5). Next, DEGs were defined by $P < 0.05$ using a Wilcoxon rank-sum test with Bonferroni correction. As TEAseq yields whole transcriptome data, the feature set considered for further analysis was subset to the 5,101-plex XP panel. The $\log_2$FC for each gene evaluated by all three comparisons was then visualized by scatterplot, with the color of each point corresponding to the TEAseq RNA modality (Fig. 4M). Next, features with positive enrichment in Tfh by all three comparisons (avg_log2FC > 0) were extracted to define a list of GC Tfh-enriched features (fig. S17, data file S5).

Evaluating gene expression in Tfh as a function of distance from LZ GC B cells

To quantify changes in Tfh gene expression based on proximity to LZ GC B cells, first normalized RNA data were extracted for a panel of genes (Fig. 4O) across XP L2 Tfh cells. Next, the Euclidean distance between each Tfh and the nearest annotated LZ GC B cell was determined. Tfh within 6 µm of LZ GC B cells were excluded as likely segmentation errors given the expected normal size of lymphocytes. Next, a Gaussian kernel-smoothing function was used to estimate average gene expression in Tfh over a grid spanning 6 to 100 µm (evaluated in 0.1 µm intervals with a smoothing bandwidth of 5 µm). Finally, the smoothed output data were visualized by plotting mean normalized RNA versus distance (Fig. 4O).

Comparison of Tfh populations within the GC

All L2 XP Tfh (235,811 cells) were defined as *GNG4* $RNA^+$ (52,532) versus $RNA^-$ (183,279) based on a normalized RNA signal threshold of >= 1. Next, the FindMarkers function was used to compare *GNG4* RNA groups, subset to Tfh residing within the merged GC CN (data file S5). Default thresholds were lowered for downstream visualization by volcano plot (min.pct = 0, logfc.threshold = 0, min.cells.feature = 0, min.cells.group = 0). DEGs were defined by $|\log_2\text{FC}| >$

0.5 and $P$ < 1e-10 using a Wilcoxon rank-sum test, then visualized using a volcano plot (Fig. 4P). A similar analysis was performed for *FOXP3* $RNA^{+}$ versus $RNA^{-}$ GC Tfh, again using a normalized RNA signal threshold of >= 1. DEGs were defined by $|log_2FC|$ > 0.25 as well as Wilcoxon rank-sum $P$ < 1e-5, then visualized using a volcano plot (fig. S18).

**Public data reanalysis**

Reanalysis of adult tonsil Visium HD and lymph node Visium v1 spatial transcriptomics data

Publicly available adult tonsil Visium HD and lymph node Visium v1 data were obtained from 10x Genomics (*137, 138*). Reanalysis of the provided 'loupe' files was performed using Loupe Browser (8.0.0). Graph-based clustering was used to define clusters in each dataset. DEGs between all clusters were determined, then filtered by $P$ < 0.05 adjusted using the Benjamini-Hochberg correction for multiple comparisons (data file S6). In each dataset, a single cluster was annotated as 'GC' based on enrichment in signature DEGs such as *BCL6, AICDA,* and *S1PR2* (Cluster 8 in tonsil Visium HD and Cluster 7 in lymph node Visium v1).

Reanalysis of mouse scRNAseq data across tissues as well as health and disease states

Publicly available scRNAseq data in mice across several contexts were obtained (*141-146*). Bone marrow, inguinal lymph node, and both normal spleen datasets were explored using the online Broad Institute Single Cell Portal (fig. S21). Gastrointestinal tissue data were explored using the online Cell x Gene environment. Mouse LCMV as well as B16 and MC38 tumor model data were explored in the Swiss Portal for Immune Cell Analysis. For all datasets, author-provided annotations were used to visualize cells in the provided dimensionality reduced embedding for parallel comparison with *Gng4* RNA expression.

Reanalysis of human scRNAseq data in longitudinally sampled lymph nodes post-vaccination

scRNAseq data were obtained from two studies of longitudinally sampled human lymph nodes following vaccination with BNT162b2 mRNA against SARS-CoV-2 (*24*) or quadrivalent inactivated influenza (*25*). In the BNT162b2 study, expression of *GNG4* RNA as well as a panel of CD4 and CD8 T cell subset-defining features was visualized across clusters defined by Borcherding *et al.* using the Seurat FeaturePlot and DotPlot functions (fig. S24A-S24C). Gene expression in the single 'GC Tfh' cluster (c3) was then compared to all other author-annotated Tfh and nonTfh clusters as a group across d28, d60, and d201 timepoints. The d110 timepoint was excluded due to low recovered cell numbers. Percent detection and average normalized expression of genes was visualized by DotPlot and resulting values were extracted for quantitative comparison of changes in 'GC Tfh' *GNG4* expression between timepoints (Fig. 5I).

In the quadrivalent inactivated influenza vaccination study, the provided object of subclustered Tfh-like cells included a single annotated 'GC' cluster (fig. S24D), and a level 1 clustering object of all T cell populations (fig. S24G). To enable comparison of gene expression in 'GC

Tfh’ versus other T cells, the ‘GC’ annotation from subclustered Tfh was applied to the L1 T cell object (fig. S24J). The DotPlot function was used to compare *GNG4* expression across clusters of the Tfh object (fig. S24F) as well as the L1 object using original annotations (fig. S24I) and the relabeled ‘GC Tfh’ cluster identity (fig. S24J). Next, *GNG4* and other features of interest were compared in GC Tfh versus all other Tfh and nonTfh clusters as a group across d0 through d180 timepoints. The d26 timepoint was removed due to low GC Tfh cell numbers. Timepoints corresponding to the same day post-vaccination in each year of the study were merged. Percent detection and average normalized expression of genes was visualized by DotPlot and resulting values were extracted for quantitative comparison of changes in *GNG4* expression by ‘GC Tfh’ between day <= 28 (‘Early’) and day >= 60 (‘Late’) timepoints (Fig. 5J).

Reanalysis of *GNG4* genome wide association studies (GWAS) in human arthritis

During reanalysis of published RA fine-mapping GWAS data (*157*), a discrepancy in annotation of the originally reported rs1188620266 variant was noted between versions of the Genome Aggregation Database (gnomAD) used at the time of the original study versus our recent reanalysis. Consultation with the original study authors confirmed that while the original study leveraged gnomAD v2.0 variant annotations based on the hg19 assembly, gnomAD v4.1.0 is the current major version used for hg38. Between versions, the variant originally annotated as rs1188620266 (1:235800357:CAA:C) was remapped to rs61512163 (1:235637057:CAA:C). Further, in gnomAD v4.1.0, the rs1188620266 annotation was reassigned to a different variant (1:235637057:C:A). In accordance with current naming standards and for consistency with the hg38 reference build used in our study, we have chosen to use the rs61512163 naming convention here.

For linkage disequilibrium (LD) analysis summarized in data file S9, pairwise LD statistics were computed using the online LDpair tool (https://ldlink.nih.gov/?tab=ldpair). For each pair of SNPs, analysis was run with all populations selected using the default value of ‘GRCh37’ selected for 1000 Genomes genome build.

**Gγ4 Antibody Conjugation**

Gγ4-specific antibodies (NSJ, RQ6130) were conjugated to PE using the Lightning-Link R-PE Conjugation Kit (Abcam, ab102918) following manufacturer recommendations. After adding conjugation reagent and quencher solutions, the concentration of the input Gγ4 antibody was 0.833 μg/μL. The Gγ4-PE conjugate was titrated to a range of 1:4500-1:5000 for flow cytometry intracellular staining (table S5) of human PBMC and tonsil mononuclear cell samples.

**Supplementary Figures:**

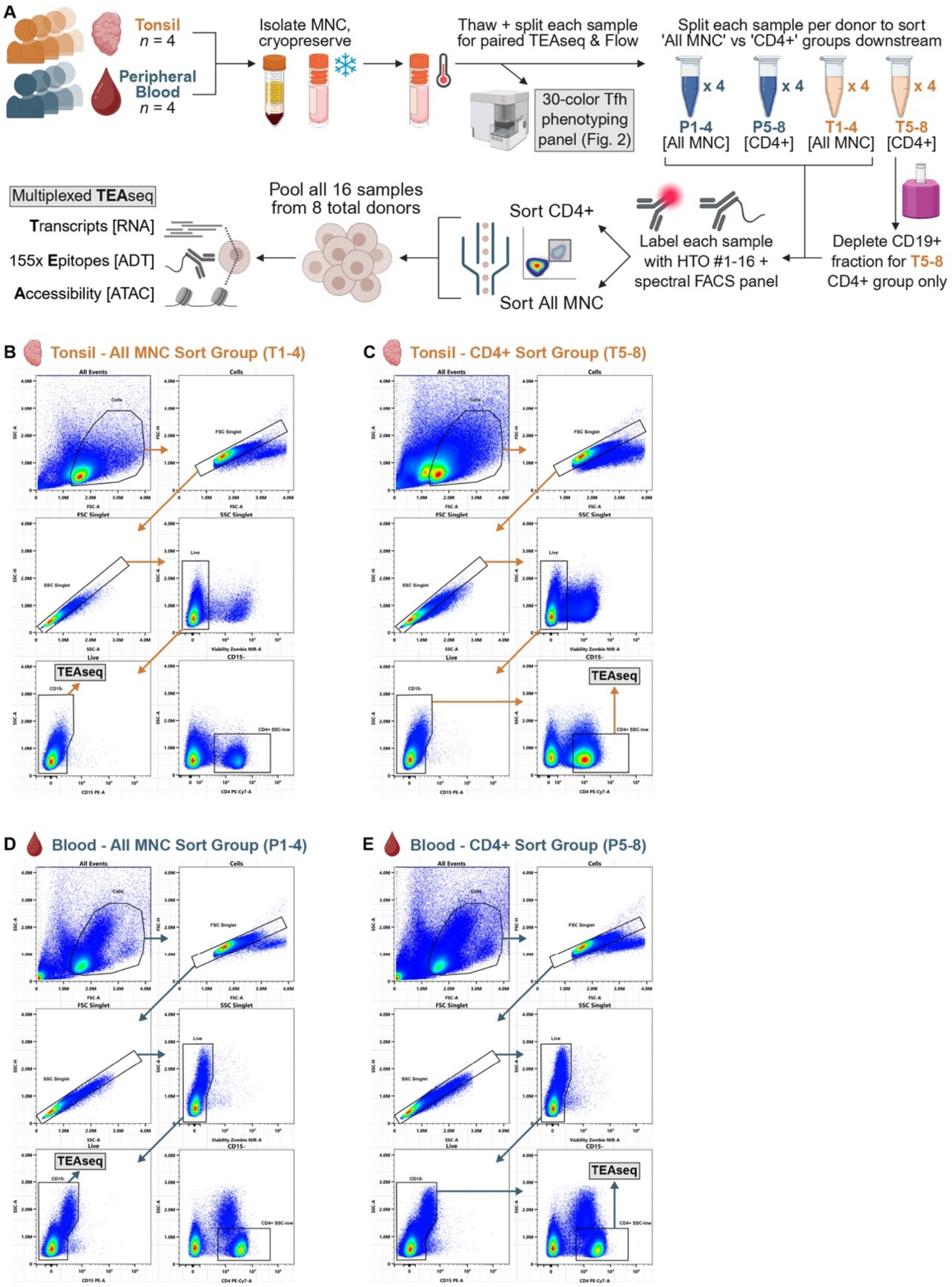


**Fig. S1 - Paired TEAseq and flow cytometry experimental design and cell sorting strategy.** **(A)** Experimental schematic summarizing paired TEAseq and spectral flow cytometry profiling of mononuclear cell (MNC) populations from peripheral blood (P, blue) and tonsil (T, orange) samples. **(B-E)** Representative pseudocolor plots from sorting all live cells ('All MNC') as well as CD4-enriched cells ($CD4^+$) in **(B-C)** tonsil and **(D-E)** PBMC samples.

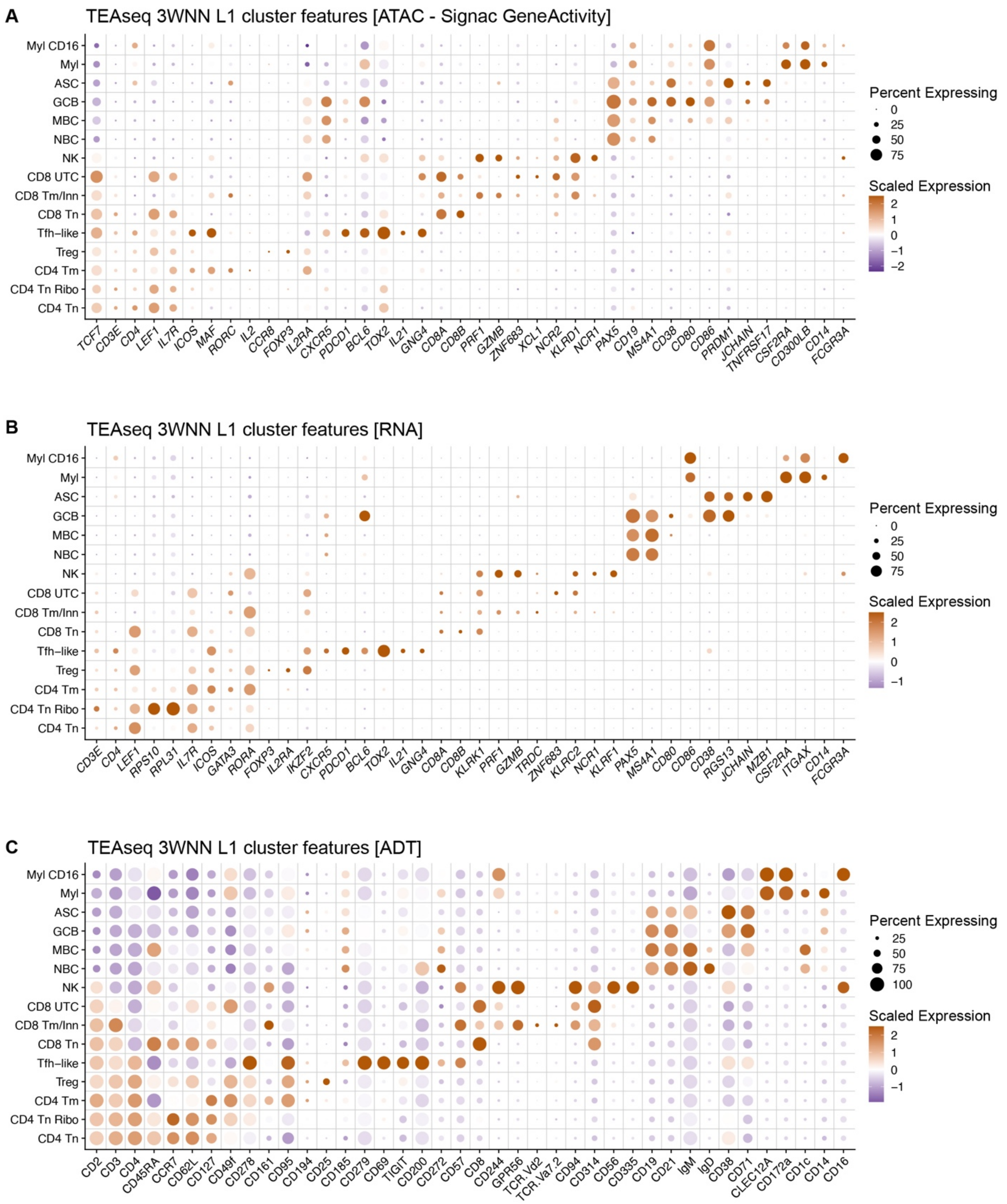


**Fig. S2 – Trimodal annotation of Level 1 mononuclear cell states in tonsil and peripheral blood TEAseq dataset. (A-C)** Scaled average expression and percent expression of differentially expressed features between all 15 clusters resolved by three-way weighted-nearest neighbor (3WNN) analysis of the Level 1 (L1) TEAseq dataset (32,206 cells). Three feature modalities are shown: **(A)** chromatin accessibility (from ATAC-based GeneActivity analysis), **(B)** mRNA transcripts, and **(C)** cell surface proteins (from antibody-derived tag (ADT) labeling).

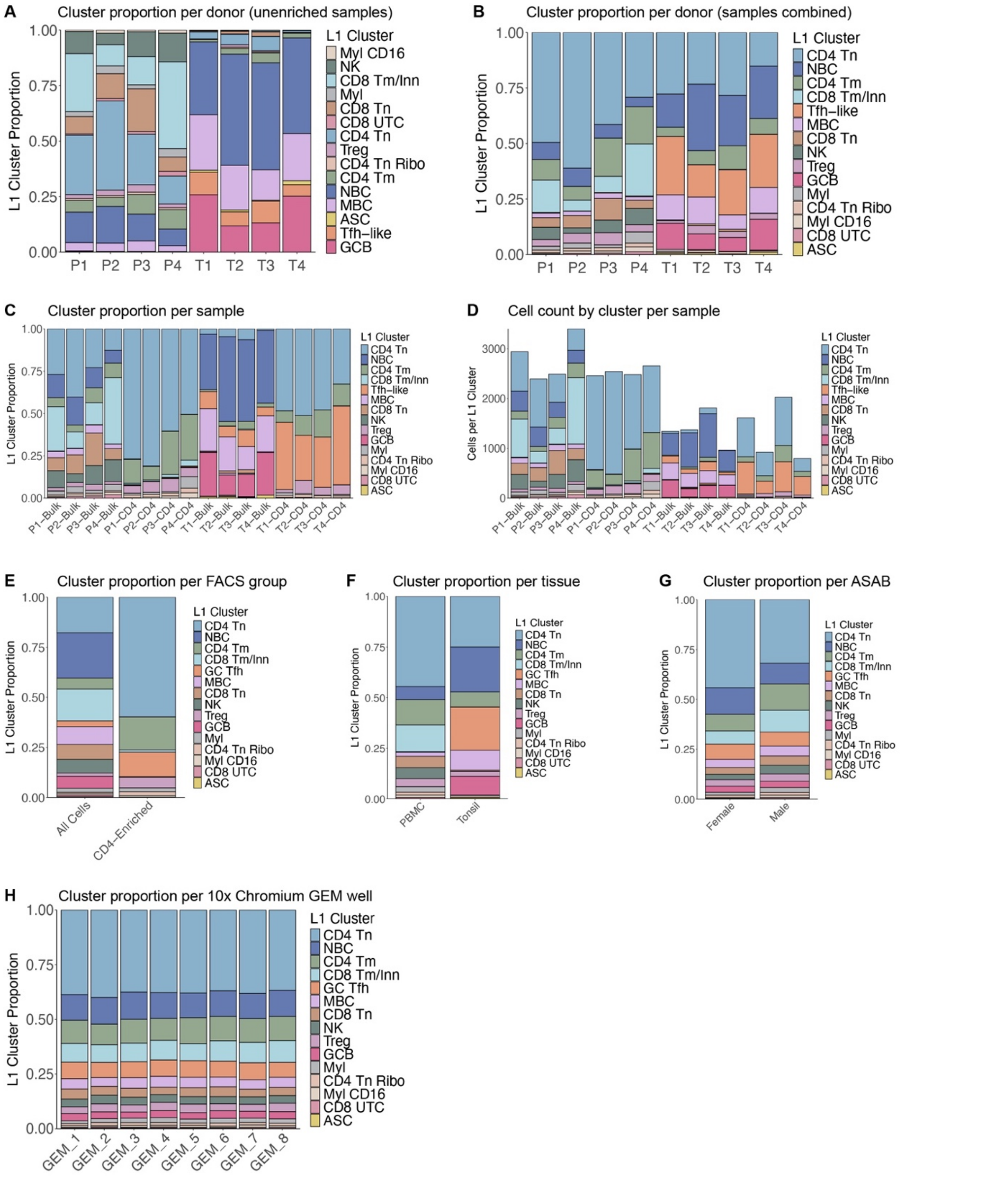


**Fig. S3 - Distribution of Level 1 cell types in TEAseq dataset across biological and technical variables. (A-C)** Stacked barplots of 3WNN TEAseq L1 cluster proportions across **(A)** 'Bulk' FACS samples for peripheral blood versus tonsil donors **(B)** both 'Bulk' and enriched 'CD4' samples per donor, and **(C)** all 16 FACS samples. **(D)** Stacked barplot of cell counts per cluster across all 16 FACS samples. **(E-H)** Stacked barplots of cluster proportions in samples across **(E)** FACS groups **(F)** tissues **(G)** assigned sex at birth, and **(H)** GEM wells.

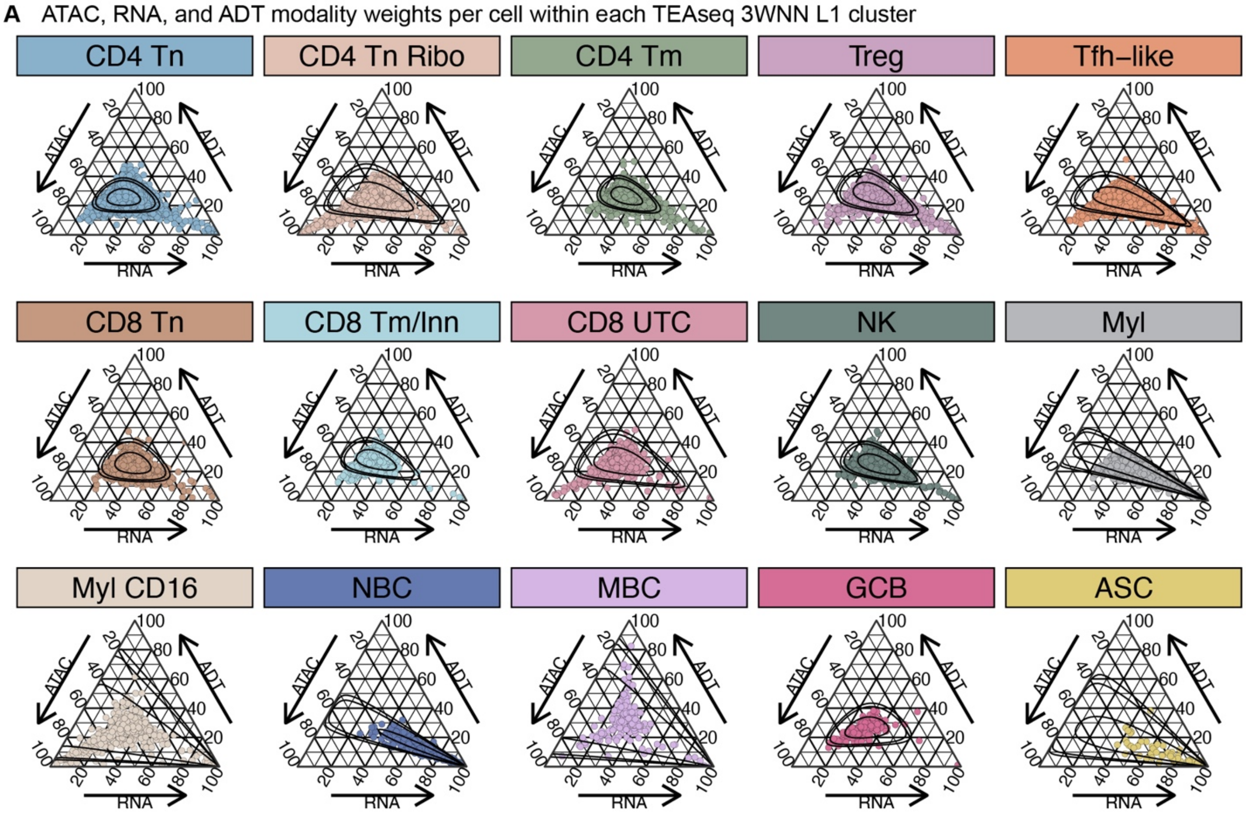


**Fig. S4 - Gene expression modality weights vary between cells of each TEAseq 3WNN L1 cluster. (A)** ATAC, RNA, and ADT modality weights per cell from 3WNN analysis of the L1 TEAseq object, visualized together using ternary plots and grouped by cluster. ATAC, RNA, and ADT weights per cell sum to one. Contours indicate 50, 90, and 95% confidence intervals around the cluster mean trimodal weighting value.

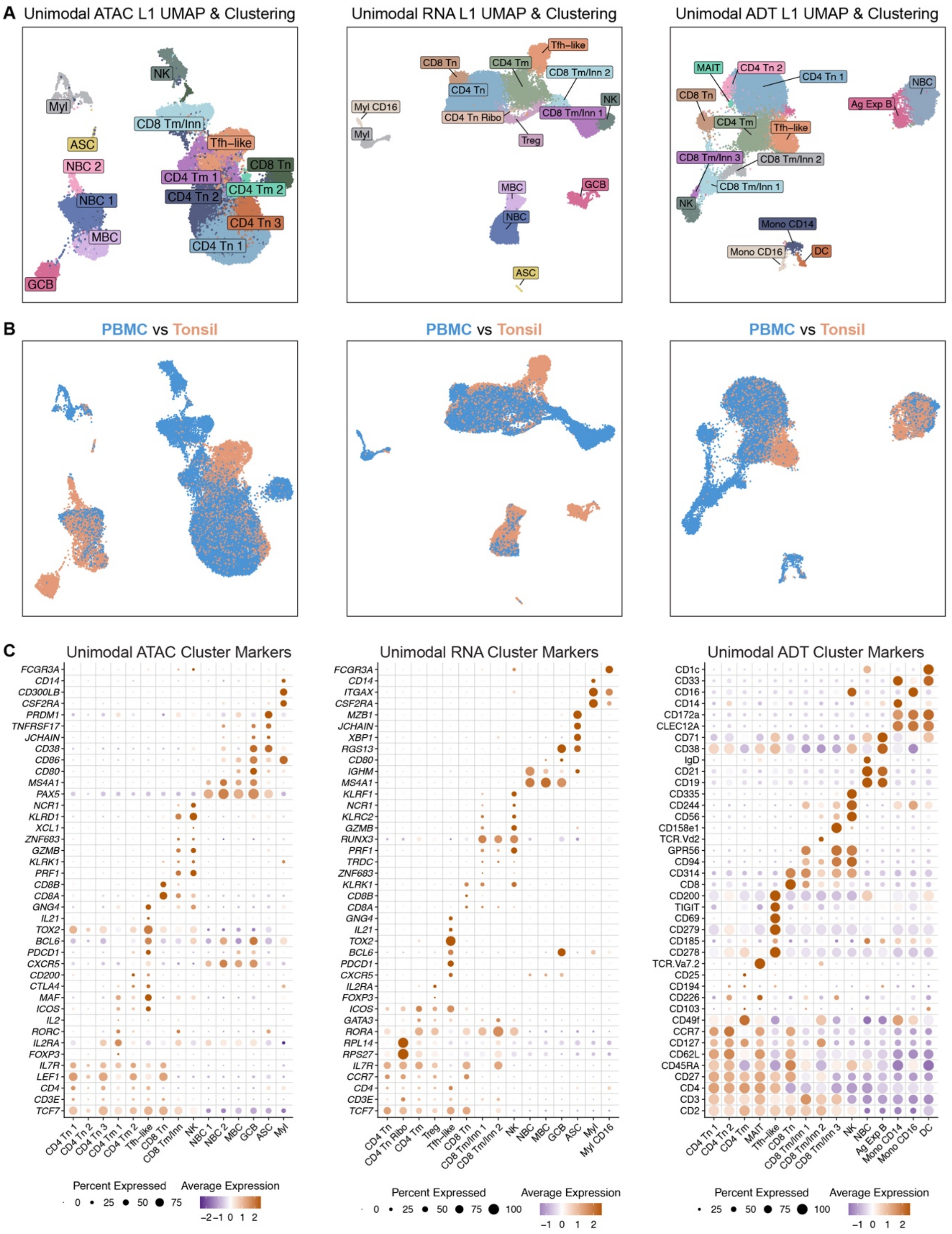


**Fig. S5 - Separate unimodal analyses of trimodal TEAseq data identify shared and distinct immune cell states. (A)** UMAP embeddings of clusters resolved by unimodal ATAC, RNA, or ADT-based analyses of the Level 1 (L1) TEAseq dataset (32,206 cells). **(B)** UMAP embeddings per modality colored by tissue. **(C)** Scaled average expression and percent expression of differentially expressed features between all 15 clusters resolved by each unimodal analysis. ATAC data shown per gene derive from GeneActivity inference of overall locus accessibility.

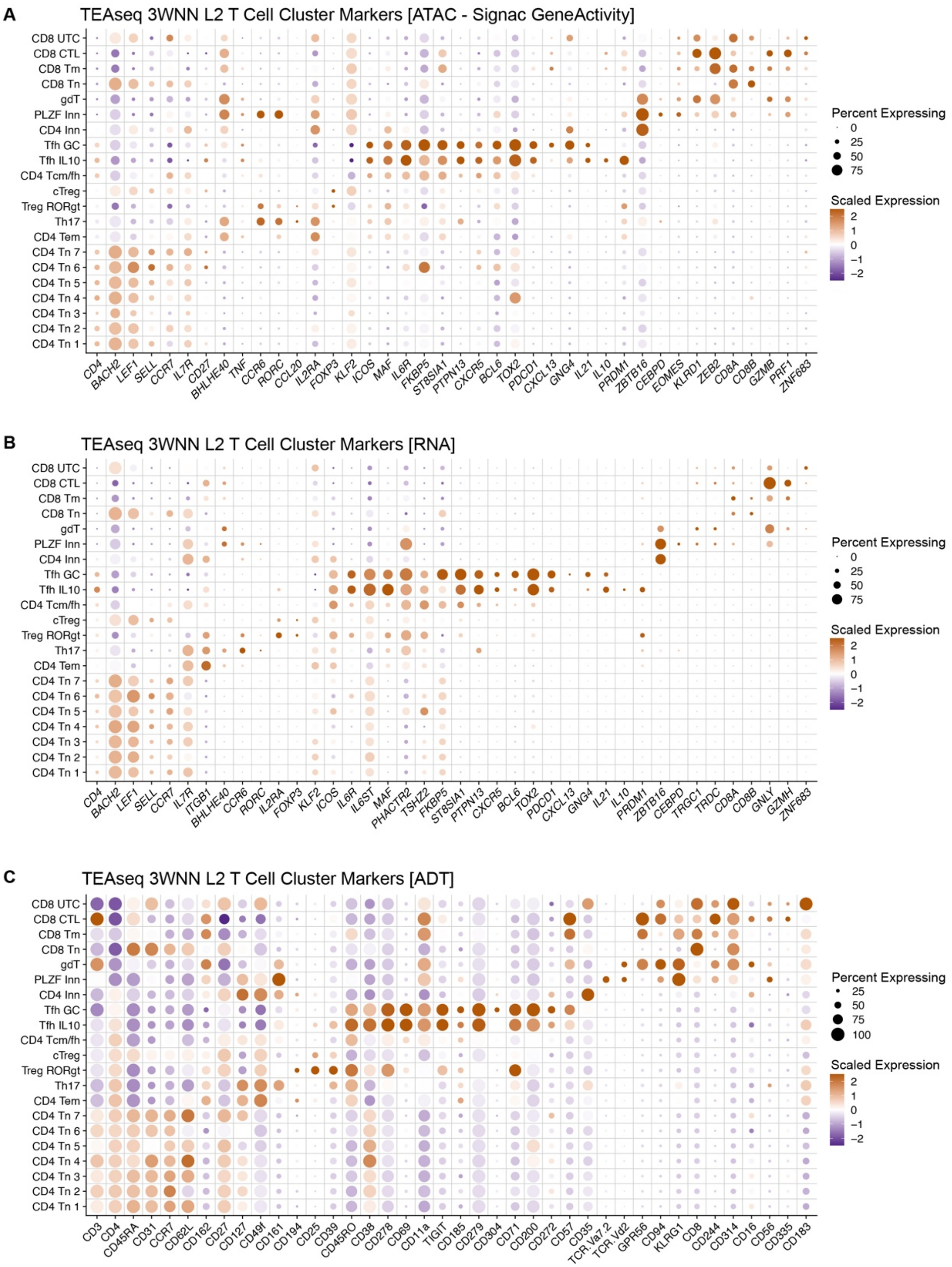


**Fig. S6 - Trimodal annotation of Level 2 Tfh versus nonTfh states within TEAseq dataset.** **(A-C)** Scaled average expression and percent expression of differentially expressed features between all 21 clusters resolved by three-way weighted-nearest neighbor (3WNN) analysis of the Level 2 (L2) TEAseq dataset (23,442 cells). Three modalities of gene expression are shown: **(A)** chromatin accessibility (from the ATAC-based GeneActivity assay), **(B)** mRNA transcripts, and **(C)** cell surface proteins (from antibody-derived tag (ADT) labeling).

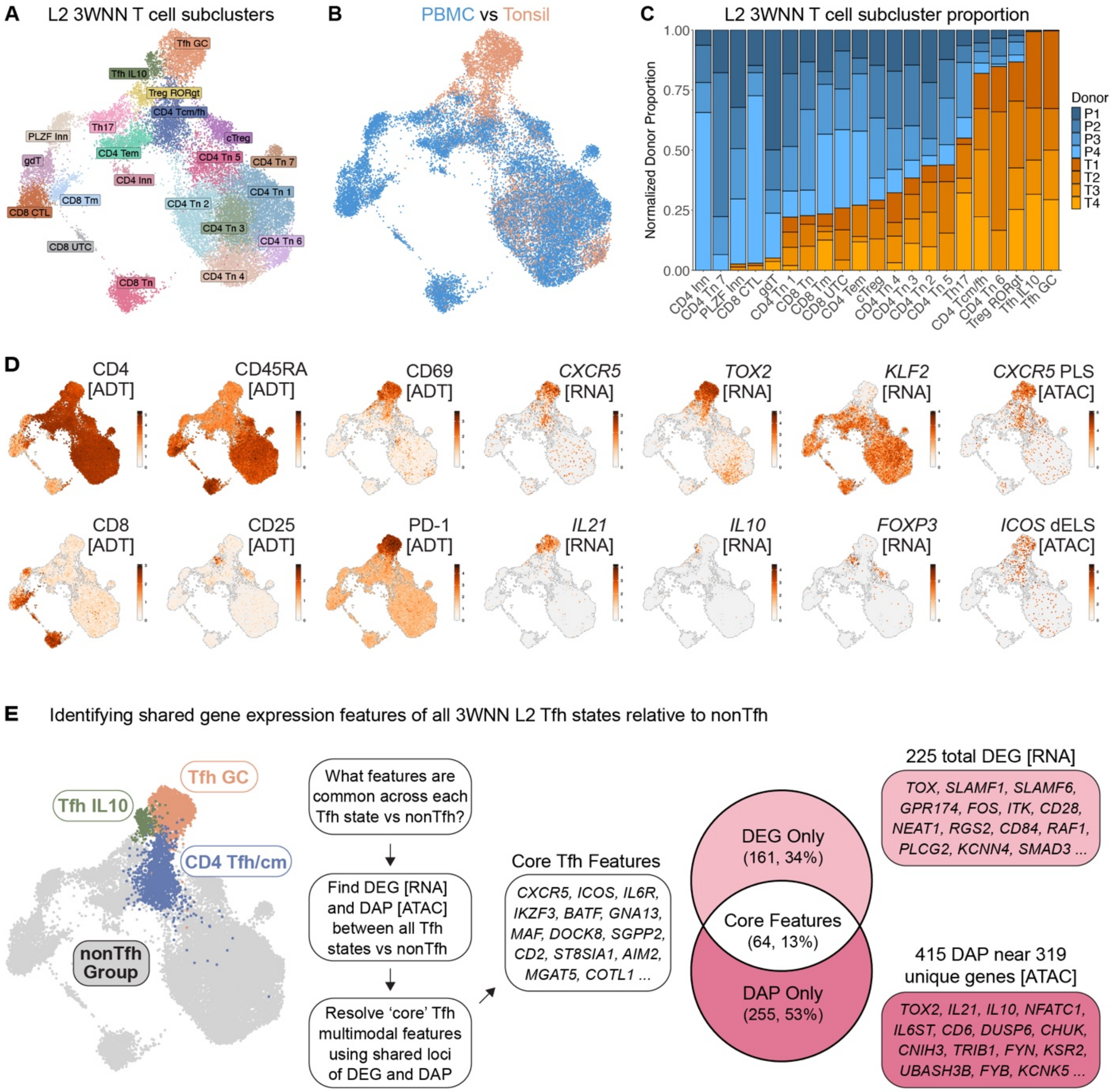


**Fig. S7 - Tfh adopt distinct states in tonsil and peripheral blood unified by a set of multimodal gene expression features. (A)** UMAP of 21 T cell clusters resolved by three-way weighted-nearest neighbor (3WNN) analysis of the Level 2 (L2) TEAseq dataset (23,442 cells). **(B)** 3WNN UMAP colored by tissue. **(C)** Stacked barplot of unenriched tonsil (T) versus PBMC (P) sample proportions per L2 T cell cluster, normalized to sample cell total. **(D)** 3WNN UMAP visualization of differentially expressed multimodal features between L2 T cell clusters. **(E)** Schematic illustrating analysis approach to identify differentially accessible peaks (DAP) and expressed genes (DEG) in each L2 Tfh cluster relative to all other clusters (nonTfh). Venn diagram shows 'core' Tfh features defined by overlap of Tfh-enriched DAPs and DEGs.

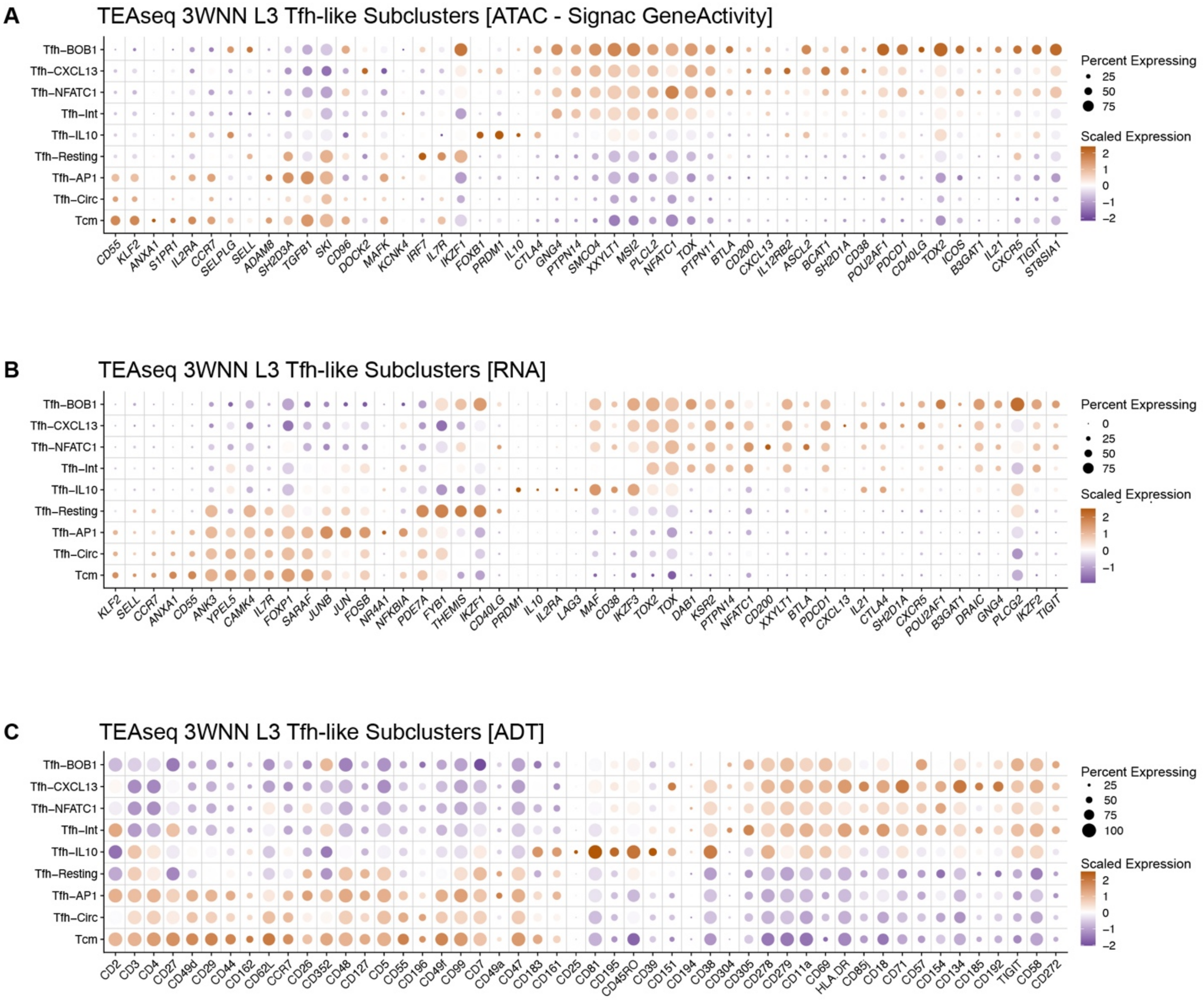


**Fig. S8 - Trimodal annotation of Level 3 Tfh versus Tcm states within TEAseq dataset.** **(A-C)** Scaled average expression and percent expression of differentially expressed features between all 9 clusters resolved by three-way weighted-nearest neighbor (3WNN) analysis of the Level 3 (L3) TEAseq dataset (3,657 cells). Three modalities of gene expression are shown: **(A)** chromatin accessibility (from ATAC-based GeneActivity analysis), **(B)** mRNA transcripts, and **(C)** cell surface proteins (from antibody-derived tag (ADT) labeling).

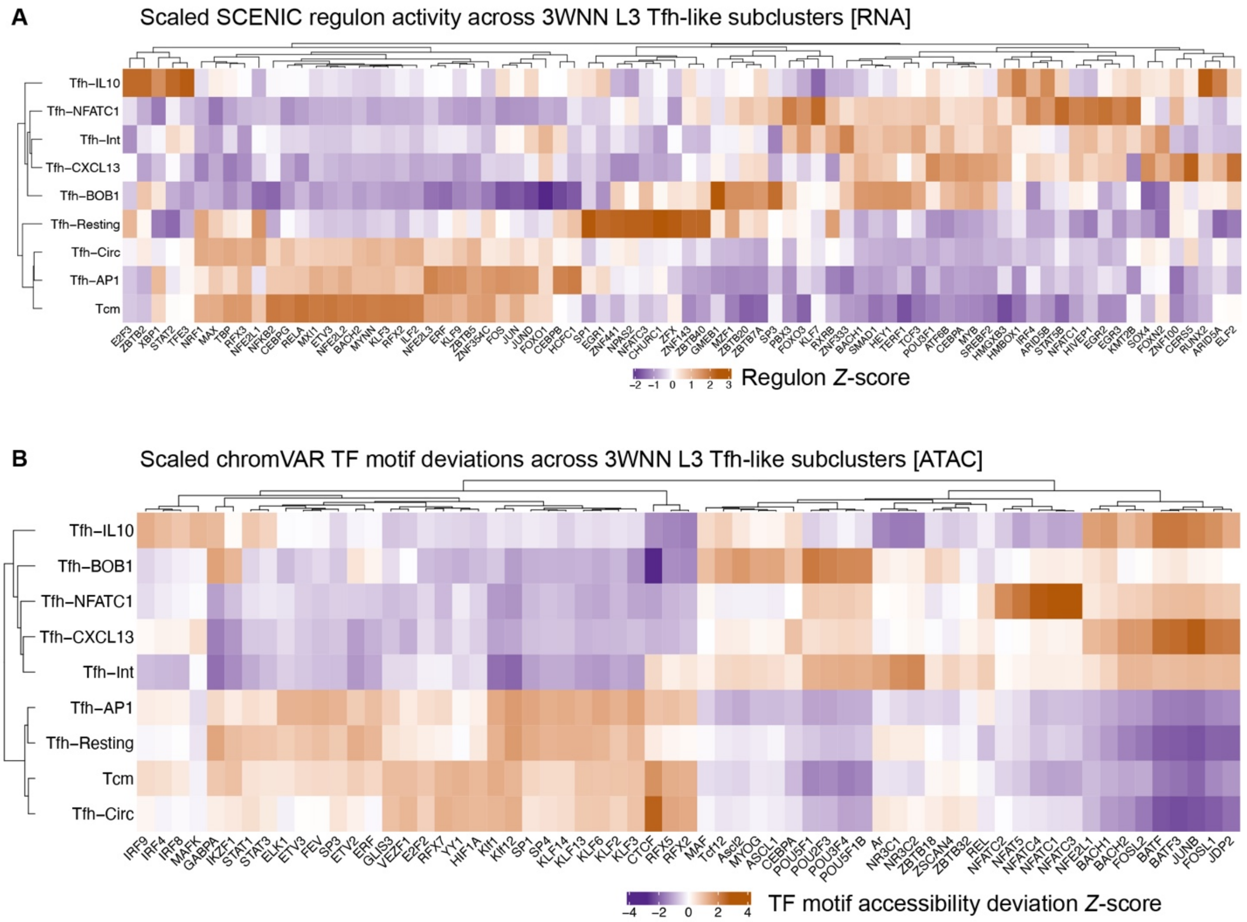


**Fig. S9 - SCENIC and chromVAR analyses infer distinct patterns of gene regulatory factor activity between Tfh states. (A)** Heatmap of transcription factor (TF) regulon activity inferred by SCENIC analysis, scaled across all nine Tfh-like and Tcm states resolved by three-way weighted-nearest neighbor (3WNN) analysis of the Level 3 (L3) TEAseq object (3,657 cells). **(B)** Heatmap of differential TF motif accessibility inferred by chromVAR analysis, scaled across L3 clusters. Clusters (rows) and TFs (columns) are hierarchically clustered for both heatmaps.

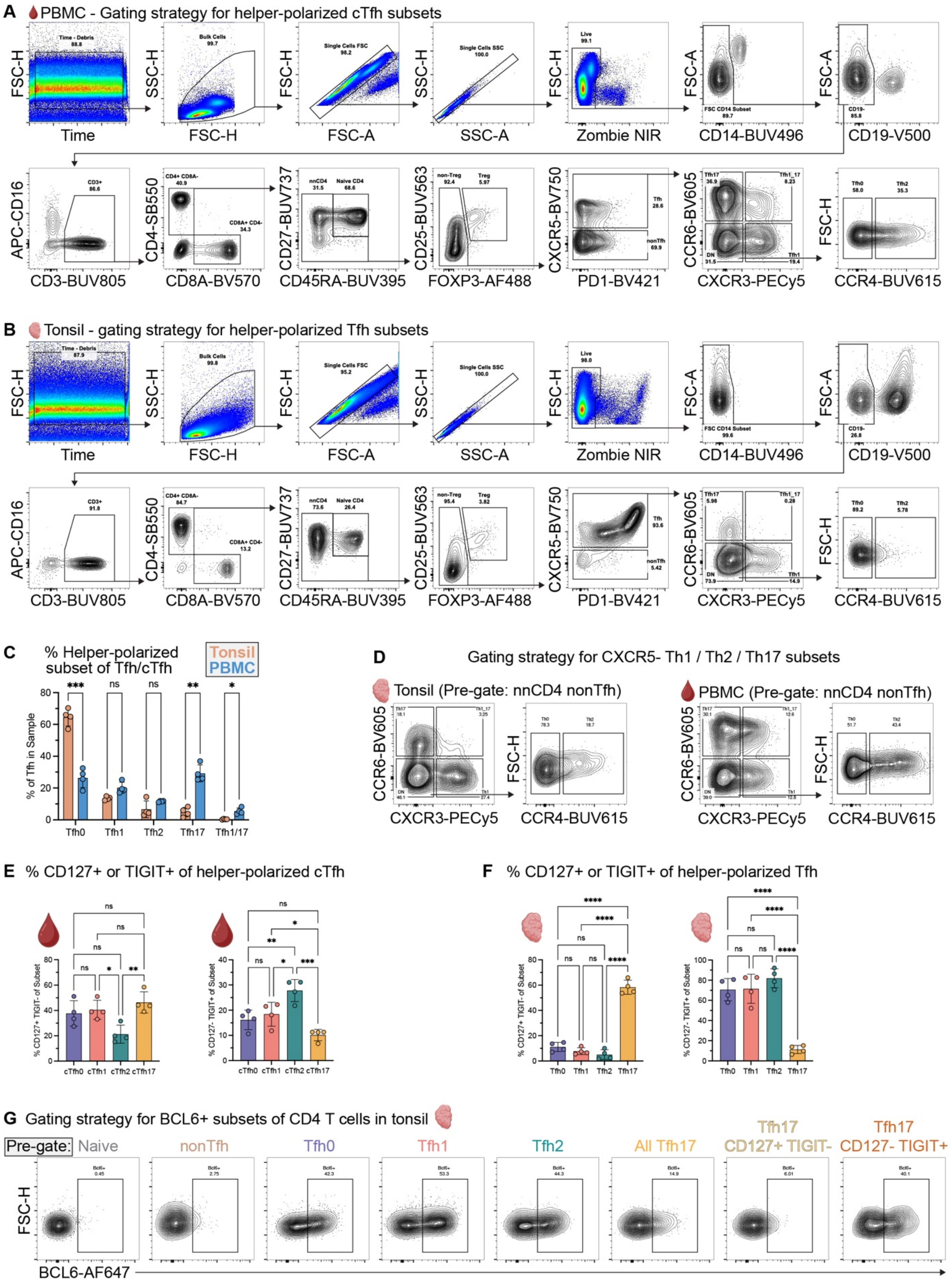


**Fig. S10 - Spectral flow cytometry gating strategy and phenotyping of helper-polarized Tfh subsets in peripheral blood and tonsil. (A-B)** Representative gating strategies for **(A)**

peripheral blood and **(B)** tonsil mononuclear cells profiled by 30-color spectral flow cytometry (*n* = 4 donors per tissue, unpaired). **(C)** Barplot showing percentage of Tfh corresponding to each polarized subset between tissues (unpaired *t*-test with Welch correction). **(D)** Representative gating strategy for $CXCR5^{-}$ T helper subsets. **(E-F)** Barplots showing percentage of each polarized Tfh subset that express TIGIT or CD127 in **(E)** peripheral blood and **(F)** tonsil (one-way ANOVA with Holm-Šídák's multiple comparisons test). **(G)** Representative gating strategy for BCL6 expression across polarized Tfh subsets and nonTfh. Statistics: $*P < 0.05$, $**P < 0.01$, $***P < 0.001$, $****P < 0.0001$.

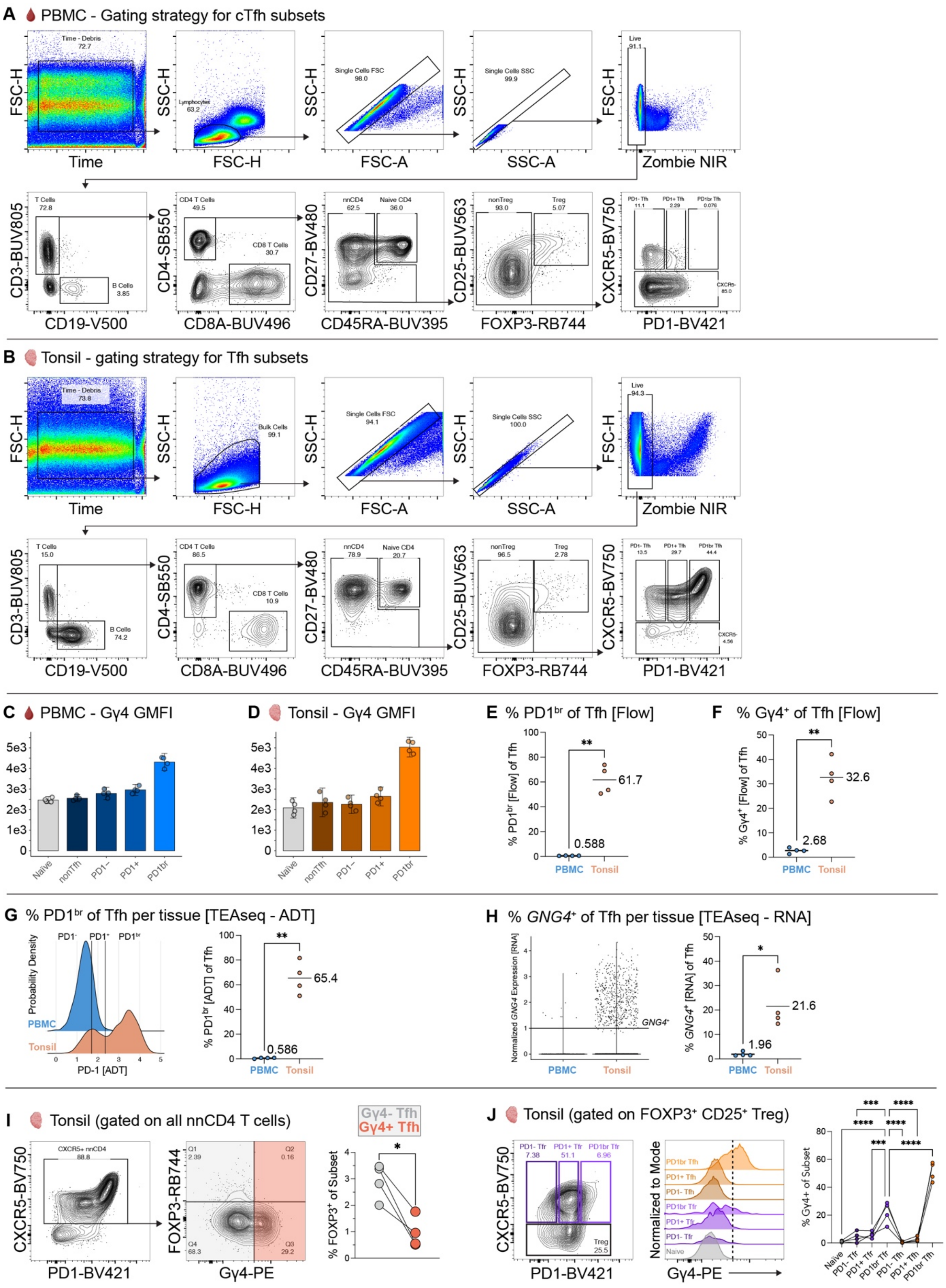


**Fig. S11 - Gating strategy and quantification of Gγ4 protein and *GNG4* RNA expression in human CD4 T cells. (A-B)** Representative gating strategy for naive, CXCR5⁻ nonTfh, and

CXCR5$^+$ Tfh subsets of CD4 T cells in **(A)** PBMC and **(B)** tonsil samples ($n$ = 4 donors per tissue, unpaired). **(C-D)** Barplots showing Gγ4 GMFI across Tfh subsets defined by PD1 expression versus nonTfh and naive CD4 T cells in **(C)** PBMC and **(D)** tonsil samples. Two-sided $t$-distribution 95% confidence intervals for each subset are shown. **(E)** Percentage of Tfh with bright PD1 (PD1$^{br}$) or **(F)** positive Gγ4 protein expression in tonsil versus PBMC samples (unpaired $t$-test with Welch correction). **(G)** Ridge plot shows gating of Tfh subsets based on PD1-ADT expression in Level 4 TEAseq analysis, with scatter plot showing the percentage of PD1$^{br}$ Tfh between tonsil and PBMC samples (unpaired $t$-test with Welch correction). **(H)** Violin plot defining threshold for positive *GNG4* RNA expression in Level 4 TEAseq Tfh cells, with scatter plot showing the percentage of *GNG4* RNA$^+$ Tfh in tonsil versus PBMC samples (unpaired $t$-test with Welch correction). **(I)** Representative gating strategy for FOXP3 versus Gγ4 protein expression in CXCR5$^+$ nnCD4 T cells from tonsil, with connected dot plot showing the percentage of FOXP3$^+$ cells in Gγ4$^+$ versus Gγ4$^-$ Tfh subsets per donor (paired $t$-test). **(J)** Representative gating strategy for CXCR5 versus PD1 protein expression in Treg cells from tonsil, with histogram showing Gγ4 protein expression between Tfh and Tfr subsets split by PD1 expression. Dashed line indicates threshold defining positive Gγ4 protein expression level relative to naive CD4 T cell internal negative control. Connected dot plot shows the percentage of Gγ4$^+$ cells across naive CD4 T cell, Treg, and Tfh subsets (one-way ANOVA with Holm-Šídák's correction for multiple comparisons). Statistics: *$P < 0.05$, **$P < 0.01$, ***$P < 0.001$, ****$P < 0.0001$.

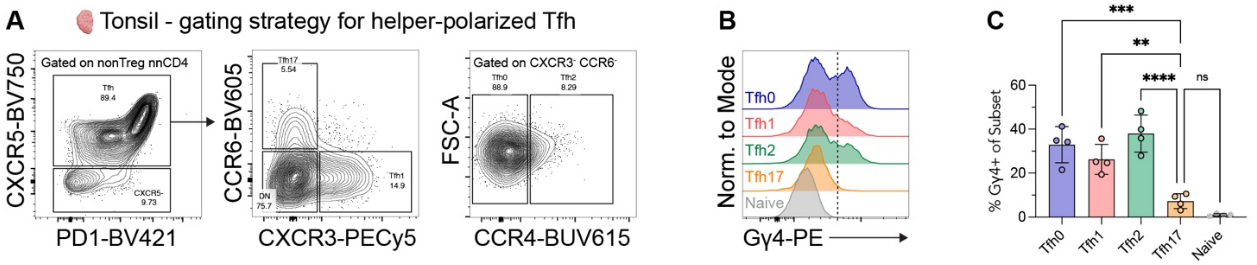


**Fig. S12 - Tfh17 express decreased Gγ4 protein relative to other helper-polarized subsets of Tfh in tonsil. (A)** Representative gating strategy used to define polarized subsets of Tfh in tonsil samples. **(B)** Histogram of Gγ4 protein expression between polarized Tfh subsets. Vertical dashed line indicates threshold defining positive Gγ4 protein expression level relative to naive CD4 T cell internal negative control. **(C)** Scatter plot shows percentage of Gγ4$^+$ cells in each polarized Tfh subset (one-way ANOVA with Holm-Šídák's correction for multiple comparisons). Statistics: **$P < 0.01$, ***$P < 0.001$, ****$P < 0.0001$.

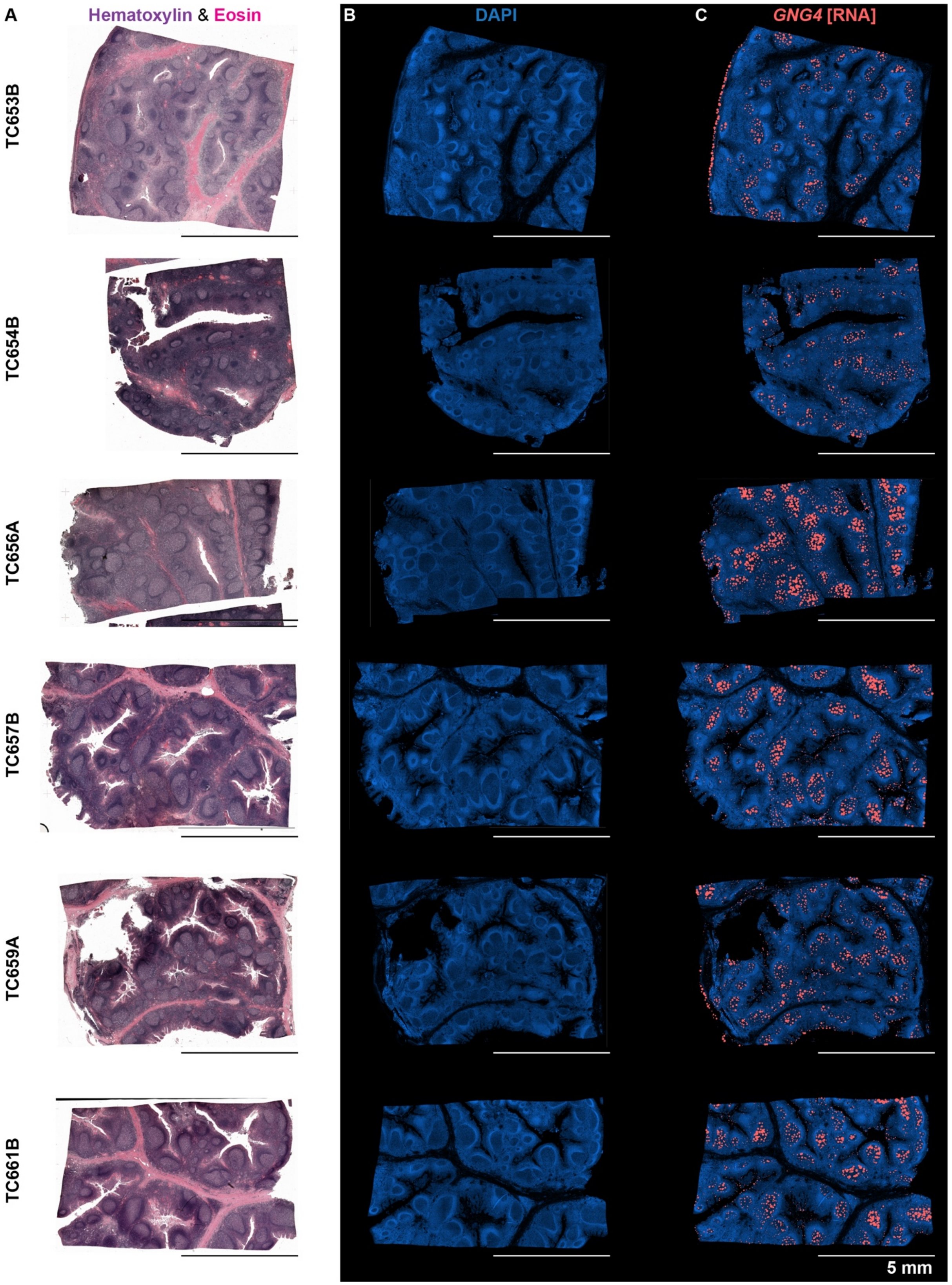


**Fig. S13 - Anatomic resolution of *GNG4* transcripts within germinal center and epithelial regions of tonsil tissue. (A)** Aligned whole-slide images of hematoxylin and eosin staining for all six tonsil samples analyzed using the Xenium Prime single-cell spatial transcriptomics assay. **(B)** DAPI images from the Xenium Multi-Tissue Stain per sample. **(C)** Scaled *GNG4* RNA probe signal (red circles) overlaid on DAPI image for each sample.

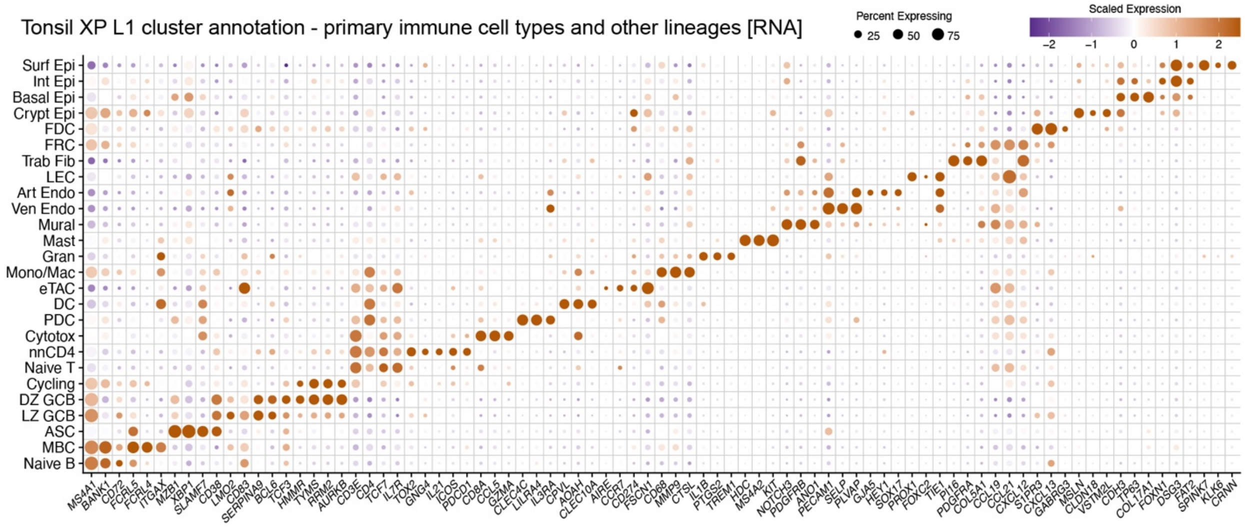


**Fig. S14 - Resolution of diverse immune and non-immune cell types in tonsil single-cell spatial transcriptomics. (A)** Percent expression and scaled average expression values for differentially expressed RNA features between all 26 manually annotated cell types from Level 1 clustering analysis of the Xenium Prime (XP) dataset (6,290,476 cells total).

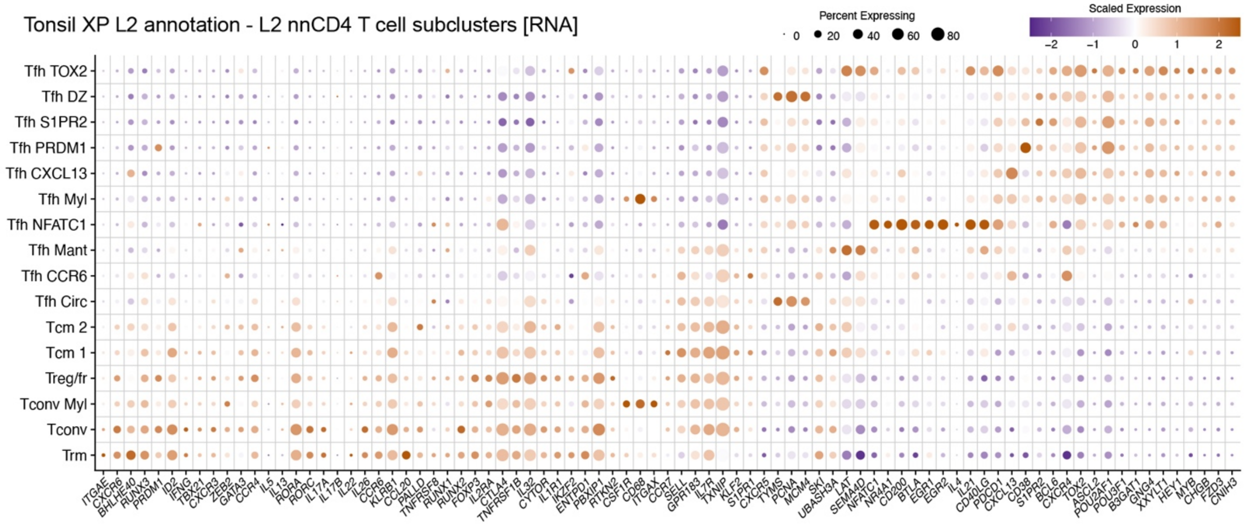


**Fig. S15 – Level 2 subclustering analysis of nnCD4 T cells in tonsil Xenium Prime dataset reveals distinct Tfh versus nonTfh states.** Percent expression and scaled average expression values for differentially expressed RNA features between all 16 Level 2 (L2) cell types resolved by subclustering analysis of nnCD4 T cells (330,994 total) from the L1 Xenium Prime dataset.

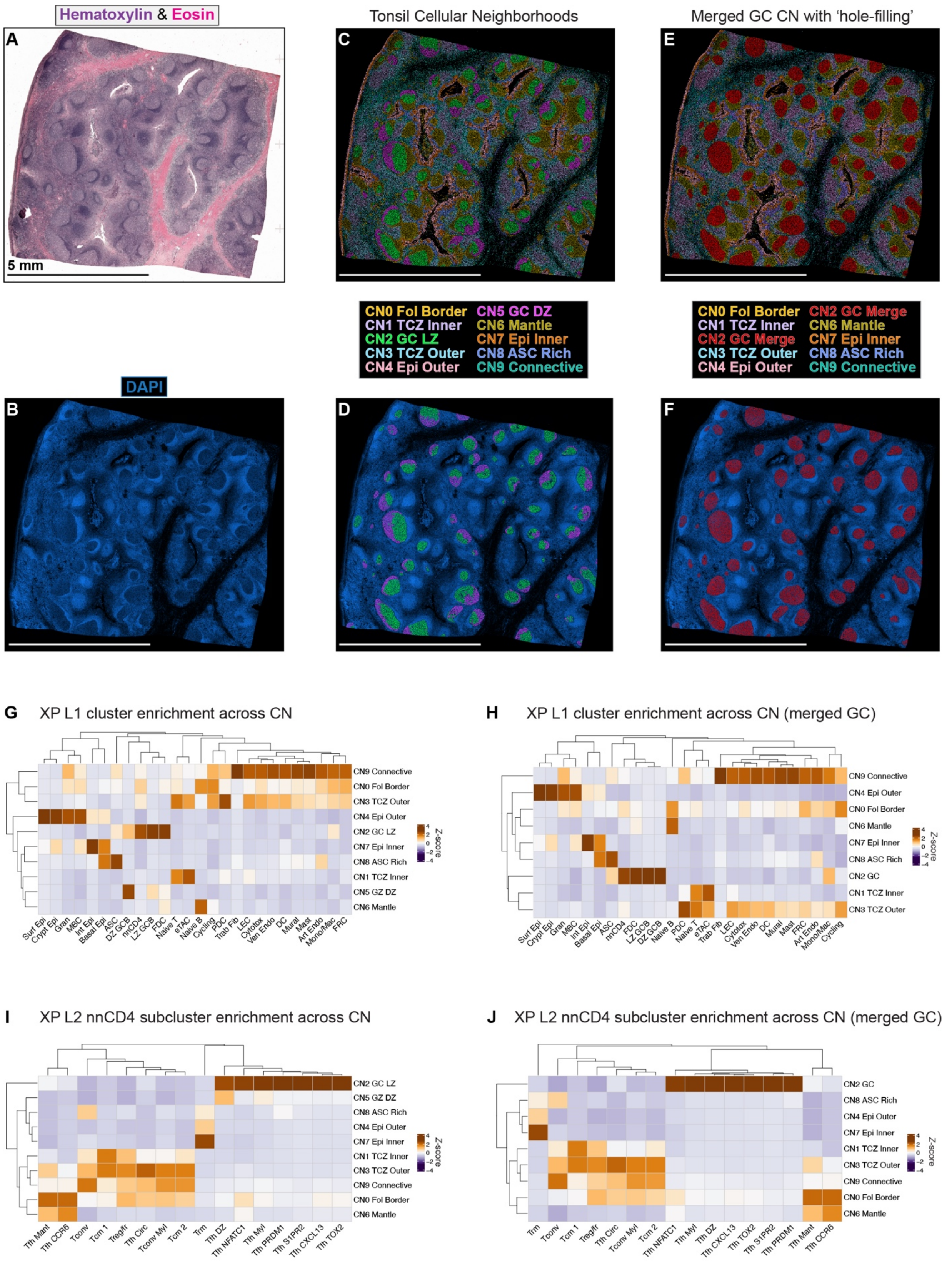


**Fig. S16 - Immune and non-immune cell types form diverse cellular neighborhoods in tonsils with distinct CD4 T cell subset composition. (A)** Representative H&E and **(B)** DAPI images from tonsil sample TC653B. **(C)** Segmented cells colored by annotated cellular neighborhood (CN). **(D)** CN annotations overlaid on DAPI image. **(E)** Segmented cells colored by CN annotation in the merged GC CN analysis. **(F)** Merged GC CN annotations overlaid on DAPI image. **(G-J)** Heatmaps with hierarchically clustered rows (CN) and columns (cluster cell types) showing *Z*-scored cell counts per cluster across CN: **(G)** Level 1 (L1) cell types across CNs, **(H)** L1 cell types across CNs in merged GC analysis, **(I)** L2 nnCD4 T cell subclusters across CNs, **(J)** L2 nnCD4 T cell subclusters across CNs in merged GC analysis.

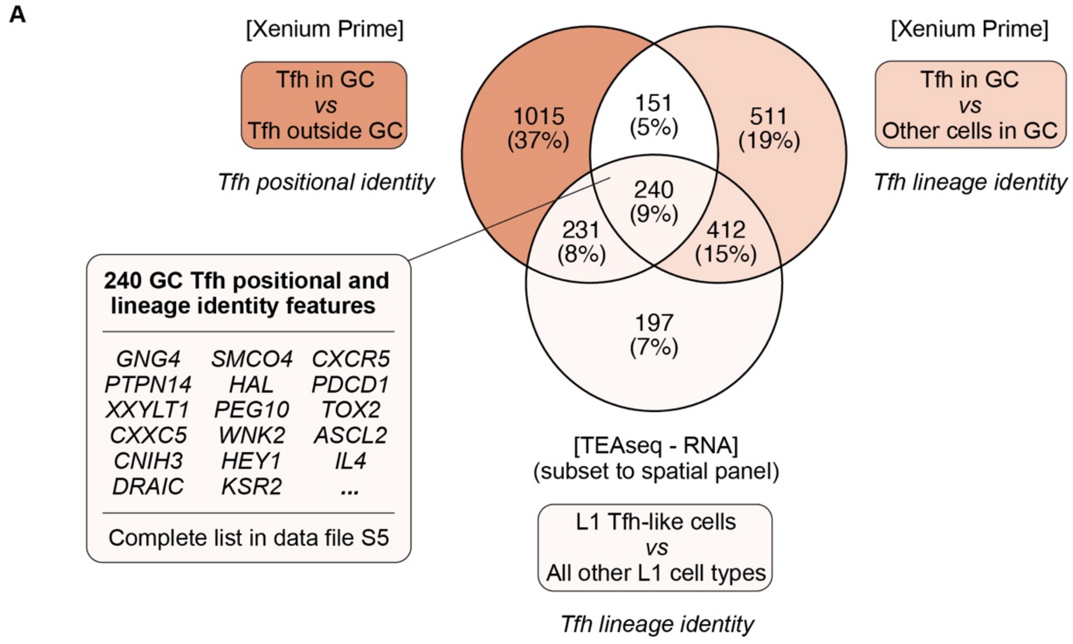


**Fig. S17 - Combined spatial transcriptomic and TEAseq analyses resolve a core set of GC Tfh positional and lineage identity features. (A)** Venn diagram showing the intersection of differentially expressed RNA features resolved by three different comparisons: (1) GC versus nonGC Tfh [Xenium Prime], (2) Tfh versus all other cell types within the GC [Xenium Prime], and (3) Tfh-like cells versus all other mononuclear cell subsets defined in trimodal clustering analysis of the Level 1 TEAseq dataset. All features were filtered based on positive average $\log_2$FC and $P < 0.05$ using a Wilcoxon rank-sum test with Bonferroni correction. Genes measured by the whole-transcriptome TEAseq assay but not Xenium Prime were excluded.

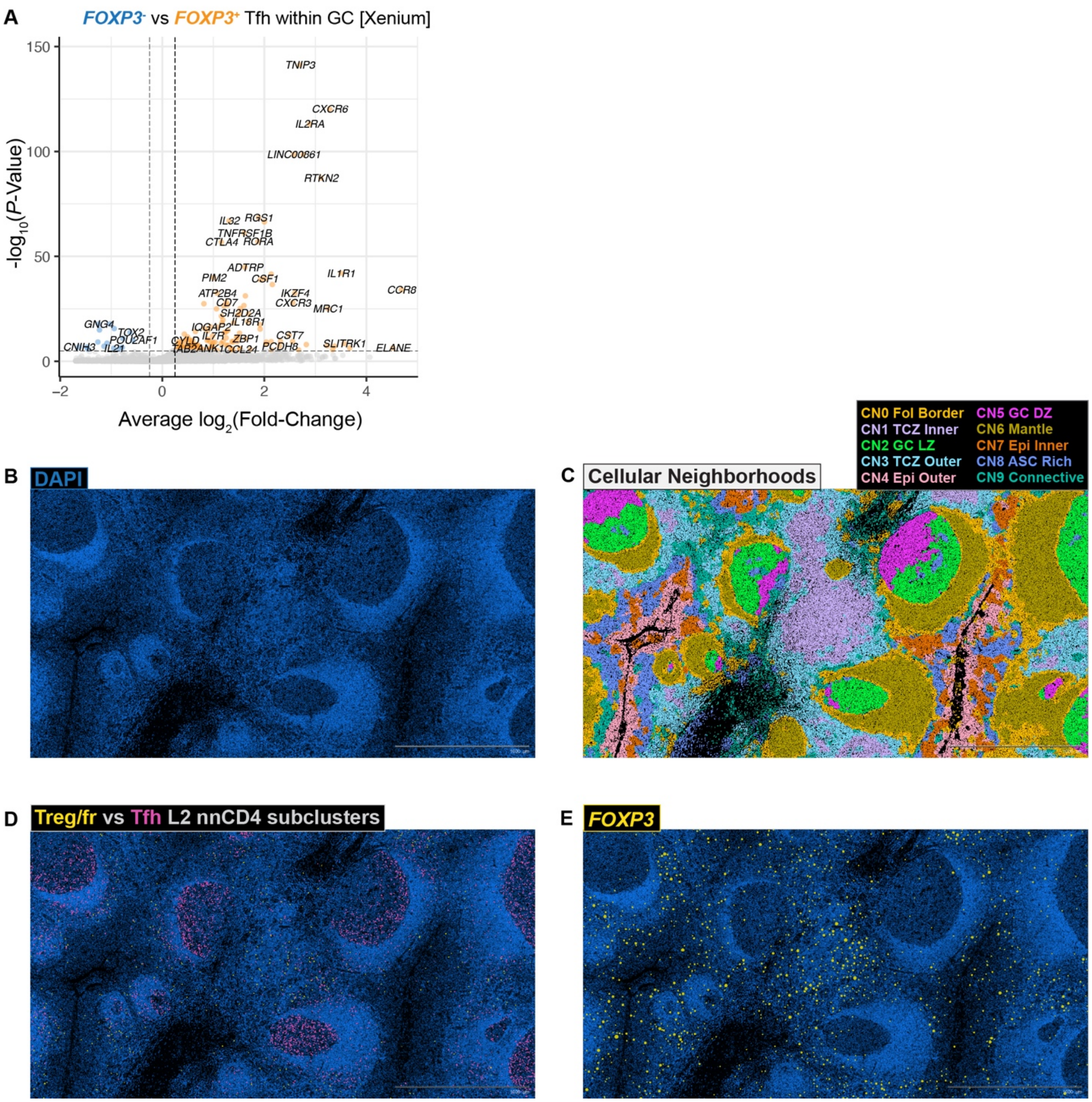


**Fig. S18 - *FOXP3*+ Tfh within the GC exhibit Treg-like transcriptional features and reduced *GNG4* expression. (A)** Volcano plot showing differentially expressed genes (DEG) between *FOXP3* RNA$^+$ versus RNA$^-$ Tfh positioned within merged GC cellular neighborhoods. DEG thresholds were defined as $|\log_2 FC| > 0.25$ and $P < 1e\text{-}5$ by Wilcoxon rank-sum test. **(B-E)** Representative magnified field of view from tonsil TC653B. **(B)** DAPI nuclear staining. **(C)** Segmented cells colored by cellular neighborhood (CN) annotations. **(D)** Spatial positioning of Treg/fr and all Tfh cell types resolved in Level 2 clustering analysis of Xenium Prime data overlaid on DAPI image. **(E)** Scaled *FOXP3* RNA probe signal overlaid on DAPI image.

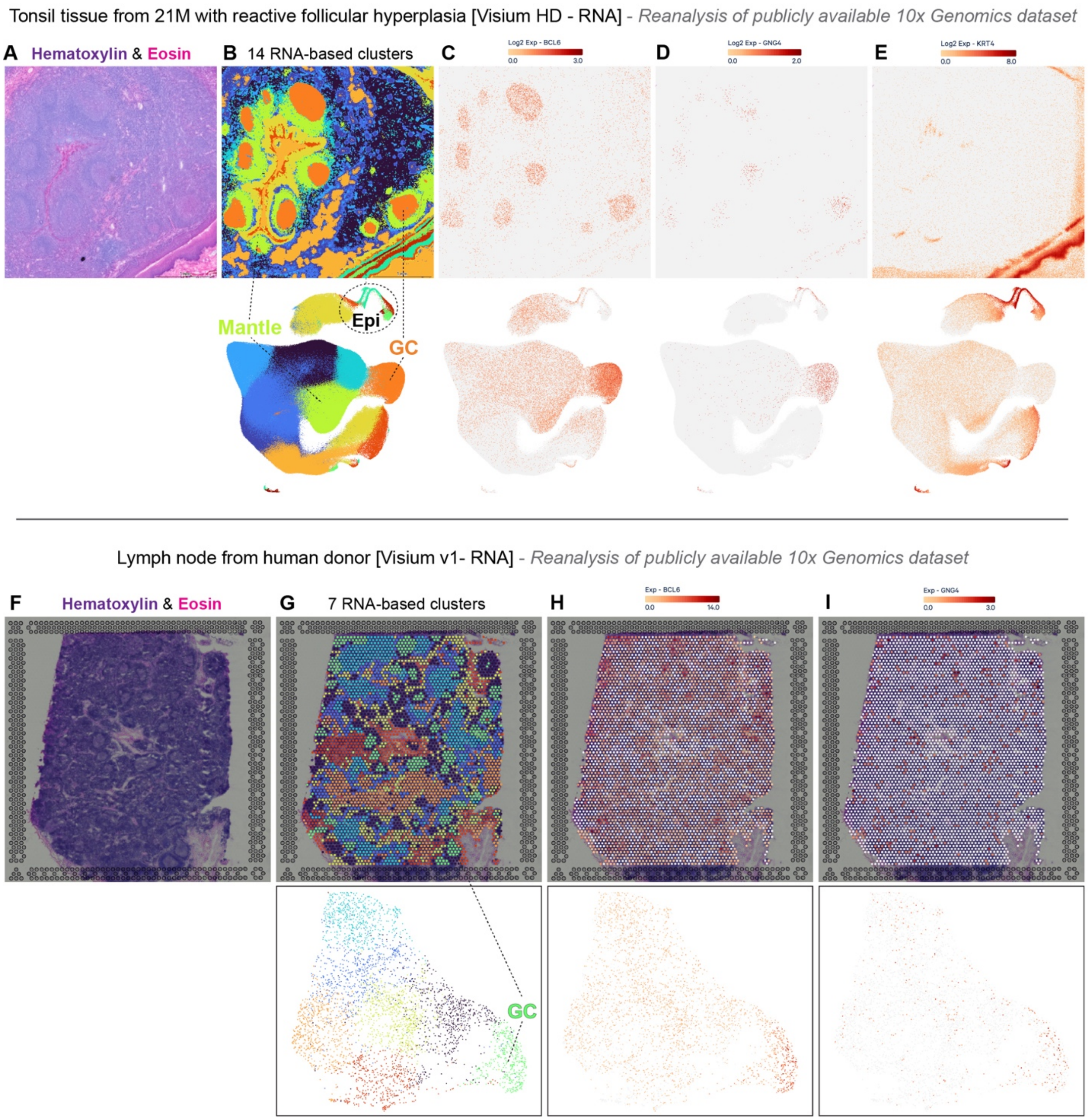


**Fig. S19 - *GNG4* expression is spatially enriched within GC of adult tonsil and lymph node tissue as well as epithelial cells.** Publicly available **(A-E)** tonsil Visium HD and **(F-I)** lymph node Visium v1 data from 10x Genomics were reanalyzed to assess spatial patterning of *GNG4* expression **(A)** H&E image of tonsil sample. **(B)** UMAP embedding showing several annotated graph-based clusters and normalized expression of **(C)** *BCL6* **(D)** *GNG4* and **(E)** *KRT4* RNA. **(F)** H&E image of lymph node sample. **(G)** UMAP embedding showing several annotated graph-based clusters and normalized expression of **(H)** *BCL6* and **(I)** *GNG4* transcripts.

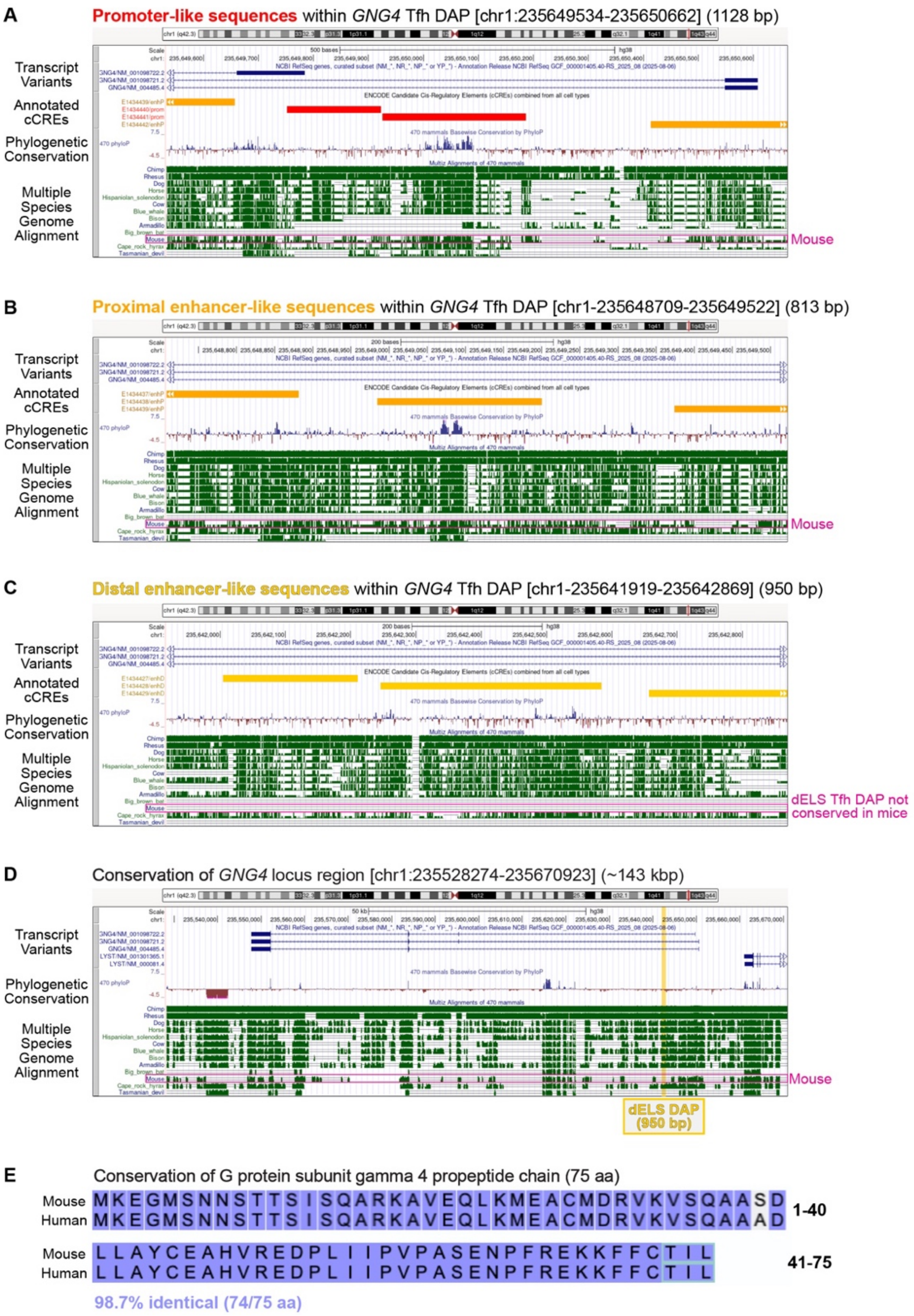


**Fig. S20 - Annotation and species conservation of *GNG4* regions with enriched accessibility in human GC-like Tfh states. (A-D)** UCSC Genome Browser (*140*) visualization of the human *GNG4* locus including annotations of candidate *cis*-regulatory elements (cCRE) from ENCODE (*68*) as well as species conservation based on PhyloP scores and Multiz multiple alignment. Transcript variants derive from the NCBI RefSeq database. **(A-C)** Visualization of *GNG4* regions defined as enriched ATAC peaks in GC-like Tfh states as well as **(D)** the entire *GNG4* locus and surrounding annotated genes. **(E)** Alignment of Gγ4 polypeptide sequence between *Mus musculus* and *Homo sapiens* (*191*). Teal box indicates cleaved C-terminal residues.

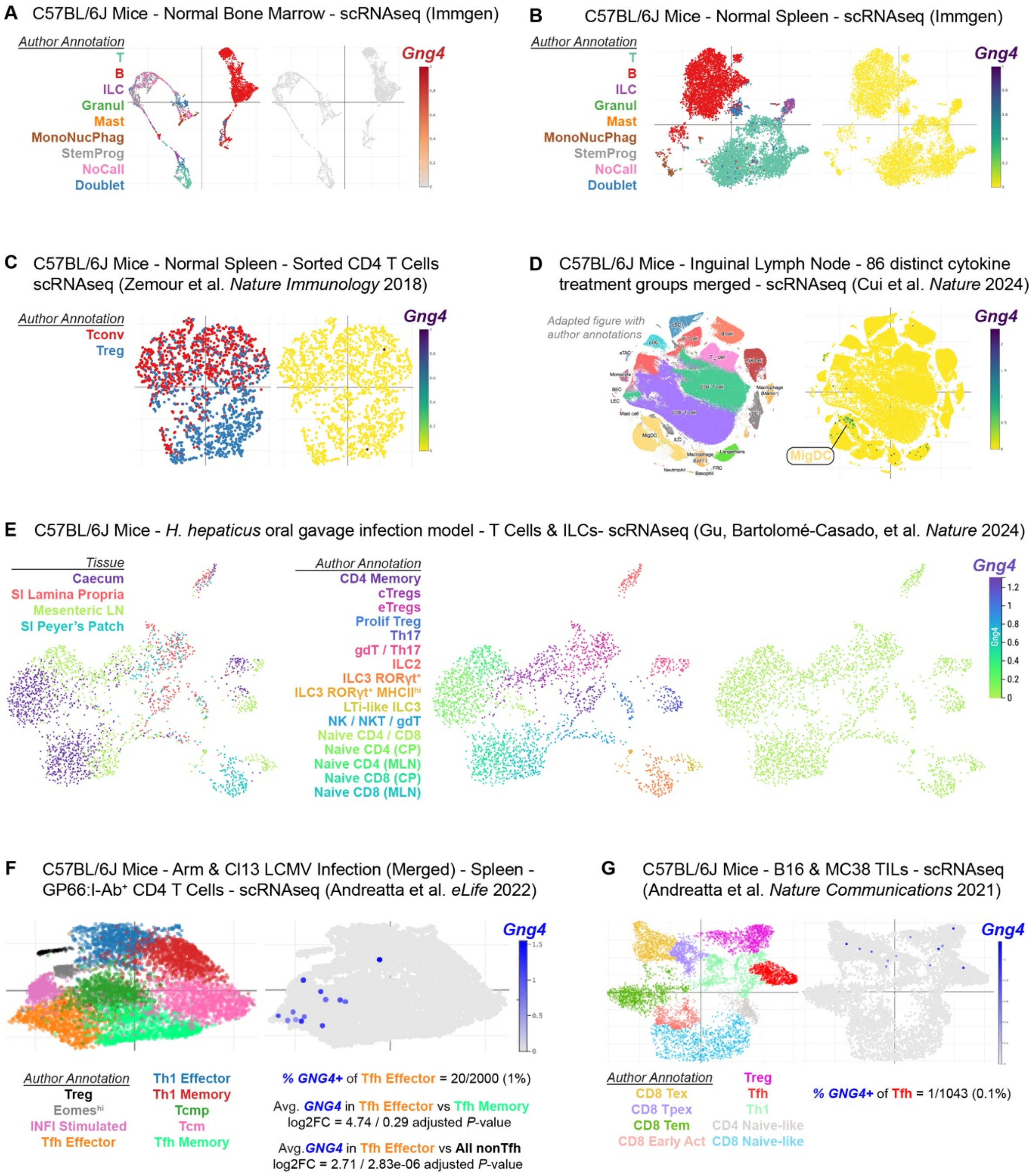


**Fig. S21 - Sparse *Gng4* RNA expression in T cells from C57BL/6J laboratory mice across diverse health and disease states. (A-G)** Dimensionally reduced embeddings of scRNAseq data from mice for reanalysis of *Gng4* expression across multiple tissues and contexts: **(A)** bone marrow (*146*), **(B)** spleen (*146*), **(C)** splenic CD4 T cells (*145*), **(D)** inguinal lymph node including 86 distinct cytokine treatment groups (*143*), **(E)** T and ILC subsets from GI tissues during *H. hepaticus* infection (*144*), **(F)** LCMV-specific CD4 T cells in spleen from both acute and chronic LCMV models (*142*), and **(G)** tumor-infiltrating lymphocyte (TIL) subsets from both B16 and MC38 tumor models (*141*). Results of differential RNA expression tests are shown in Panel F for indicated populations (Wilcoxon rank-sum test with Bonferroni correction).

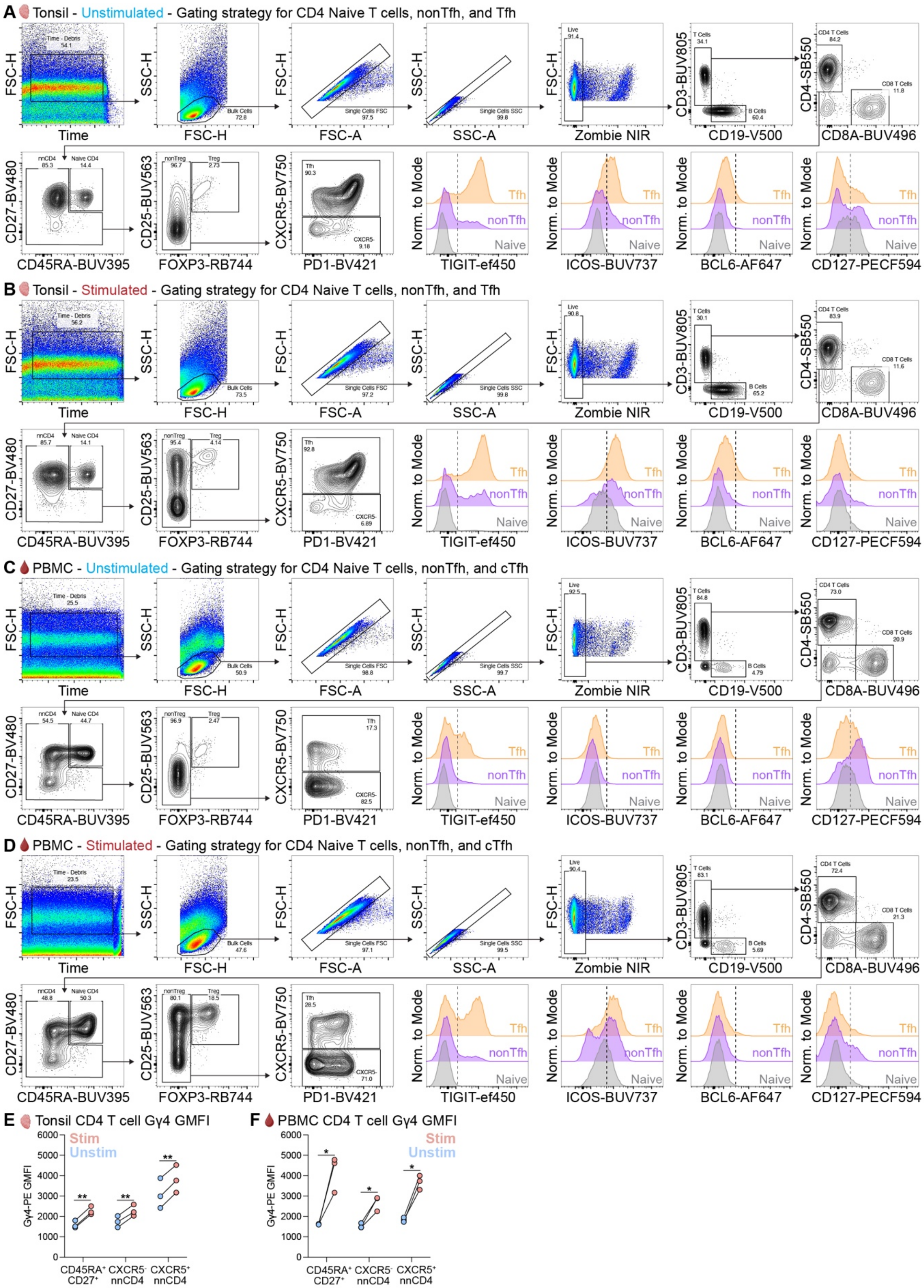


**Fig. S22 - Spectral flow cytometry gating strategy for *in vitro* stimulation of mononuclear cells and Gγ4 expression. (A-D)** Representative gating strategy for **(A-B)** tonsil and **(C-D)**

PBMC samples in unstimulated versus anti-CD3/CD28/CD49d conditions to define $CD45RA^{+}CD27^{+}$ CD4, $CXCR5^{-}$ nnCD4, and $CXCR5^{+}$ nnCD4 T cell subsets. Histograms show TIGIT, ICOS, BCL6, and CD127 expression, with vertical dashed line indicating threshold for comparison of unstimulated versus stimulated conditions. **(E)** Connected dot plot shows Gγ4 GMFI in each CD4 T cell subset between conditions for tonsil and **(F)** PBMC samples (paired *t*-test with two-stage step-up Benjamini, Krieger, and Yekutieli FDR threshold of 5%. *$Q < 0.05$, **$Q < 0.01$. $n = 3$ donors per tissue, paired between conditions).

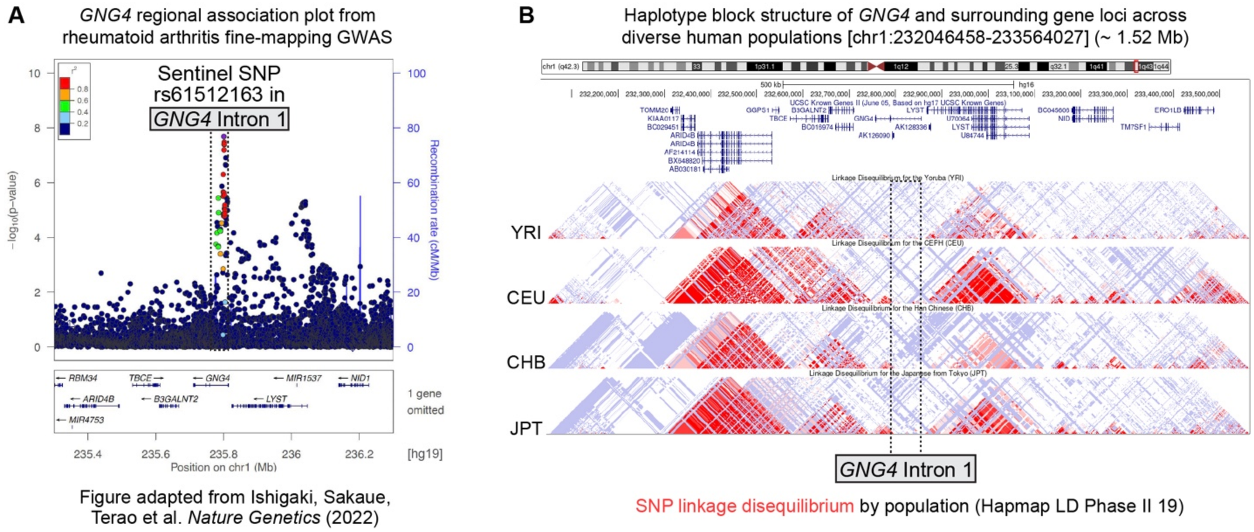


**Fig. S23 - Rheumatoid arthritis risk variants of *GNG4* map to the first intron between haplotype blocks. (A)** Adapted LocusZoom visualization from rheumatoid arthritis GWAS (*157*) showing fine-mapping of risk variants to the first intronic region of *GNG4*. **(B)** Visualization of SNP linkage disequilibrium surrounding the *GNG4* locus across diverse populations in the UCSC Genome Browser (*140*). The first intronic region of *GNG4* is indicated by dashed black lines (YRI, Yoruba in Ibadan, Nigeria; CEU, Utah residents with Northern and Western European ancestry; CHB, Han Chinese in Beijing, China; JPT, Japanese in Tokyo, Japan).

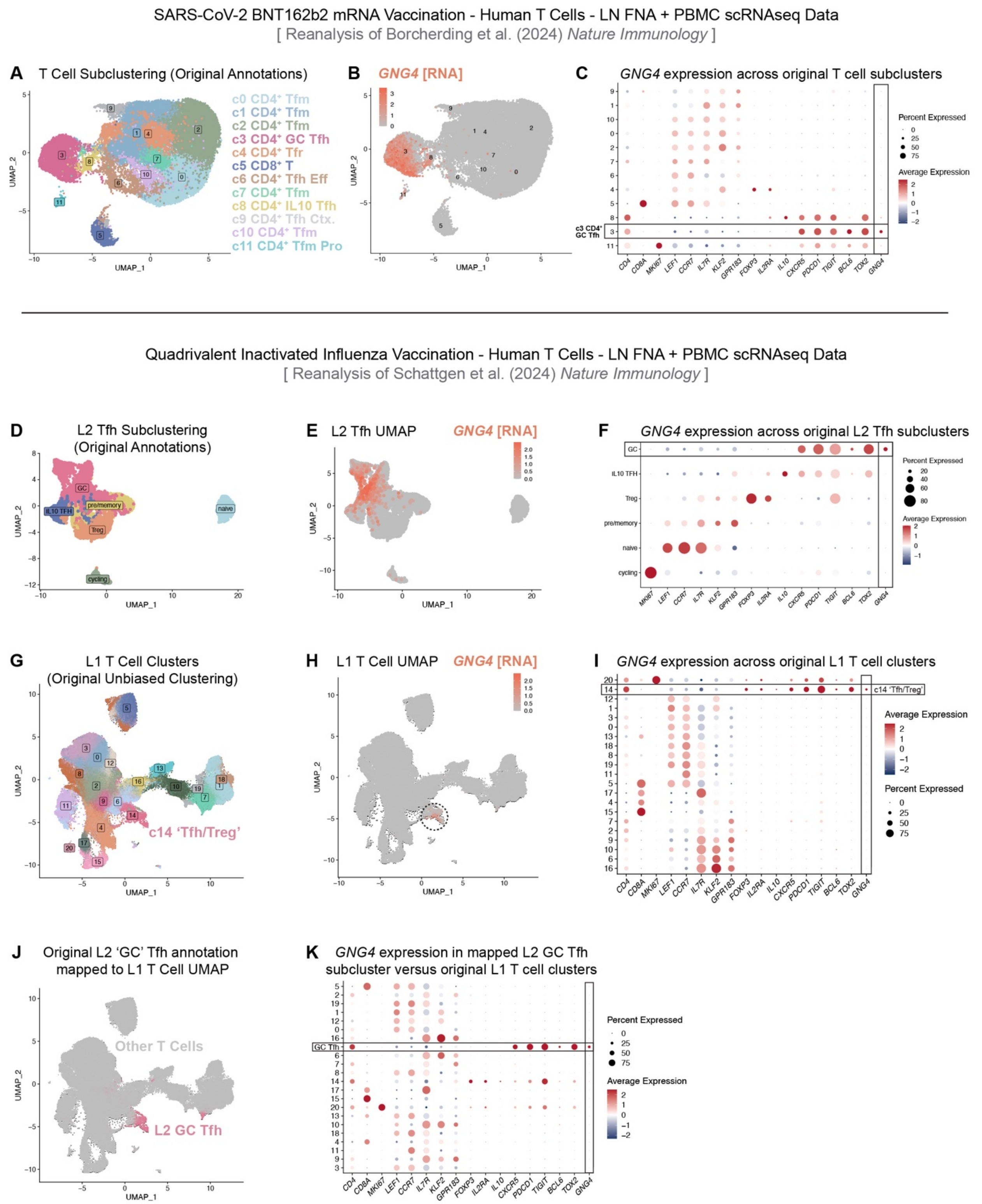


**Fig. S24 - *GNG4* expression is specifically induced in human GC-like Tfh during both SARS-CoV-2 and influenza vaccine responses. (A-C)** Reanalysis of scRNAseq data from longitudinally sampled axillary lymph nodes following vaccination with BNT162b2 mRNA against SARS-CoV-2 (*24*). **(A)** UMAP embedding of CD4 and CD8 T cell clusters in lymph node annotated by Borcherding *et al.* with **(B)** normalized expression of *GNG4* overlaid. **(C)** Percent and mean expression of T cell subset signature genes as well as *GNG4* across annotated cell types (cluster 3 = 'GC Tfh'). **(D-K)** Reanalysis of scRNAseq data from longitudinally sampled axillary lymph nodes following quadrivalent inactivated influenza vaccination (IIV) (*25*). **(D)** UMAP embedding of Level 2 (L2) Tfh subclusters annotated by Schattgen *et al.* with **(E)** normalized expression of *GNG4* overlaid. **(F)** Percent and average expression of differentially expressed genes between T cell states, including *GNG4*. **(G)** Level 1 (L1) UMAP and clustering of CD4 and CD8 T cell types in lymph node IIV study with **(H)** overlaid *GNG4* RNA expression. **(I)** Percent and average expression of differentially expressed features across L1 T cell subsets. **(J)** Cell barcodes corresponding to 'GC Tfh' in Schattgen *et al.* L2 Tfh subclustering analysis were mapped back to the L1 T cell object and UMAP embedding to enable contrast with cells not included in the L2 analysis. **(K)** Percent and mean expression of signature features between CD4 and CD8 T cell subsets, including *GNG4*.

#### Supplementary Tables

**Table S1.** Human PBMC and tonsil donor demographic information.

| Cohort for TEAseq and 30-color flow cytometry data (Panel A for cell sorting, Panel B for Fig. 2 flow data, antibodies listed in Table S5) | | | | |
|---|---|---|---|---|
| Tonsil ID | ASAB | Age | Tonsillectomy Indication | Source |
| TC572 | F | 5 | Sleep-disordered breathing | Discarded surgical tissue (CHOP Non-IRB) |
| TC574 | F | 7 | Sleep-disordered breathing | Discarded surgical tissue (CHOP Non-IRB) |
| TC569 | M | 6 | Sleep-disordered breathing | Discarded surgical tissue (CHOP Non-IRB) |
| TC570 | M | 7 | Obstructive sleep apnea | Discarded surgical tissue (CHOP Non-IRB) |
| PBMC ID | ASAB | Age | Donor Type | Source |
| 15920-4 | F | 6 | Healthy donor | Sarah E. Henrickson Lab, CHOP IRB #15920 |
| 15920-72 | F | 4 | Healthy donor | Sarah E. Henrickson Lab, CHOP IRB #15920 |
| 15920-75 | M | 7 | Healthy donor | Sarah E. Henrickson Lab, CHOP IRB #15920 |
| 15920-124 | M | 7 | Healthy donor | Sarah E. Henrickson Lab, CHOP IRB #15920 |

| Cohort for Gγ4 protein expression phenotyping in Tfh by spectral flow cytometry (Panel C antibodies listed in Table S5, data in Fig. 3) | | | | |
|---|---|---|---|---|
| Tonsil ID | ASAB | Age | Tonsillectomy Indication | Source |
| TON1 | 2 | F | Oropharyngael dysphasia | Neil Romberg Lab, discarded surgical tissue (CHOP Non-IRB) |
| TON2 | 4 | M | Obstructive sleep apnea | Neil Romberg Lab, discarded surgical tissue (CHOP Non-IRB) |
| TON3 | 13 | F | Obstructive sleep apnea | Neil Romberg Lab, discarded surgical tissue (CHOP Non-IRB) |
| TC574 | 7 | F | Sleep-disordered breathing | Neil Romberg Lab, discarded surgical tissue (CHOP Non-IRB) |
| PBMC ID | ASAB | Age | Donor Type | Source |
| ND584 | F | 26 | Healthy Donor | University of Pennsylvania Human Immunology Core |
| ND647 | M | 31 | Healthy Donor | University of Pennsylvania Human Immunology Core |
| ND655 | F | 26 | Healthy Donor | University of Pennsylvania Human Immunology Core |
| ND661 | F | 51 | Healthy Donor | University of Pennsylvania Human Immunology Core |

| Cohort for palatine tonsil Xenium Prime single-cell spatial transcriptomics (data shown in Fig. 4) | | | | |
|---|---|---|---|---|
| Tonsil | ASAB | Age | Tonsillectomy Indication | Source |
| TC653 | M | 6 | Sleep-disordered breathing | Neil Romberg Lab, discarded surgical tissue (CHOP Non-IRB) |
| TC654 | F | 8 | Sleep-disordered breathing | Neil Romberg Lab, discarded surgical tissue (CHOP Non-IRB) |
| TC656 | M | 4 | Obstructive sleep apnea | Neil Romberg Lab, discarded surgical tissue (CHOP Non-IRB) |
| TC657 | F | 8 | Sleep-disordered breathing | Neil Romberg Lab, discarded surgical tissue (CHOP Non-IRB) |
| TC659 | M | 4 | Sleep-disordered breathing | Neil Romberg Lab, discarded surgical tissue (CHOP Non-IRB) |
| TC661 | F | 8 | Sleep-disordered breathing | Neil Romberg Lab, discarded surgical tissue (CHOP Non-IRB) |

| Cohort for mononuclear cell *in vitro* stimulation assay and spectral flow cytometry (Panel D antibodies listed in table S5, data in Fig. 5) | | | | |
|---|---|---|---|---|
| Tonsil ID | ASAB | Age | Tonsillectomy Indication | Source |
| TC564 | F | 6 | Obstructive sleep apnea | Neil Romberg Lab, discarded surgical tissue (CHOP Non-IRB) |
| TC579 | F | 5 | Obstructive sleep apnea | Neil Romberg Lab, discarded surgical tissue (CHOP Non-IRB) |
| TC638 | M | 7 | Obstructive sleep apnea | Neil Romberg Lab, discarded surgical tissue (CHOP Non-IRB) |
| PBMC ID | ASAB | Age | Donor Type | Source |
| ND639 | F | 33 | Healthy donor | University of Pennsylvania Human Immunology Core |
| ND651 | F | 25 | Healthy donor | University of Pennsylvania Human Immunology Core |
| ND661 | F | 51 | Healthy donor | University of Pennsylvania Human Immunology Core |

**Table S2.** TotalSeqA HTO and ADT antibodies used for TEAseq experiment.

| HTO and ADT costained with FACS antibodies before TEAseq | | | | | | | | | | | | | |
|---|---|---|---|---|---|---|---|---|---|---|---|---|---|
| TotalSeqA ID | Target | TotalSeq Barcode | Clone | Vendor | Catalog # | Lot # | RRID # | Gene | Note | Donor in TEAseq | CD19-Depleted S | FACS Sort | Sample ID |
| HTO_A0251 | HTO_1 | GTCAACTCTTTAGCG | LNH-94 (CD298 Clone), 2M2 (B2M Clone) | BioLegend | 394601 | B342028 | AB_2750015 | ATP1B3, B2M | Costained with FACS | 15920-4 | No | All Mononuclear | PB1 |
| HTO_A0252 | HTO_2 | TGATGGCCTATTGGG | LNH-94 (CD298 Clone), 2M2 (B2M Clone) | BioLegend | 394603 | B369100 | AB_2750016 | ATP1B3, B2M | Costained with FACS | 15920-72 | No | All Mononuclear | PB2 |
| HTO_A0253 | HTO_3 | TTCCGCCTCTCTTTG | LNH-94 (CD298 Clone), 2M2 (B2M Clone) | BioLegend | 394605 | B364258 | AB_2750017 | ATP1B3, B2M | Costained with FACS | 15920-75 | No | All Mononuclear | PB3 |
| HTO_A0254 | HTO_4 | AGTAAGTTCAGCGTA | LNH-94 (CD298 Clone), 2M2 (B2M Clone) | BioLegend | 394607 | B368269 | AB_2750018 | ATP1B3, B2M | Costained with FACS | 15920-124 | No | All Mononuclear | PB4 |
| HTO_A0255 | HTO_5 | AAGTATCGTTTCGCA | LNH-94 (CD298 Clone), 2M2 (B2M Clone) | BioLegend | 394609 | B366838 | AB_2750019 | ATP1B3, B2M | Costained with FACS | 15920-4 | No | CD4+ Enriched | PB5 |
| HTO_A0256 | HTO_6 | GGTTGCCAGATGTCA | LNH-94 (CD298 Clone), 2M2 (B2M Clone) | BioLegend | 394611 | B344487 | AB_2750020 | ATP1B3, B2M | Costained with FACS | 15920-72 | No | CD4+ Enriched | PB6 |
| HTO_A0257 | HTO_7 | TGTCTTTCCTGCCAG | LNH-94 (CD298 Clone), 2M2 (B2M Clone) | BioLegend | 394613 | B343232 | AB_2750021 | ATP1B3, B2M | Costained with FACS | 15920-75 | No | CD4+ Enriched | PB7 |
| HTO_A0258 | HTO_8 | CTCCTCTGCAATTAC | LNH-94 (CD298 Clone), 2M2 (B2M Clone) | BioLegend | 394615 | B351089 | AB_2750022 | ATP1B3, B2M | Costained with FACS | 15920-124 | No | CD4+ Enriched | PB8 |
| HTO_A0259 | HTO_9 | CAGTAGTCACGGTCA | LNH-94 (CD298 Clone), 2M2 (B2M Clone) | BioLegend | 394617 | B359288 | AB_2750023 | ATP1B3, B2M | Costained with FACS | TC572 | No | All Mononuclear | TON1 |
| HTO_A0260 | HTO_10 | ATTGACCCGCGTTAG | LNH-94 (CD298 Clone), 2M2 (B2M Clone) | BioLegend | 394619 | B351604 | AB_2750024 | ATP1B3, B2M | Costained with FACS | TC574 | No | All Mononuclear | TON2 |
| HTO_A0296 | HTO_11 | CGATTGTAGACCTTT | LNH-94 (CD298 Clone), 2M2 (B2M Clone) | BioLegend | 394651 | B367989 | AB_2924590 | ATP1B3, B2M | Costained with FACS | TC569 | No | All Mononuclear | TON3 |
| HTO_A0262 | HTO_12 | TAACGACCAGCCATA | LNH-94 (CD298 Clone), 2M2 (B2M Clone) | BioLegend | 394623 | B363796 | AB_2750025 | ATP1B3, B2M | Costained with FACS | TC570 | No | All Mononuclear | TON4 |
| HTO_A0263 | HTO_13 | AAATCTCTCAGGCTC | LNH-94 (CD298 Clone), 2M2 (B2M Clone) | BioLegend | 394625 | B348202 | AB_2750026 | ATP1B3, B2M | Costained with FACS | TC572 | Yes | CD4+ Enriched | TON5 |
| HTO_A0264 | HTO_14 | CTGTATGTCCGATTG | LNH-94 (CD298 Clone), 2M2 (B2M Clone) | BioLegend | 394627 | B366625 | AB_2750027 | ATP1B3, B2M | Costained with FACS | TC574 | Yes | CD4+ Enriched | TON6 |
| HTO_A0265 | HTO_15 | TAAGATTCAGAGCGA | LNH-94 (CD298 Clone), 2M2 (B2M Clone) | BioLegend | 394629 | B343235 | AB_2750028 | ATP1B3, B2M | Costained with FACS | TC569 | Yes | CD4+ Enriched | TON7 |
| HTO_A0276 | HTO_16 | CTCAGTGCATTCTGG | LNH-94 (CD298 Clone), 2M2 (B2M Clone) | BioLegend | 394681 | B352817 | AB_2910455 | ATP1B3, B2M | Costained with FACS | TC570 | Yes | CD4+ Enriched | TON8 |
| ADT_A0072 | CD4 | TGTTCCCGCTCAACT | RPA-T4 | BioLegend | 300563 | B400905 | AB_2734247 | CD4 | Costained with FACS | All Samples | N/A | N/A | N/A |

| TotalSeqA ADT cocktail used for TEAseq | | | | | | | | | |
|---|---|---|---|---|---|---|---|---|---|
| TotalSeqA ID | Target | TotalSeq Barcode | Clone | Vendor | Catalog # | Lot # | RRID # | Gene | Note |
| ADT_A0148 | CD197 | AGTTCAGTCAACCGA | G043H7 | BioLegend | 353247 | B380062 | AB_2750357 | CCR7 | Spiked into cocktail |
| ADT_A0006 | CD86 | GTCTTTGTCAGTGCA | IT2.2 | BioLegend | 399907 | B342925 | AB_2888692 | CD86 | Included in HUC v1.0 |
| ADT_A0007 | CD274 | GTTGTCCGACAATAC | 29E.2A3 | BioLegend | 399907 | B342925 | AB_2888692 | CD274 | Included in HUC v1.0 |
| ADT_A0020 | CD270 | TGATAGAAACAGACC | 122 | BioLegend | 399907 | B342925 | AB_2888692 | TNFRSF14 | Included in HUC v1.0 |
| ADT_A0023 | CD155 | ATCACATCGTTGCCA | SKII.4 | BioLegend | 399907 | B342925 | AB_2888692 | PVR | Included in HUC v1.0 |
| ADT_A0024 | CD112 | AACCTTCCGTCTAAG | TX31 | BioLegend | 399907 | B342925 | AB_2888692 | NECTIN2 | Included in HUC v1.0 |
| ADT_A0026 | CD47 | GCATTCTGTCACCTA | CC2C6 | BioLegend | 399907 | B342925 | AB_2888692 | CD47 | Included in HUC v1.0 |
| ADT_A0029 | CD48 | CTACGACGTAGAAGA | BJ40 | BioLegend | 399907 | B342925 | AB_2888692 | CD48 | Included in HUC v1.0 |
| ADT_A0031 | CD40 | CTCAGATGGAGTATG | 5C3 | BioLegend | 399907 | B342925 | AB_2888692 | CD40 | Included in HUC v1.0 |
| ADT_A0032 | CD154 | GCTAGATAGATGCAA | 24-31 | BioLegend | 399907 | B342925 | AB_2888692 | CD40LG | Included in HUC v1.0 |
| ADT_A0033 | CD52 | CTTTGTACGAGCAAA | HI186 | BioLegend | 399907 | B342925 | AB_2888692 | CD52 | Included in HUC v1.0 |
| ADT_A0034 | CD3_UCHT1 | CTCATTGTAACTCCT | UCHT1 | BioLegend | 399907 | B342925 | AB_2888692 | CD3D | Included in HUC v1.0 |
| ADT_A0046 | CD8 | GCGCAACTTGATGAT | SK1 | BioLegend | 399907 | B342925 | AB_2888692 | CD8A | Included in HUC v1.0 |
| ADT_A0047 | CD56 | TCCTTTCCTGATAGG | 5.1H11 | BioLegend | 399907 | B342925 | AB_2888692 | NCAM1 | Included in HUC v1.0 |
| ADT_A0050 | CD19 | CTGGGCAATTACTCG | HIB19 | BioLegend | 399907 | B342925 | AB_2888692 | CD19 | Included in HUC v1.0 |
| ADT_A0052 | CD33 | TAACTCAGGGCCTAT | P67.6 | BioLegend | 399907 | B342925 | AB_2888692 | CD33 | Included in HUC v1.0 |
| ADT_A0053 | CD11c | TACGCCTATAACTTG | S-HCL-3 | BioLegend | 399907 | B342925 | AB_2888692 | ITGAX | Included in HUC v1.0 |
| ADT_A0058 | HLA.ABC | TATGCGAGGCTTATC | W6/32 | BioLegend | 399907 | B342925 | AB_2888692 | HLA-A | Included in HUC v1.0 |
| ADT_A0063 | CD45RA | TCAATCCTTCCGCTT | HI100 | BioLegend | 399907 | B342925 | AB_2888692 | PTPRC | Included in HUC v1.0 |
| ADT_A0064 | CD123 | CTTCACTCTGTCAGG | 6H6 | BioLegend | 399907 | B342925 | AB_2888692 | IL3RA | Included in HUC v1.0 |
| ADT_A0066 | CD7 | TGGATTCCCGGACTT | CD7-6B7 | BioLegend | 399907 | B342925 | AB_2888692 | CD7 | Included in HUC v1.0 |
| ADT_A0070 | CD49f | TTCCGAGGATGATCT | GoH3 | BioLegend | 399907 | B342925 | AB_2888692 | ITGA6 | Included in HUC v1.0 |
| ADT_A0071 | CD194 | AGCTTACCTGCACGA | L291H4 | BioLegend | 399907 | B342925 | AB_2888692 | CCR4 | Included in HUC v1.0 |
| ADT_A0072 | CD4_RPA.T4 | TGTTCCCGCTCAACT | RPA-T4 | BioLegend | 399907 | B342925 | AB_2888692 | CD4 | Included in HUC v1.0 |
| ADT_A0073 | CD44 | TGGCTTCAGGTCCTA | IM7 | BioLegend | 399907 | B342925 | AB_2888692 | CD44 | Included in HUC v1.0 |
| ADT_A0081 | CD14_M5E2 | TCTCAGACCTCCGTA | M5E2 | BioLegend | 399907 | B342925 | AB_2888692 | CD14 | Included in HUC v1.0 |
| ADT_A0083 | CD16 | AAGTTCACTCTTTGC | 3G8 | BioLegend | 399907 | B342925 | AB_2888692 | FCGR3A | Included in HUC v1.0 |
| ADT_A0085 | CD25 | TTTGTCCTGTACGCC | BC96 | BioLegend | 399907 | B342925 | AB_2888692 | IL2RA | Included in HUC v1.0 |
| ADT_A0087 | CD45RO | CTCCGAATCATGTTG | UCHL1 | BioLegend | 399907 | B342925 | AB_2888692 | PTPRC | Included in HUC v1.0 |
| ADT_A0088 | CD279 | ACAGCGCCGTATTTA | EH12.2H7 | BioLegend | 399907 | B342925 | AB_2888692 | PDCD1 | Included in HUC v1.0 |
| ADT_A0089 | TIGIT | TTGCTTACCGCCAGA | A15153G | BioLegend | 399907 | B342925 | AB_2888692 | TIGIT | Included in HUC v1.0 |
| ADT_A0090 | Isotype_MOPC.21 | GCCGGACGACATTAA | MOPC-21 | BioLegend | 399907 | B342925 | AB_2888692 | N/A | Included in HUC v1.0 |
| ADT_A0091 | Isotype_MOPC.173 | CTCCTACCTAAACTG | MOPC-173 | BioLegend | 399907 | B342925 | AB_2888692 | N/A | Included in HUC v1.0 |
| ADT_A0092 | Isotype_MPC.11 | ATATGTATCACGCGA | MPC-11 | BioLegend | 399907 | B342925 | AB_2888692 | N/A | Included in HUC v1.0 |
| ADT_A0095 | Isotype_RTK4530 | GATTCTTGACGACCT | RTK4530 | BioLegend | 399907 | B342925 | AB_2888692 | N/A | Included in HUC v1.0 |
| ADT_A0100 | CD20_2H7 | TTCTGGGTCCCTAGA | 2H7 | BioLegend | 399907 | B342925 | AB_2888692 | MS4A1 | Included in HUC v1.0 |
| ADT_A0101 | CD335 | ACAATTTGAACAGCG | 9E2 | BioLegend | 399907 | B342925 | AB_2888692 | NCR1 | Included in HUC v1.0 |
| ADT_A0124 | CD31 | ACCTTTATGCCACGG | WM59 | BioLegend | 399907 | B342925 | AB_2888692 | PECAM1 | Included in HUC v1.0 |
| ADT_A0127 | Podoplanin | GGTTACTCGTTGTGT | NC-08 | BioLegend | 399907 | B342925 | AB_2888692 | PDPN | Included in HUC v1.0 |
| ADT_A0134 | CD146 | CCTTGGATAACATCA | P1H12 | BioLegend | 399907 | B342925 | AB_2888692 | MCAM | Included in HUC v1.0 |
| ADT_A0136 | IgM | TAGCGAGCCCGTATA | MHM-88 | BioLegend | 399907 | B342925 | AB_2888692 | IGHM | Included in HUC v1.0 |
| ADT_A0138 | CD5 | CATTAACGGGATGCC | UCHT2 | BioLegend | 399907 | B342925 | AB_2888692 | CD5 | Included in HUC v1.0 |
| ADT_A0140 | CD183 | GCGATGGTAGATTAT | G025H7 | BioLegend | 399907 | B342925 | AB_2888692 | CXCR3 | Included in HUC v1.0 |
| ADT_A0141 | CD195 | CCAAAGTAAGAGCCA | J418F1 | BioLegend | 399907 | B342925 | AB_2888692 | CCR5 | Included in HUC v1.0 |
| ADT_A0142 | CD32 | GCTTCCGAATTACCG | FUN-2 | BioLegend | 399907 | B342925 | AB_2888692 | FCGR2A | Included in HUC v1.0 |
| ADT_A0143 | CD196 | GATCCCTTTGTCACT | G034E3 | BioLegend | 399907 | B342925 | AB_2888692 | CCR6 | Included in HUC v1.0 |
| ADT_A0144 | CD185 | AATTCAACCGTCGCC | J252D4 | BioLegend | 399907 | B342925 | AB_2888692 | CXCR5 | Included in HUC v1.0 |
| ADT_A0145 | CD103 | GACCTCATTGTGAAT | Ber-ACT8 | BioLegend | 399907 | B342925 | AB_2888692 | ITGAE | Included in HUC v1.0 |
| ADT_A0146 | CD69 | GTCTCTTGGCTTAAA | FN50 | BioLegend | 399907 | B342925 | AB_2888692 | CD69 | Included in HUC v1.0 |
| ADT_A0147 | CD62L | GTCCCTGCAACTTGA | DREG-56 | BioLegend | 399907 | B342925 | AB_2888692 | SELL | Included in HUC v1.0 |
| ADT_A0149 | CD161 | GTACGCAGTCCTTCT | HP-3G10 | BioLegend | 399907 | B342925 | AB_2888692 | KLRB1 | Included in HUC v1.0 |
| ADT_A0151 | CD152 | ATGGTTCACGTAATC | BNI3 | BioLegend | 399907 | B342925 | AB_2888692 | CTLA4 | Included in HUC v1.0 |
| ADT_A0152 | CD223 | CATTTGTCTGCCGGT | 11C3C65 | BioLegend | 399907 | B342925 | AB_2888692 | LAG3 | Included in HUC v1.0 |
| ADT_A0153 | KLRG1 | CTTATTTCCTGCCCT | SA231A2 | BioLegend | 399907 | B342925 | AB_2888692 | KLRG1 | Included in HUC v1.0 |
| ADT_A0154 | CD27 | GCACTCCTGCATGTA | O323 | BioLegend | 399907 | B342925 | AB_2888692 | CD27 | Included in HUC v1.0 |
| ADT_A0155 | CD107a | CAGCCCACTGCAATA | H4A3 | BioLegend | 399907 | B342925 | AB_2888692 | LAMP1 | Included in HUC v1.0 |
| ADT_A0156 | CD95 | CCAGCTCATTAGAGC | DX2 | BioLegend | 399907 | B342925 | AB_2888692 | FAS | Included in HUC v1.0 |
| ADT_A0158 | CD134 | AACCCACCGTTGTTA | Ber-ACT35 (ACT35) | BioLegend | 399907 | B342925 | AB_2888692 | TNFRSF4 | Included in HUC v1.0 |
| ADT_A0159 | HLA.DR | AATAGCGAGCAAGTA | L243 | BioLegend | 399907 | B342925 | AB_2888692 | HLA-DRA | Included in HUC v1.0 |
| ADT_A0160 | CD1c | GAGCTACTTCACTCG | L161 | BioLegend | 399907 | B342925 | AB_2888692 | CD1C | Included in HUC v1.0 |
| ADT_A0161 | CD11b | GACAAGTGATCTGCA | ICRF44 | BioLegend | 399907 | B342925 | AB_2888692 | ITGAM | Included in HUC v1.0 |
| ADT_A0162 | CD64 | AAGTATGCCCTACGA | 10.1 | BioLegend | 399907 | B342925 | AB_2888692 | FCGR1A | Included in HUC v1.0 |
| ADT_A0163 | CD141 | GGATAACCGCGCTTT | M80 | BioLegend | 399907 | B342925 | AB_2888692 | THBD | Included in HUC v1.0 |
| ADT_A0165 | CD314 | CGTGTTTGTTCCTCA | 1D11 | BioLegend | 399907 | B342925 | AB_2888692 | KLRK1 | Included in HUC v1.0 |
| ADT_A0167 | CD35 | ACTTCCGTCGATCTT | E11 | BioLegend | 399907 | B342925 | AB_2888692 | CR1 | Included in HUC v1.0 |
| ADT_A0168 | CD57 | AACTCCCTATGGAGG | QA17A04 | BioLegend | 399907 | B342925 | AB_2888692 | B3GAT1 | Included in HUC v1.0 |
| ADT_A0170 | CD272 | GTTATTGGACTAAGG | MIH26 | BioLegend | 399907 | B342925 | AB_2888692 | BTLA | Included in HUC v1.0 |
| ADT_A0171 | CD278 | CGCGCACCCATTAAA | C398.4A | BioLegend | 399907 | B342925 | AB_2888692 | ICOS | Included in HUC v1.0 |
| ADT_A0172 | CD275_B7.RP1 | GTTAGTGTTAGCTTG | 9F.8A4 | BioLegend | 399907 | B342925 | AB_2888692 | ICOSLG | Included in HUC v1.0 |
| ADT_A0174 | CD58 | GTTCCTATGGACGAC | TS2/9 | BioLegend | 399907 | B342925 | AB_2888692 | CD58 | Included in HUC v1.0 |
| ADT_A0176 | CD39 | TTACCTGGTATCCGT | A1 | BioLegend | 399907 | B342925 | AB_2888692 | ENTPD1 | Included in HUC v1.0 |
| ADT_A0179 | CX3CR1 | AGTATCGTCTCTGGG | K0124E1 | BioLegend | 399907 | B342925 | AB_2888692 | CX3CR1 | Included in HUC v1.0 |
| ADT_A0180 | CD24 | AGATTCCTTCGTGTT | ML5 | BioLegend | 399907 | B342925 | AB_2888692 | CD24 | Included in HUC v1.0 |
| ADT_A0181 | CD21 | AACCTAGTAGTTCGG | Bu32 | BioLegend | 399907 | B342925 | AB_2888692 | CR2 | Included in HUC v1.0 |
| ADT_A0185 | CD11a | TATATCCTTGTGAGC | TS2/4 | BioLegend | 399907 | B342925 | AB_2888692 | ITGAL | Included in HUC v1.0 |
| ADT_A0187 | CD79b | ATTCTTCAACCGAAG | CB3-1 | BioLegend | 399907 | B342925 | AB_2888692 | CD79B | Included in HUC v1.0 |
| ADT_A0189 | CD244 | TCGCTTGGATGGTAG | C1.7 | BioLegend | 399907 | B342925 | AB_2888692 | CD244 | Included in HUC v1.0 |
| ADT_A0206 | CD169 | TACTCAGCGTGTTTG | 7-239 | BioLegend | 399907 | B342925 | AB_2888692 | SIGLEC1 | Included in HUC v1.0 |
| ADT_A0214 | integrin.b7 | TCCTTGGATGTACCG | FIB504 | BioLegend | 399907 | B342925 | AB_2888692 | ITGB7 | Included in HUC v1.0 |
| ADT_A0215 | CD268 | CGAAGTCGATCCGTA | 11C1 | BioLegend | 399907 | B342925 | AB_2888692 | TNFRSF13C | Included in HUC v1.0 |
| ADT_A0216 | CD42b | TCCTAGTACCGAAGT | HIP1 | BioLegend | 399907 | B342925 | AB_2888692 | GP1BB | Included in HUC v1.0 |
| ADT_A0217 | CD54 | CTGATAGACTTGAGT | HA58 | BioLegend | 399907 | B342925 | AB_2888692 | ICAM1 | Included in HUC v1.0 |
| ADT_A0218 | CD62P | CCTTCCGTATCCCTT | AK4 | BioLegend | 399907 | B342925 | AB_2888692 | SELP | Included in HUC v1.0 |
| ADT_A0219 | CD119 | TGTGTATTCCCTTGT | GIR-208 | BioLegend | 399907 | B342925 | AB_2888692 | IFNGR1 | Included in HUC v1.0 |
| ADT_A0224 | TCR.AB | CGTAACGTAGAGCGA | IP26 | BioLegend | 399907 | B342925 | AB_2888692 | N/A | Included in HUC v1.0 |
| ADT_A0236 | Isotype_RTK2071 | ATCAGATGCCCTCAT | RTK2071 | BioLegend | 399907 | B342925 | AB_2888692 | N/A | Included in HUC v1.0 |
| ADT_A0237 | Isotype_G0114F7 | GGGAGCGATTCAACT | G0114F7 | BioLegend | 399907 | B342925 | AB_2888692 | N/A | Included in HUC v1.0 |
| ADT_A0238 | Isotype_RTK2758 | AAGTCAGGTTCGTTT | RTK2758 | BioLegend | 399907 | B342925 | AB_2888692 | N/A | Included in HUC v1.0 |
| ADT_A0240 | Isotype_RTK4174 | TCCAGGCTAGTCATT | RTK4174 | BioLegend | 399907 | B342925 | AB_2888692 | N/A | Included in HUC v1.0 |
| ADT_A0241 | Isotype_HTK888 | CCTGTCATTAAGACT | HTK888 | BioLegend | 399907 | B342925 | AB_2888692 | N/A | Included in HUC v1.0 |
| ADT_A0242 | CD192 | GAGTTCCCTTACCTG | K036C2 | BioLegend | 399907 | B342925 | AB_2888692 | CCR2 | Included in HUC v1.0 |
| ADT_A0246 | CD122 | TCATTTCCTCCGATT | TU27 | BioLegend | 399907 | B342925 | AB_2888692 | IL2RB | Included in HUC v1.0 |
| ADT_A0247 | CD267 | AGTGATGGAGCGAAC | 1A1 | BioLegend | 399907 | B342925 | AB_2888692 | TNFRSF13B | Included in HUC v1.0 |
| ADT_A0352 | FceRIa | CTCGTTTCCGTATCG | AER-37 (CRA-1) | BioLegend | 399907 | B342925 | AB_2888692 | FCER1A | Included in HUC v1.0 |
| ADT_A0353 | CD41 | ACGTTGTGGCCTTGT | HIP8 | BioLegend | 399907 | B342925 | AB_2888692 | ITGA2B | Included in HUC v1.0 |
| ADT_A0355 | CD137 | CAGTAAGTTCGGGAC | 4B4-1 | BioLegend | 399907 | B342925 | AB_2888692 | TNFRSF9 | Included in HUC v1.0 |
| ADT_A0357 | CD43 | GATTAACCAGCTCAT | CD43-10G7 | BioLegend | 399907 | B342925 | AB_2888692 | SPN | Included in HUC v1.0 |
| ADT_A0358 | CD163 | GCTTCTCCTTCCTTA | GHI/61 | BioLegend | 399907 | B342925 | AB_2888692 | CD163 | Included in HUC v1.0 |
| ADT_A0359 | CD83 | CCACTCATTTCCGGT | HB15e | BioLegend | 399907 | B342925 | AB_2888692 | CD83 | Included in HUC v1.0 |
| ADT_A0364 | CD13 | TTTCAACGCCCTTTC | WM15 | BioLegend | 399907 | B342925 | AB_2888692 | ANPEP | Included in HUC v1.0 |
| ADT_A0367 | CD2 | TACGATTTGTCAGGG | TS1/8 | BioLegend | 399907 | B342925 | AB_2888692 | CD2 | Included in HUC v1.0 |
| ADT_A0368 | CD226_11A8 | TCTCAGTGTTTGTGG | 11A8 | BioLegend | 399907 | B342925 | AB_2888692 | CD226 | Included in HUC v1.0 |
| ADT_A0369 | CD29 | GTATTCCCTCAGTCA | TS2/16 | BioLegend | 399907 | B342925 | AB_2888692 | ITGB1 | Included in HUC v1.0 |
| ADT_A0370 | CD303 | GAGATGTCCGAATTT | 201A | BioLegend | 399907 | B342925 | AB_2888692 | CLEC4C | Included in HUC v1.0 |
| ADT_A0371 | CD49b | GCTTTCTTCAGTATG | P1E6-C5 | BioLegend | 399907 | B342925 | AB_2888692 | ITGA2 | Included in HUC v1.0 |
| ADT_A0372 | CD61 | AGGTTGGAGTAGACT | VI-PL2 | BioLegend | 399907 | B342925 | AB_2888692 | ITGB3 | Included in HUC v1.0 |
| ADT_A0373 | CD81 | GTATCCTTCCTTGGC | 5A6 | BioLegend | 399907 | B342925 | AB_2888692 | CD81 | Included in HUC v1.0 |
| ADT_A0383 | CD55 | GCTCATTACCCATTA | JS11 | BioLegend | 399907 | B342925 | AB_2888692 | CD55 | Included in HUC v1.0 |
| ADT_A0384 | IgD | CAGTCTCCGTAGAGT | IA6-2 | BioLegend | 399907 | B342925 | AB_2888692 | IGHD | Included in HUC v1.0 |
| ADT_A0385 | CD18 | TATTGGGACACTTCT | TS1/18 | BioLegend | 399907 | B342925 | AB_2888692 | ITGB2 | Included in HUC v1.0 |
| ADT_A0386 | CD28 | TGAGAACGACCCTAA | CD28.2 | BioLegend | 399907 | B342925 | AB_2888692 | CD28 | Included in HUC v1.0 |
| ADT_A0389 | CD38_HIT2 | TGTACCCGCTTGTGA | HIT2 | BioLegend | 399907 | B342925 | AB_2888692 | CD38 | Included in HUC v1.0 |
| ADT_A0390 | CD127 | GTGTGTTGTCCTATG | A019D5 | BioLegend | 399907 | B342925 | AB_2888692 | IL7R | Included in HUC v1.0 |
| ADT_A0391 | CD45_HI30 | TGCAATTACCCGGAT | HI30 | BioLegend | 399907 | B342925 | AB_2888692 | PTPRC | Included in HUC v1.0 |
| ADT_A0393 | CD22 | GGGTTGTTGTCTTTG | S-HCL-1 | BioLegend | 399907 | B342925 | AB_2888692 | CD22 | Included in HUC v1.0 |
| ADT_A0394 | CD71 | CCGTGTTCCTCATTA | CY1G4 | BioLegend | 399907 | B342925 | AB_2888692 | TFRC | Included in HUC v1.0 |
| ADT_A0396 | CD26 | GGTGGCTAGATAATG | BA5b | BioLegend | 399907 | B342925 | AB_2888692 | DPP4 | Included in HUC v1.0 |
| ADT_A0398 | CD115 | AATCACGGTCCTTGT | 9-4D2-1E4 | BioLegend | 399907 | B342925 | AB_2888692 | CSF1R | Included in HUC v1.0 |
| ADT_A0404 | CD63 | GAGATGTCTGCAACT | H5C6 | BioLegend | 399907 | B342925 | AB_2888692 | CD63 | Included in HUC v1.0 |
| ADT_A0406 | CD304 | GGACTAAGTTTCGTT | 12C2 | BioLegend | 399907 | B342925 | AB_2888692 | NRP1 | Included in HUC v1.0 |
| ADT_A0407 | CD36 | TTCTTTGCCTTGCCA | 5-271 | BioLegend | 399907 | B342925 | AB_2888692 | CD36 | Included in HUC v1.0 |
| ADT_A0408 | CD172a | CGTGTTTAACTTGAG | 15-414 | BioLegend | 399907 | B342925 | AB_2888692 | SIRPA | Included in HUC v1.0 |
| ADT_A0419 | CD72 | CAGTCGTGGTAGATA | 3F3 | BioLegend | 399907 | B342925 | AB_2888692 | CD72 | Included in HUC v1.0 |
| ADT_A0420 | CD158 | TATCAACCAACGCTT | HP-MA4 | BioLegend | 399907 | B342925 | AB_2888692 | KIR2DL1 | Included in HUC v1.0 |
| ADT_A0446 | CD93 | GCGCTACTTCCTTGA | VIMD2 | BioLegend | 399907 | B342925 | AB_2888692 | CD93 | Included in HUC v1.0 |
| ADT_A0447 | CD200 | CACGTAGACCTTTGC | OX-104 | BioLegend | 399907 | B342925 | AB_2888692 | CD200 | Included in HUC v1.0 |
| ADT_A0575 | CD49a | ACTGATGGACTCAGA | TS2/7 | BioLegend | 399907 | B342925 | AB_2888692 | ITGA1 | Included in HUC v1.0 |
| ADT_A0576 | CD49d | CCATTCAACTTCCGG | 9F10 | BioLegend | 399907 | B342925 | AB_2888692 | ITGA4 | Included in HUC v1.0 |
| ADT_A0577 | CD73 | CAGTTCCTCAGTTCG | AD2 | BioLegend | 399907 | B342925 | AB_2888692 | NT5E | Included in HUC v1.0 |
| ADT_A0579 | CD9 | GAGTCACCAATCTGC | HI9a | BioLegend | 399907 | B342925 | AB_2888692 | CD9 | Included in HUC v1.0 |
| ADT_A0581 | TCR.Va7.2 | TACGAGCAGTATTCA | 3C10 | BioLegend | 399907 | B342925 | AB_2888692 | N/A | Included in HUC v1.0 |
| ADT_A0582 | TCR.Vd2 | TCAGTCAGATGGTAT | B6 | BioLegend | 399907 | B342925 | AB_2888692 | N/A | Included in HUC v1.0 |
| ADT_A0586 | CD354 | TAGCCGTTTCCTTTG | TREM-26 | BioLegend | 399907 | B342925 | AB_2888692 | TREM1 | Included in HUC v1.0 |
| ADT_A0590 | CD305_LAIR1 | ATTTCCATTCCCTGT | NKTA255 | BioLegend | 399907 | B342925 | AB_2888692 | LAIR1 | Included in HUC v1.0 |
| ADT_A0591 | LOX.1 | ACCCTTTACCGAATA | 15C4 | BioLegend | 399907 | B342925 | AB_2888692 | OLR1 | Included in HUC v1.0 |
| ADT_A0599 | CD158e1 | GGACGCTTTCCTTGA | DX9 | BioLegend | 399907 | B342925 | AB_2888692 | KIR3DL1 | Included in HUC v1.0 |
| ADT_A0817 | CD109 | CACTTAACTCTGGGT | W7C5 | BioLegend | 399907 | B342925 | AB_2888692 | CD109 | Included in HUC v1.0 |
| ADT_A0822 | CD142 | CACTGCCGTCGATTA | NY2 | BioLegend | 399907 | B342925 | AB_2888692 | F3 | Included in HUC v1.0 |
| ADT_A0830 | CD319 | AGTATGCCATGTCTT | 162.1 | BioLegend | 399907 | B342925 | AB_2888692 | SLAMF7 | Included in HUC v1.0 |
| ADT_A0845 | CD99 | ACCCGTCCCTAAGAA | 3B2/TA8 | BioLegend | 399907 | B342925 | AB_2888692 | CD99 | Included in HUC v1.0 |
| ADT_A0853 | CLEC12A | CATTAGAGTCTGCCA | 50C1 | BioLegend | 399907 | B342925 | AB_2888692 | CLEC12A | Included in HUC v1.0 |
| ADT_A0861 | CD151 | CTTACCTAGTCATTC | 50-6 | BioLegend | 399907 | B342925 | AB_2888692 | CD151 | Included in HUC v1.0 |
| ADT_A0864 | CD352 | AGTTTCCACTCAGGC | NT-7 | BioLegend | 399907 | B342925 | AB_2888692 | SLAMF6 | Included in HUC v1.0 |
| ADT_A0866 | CLEC1B | TGCCAGTATCACGTA | AYP1 | BioLegend | 399907 | B342925 | AB_2888692 | CLEC1B | Included in HUC v1.0 |
| ADT_A0867 | CD94 | CTTTCCGGTCCTACA | DX22 | BioLegend | 399907 | B342925 | AB_2888692 | KLRD1 | Included in HUC v1.0 |
| ADT_A0868 | IgE | GGATGTACCGCGTAT | MHE-18 | BioLegend | 399907 | B342925 | AB_2888692 | IGHE | Included in HUC v1.0 |
| ADT_A0870 | CD150 | GTCATTGTATGTCTG | A12 (7D4) | BioLegend | 399907 | B342925 | AB_2888692 | SLAMF1 | Included in HUC v1.0 |
| ADT_A0871 | CD162 | ATATGTCAGAGCACC | KPL-1 | BioLegend | 399907 | B342925 | AB_2888692 | SELPLG | Included in HUC v1.0 |
| ADT_A0872 | CD84 | CTCCCTAGTTCCTTT | CD84.1.21 | BioLegend | 399907 | B342925 | AB_2888692 | CD84 | Included in HUC v1.0 |
| ADT_A0894 | Ig.LightChain.k | AGCTCAGCCAGTATG | MHK-49 | BioLegend | 399907 | B342925 | AB_2888692 | IGKC | Included in HUC v1.0 |
| ADT_A0896 | CD85j | CCTTGTGAGGCTATG | GHI/75 | BioLegend | 399907 | B342925 | AB_2888692 | LILRB1 | Included in HUC v1.0 |
| ADT_A0897 | CD23 | TCTGTATAACCGTCT | EBVCS-5 | BioLegend | 399907 | B342925 | AB_2888692 | FCER2 | Included in HUC v1.0 |
| ADT_A0898 | Ig.LightChain.l | CAGCCAGTAAGTCAC | MHL-38 | BioLegend | 399907 | B342925 | AB_2888692 | N/A | Included in HUC v1.0 |
| ADT_A0902 | CD328 | CTTAGCATTTCACTG | 6-434 | BioLegend | 399907 | B342925 | AB_2888692 | SIGLEC7 | Included in HUC v1.0 |
| ADT_A0912 | GPR56 | GCCTAGTTTCCGTTT | CG4 | BioLegend | 399907 | B342925 | AB_2888692 | ADGRG1 | Included in HUC v1.0 |
| ADT_A0920 | CD82 | TCCCACTTCCGCTTT | ASL-24 | BioLegend | 399907 | B342925 | AB_2888692 | CD82 | Included in HUC v1.0 |
| ADT_A0923 | NKp80 | TATAGTTCCTCTGTG | 5D12 | BioLegend | 399907 | B342925 | AB_2888692 | KLRF1 | Included in HUC v1.0 |
| ADT_A0931 | CD131 | CTGCATGAGACCAAA | 1C1 | BioLegend | 399907 | B342925 | AB_2888692 | CSF2RB | Included in HUC v1.0 |
| ADT_A0935 | CD74 | CTGTAGCATTTCCCT | LN2 | BioLegend | 399907 | B342925 | AB_2888692 | CD74 | Included in HUC v1.0 |
| ADT_A0940 | CD116 | ATGGACAGTTCGTGT | 4H1 | BioLegend | 399907 | B342925 | AB_2888692 | CSF2RA | Included in HUC v1.0 |
| ADT_A0941 | CD37 | ACAGTCACTGGGCAA | M-B371 | BioLegend | 399907 | B342925 | AB_2888692 | CD37 | Included in HUC v1.0 |
| ADT_A0944 | CD101 | CTACTTCCCTGTCAA | BB27 | BioLegend | 399907 | B342925 | AB_2888692 | CD101 | Included in HUC v1.0 |
| ADT_A1018 | HLA.DR.DP.DQ | AGCTACGAGCAGTAG | Tü39 | BioLegend | 399907 | B342925 | AB_2888692 | HLA-DRA | Included in HUC v1.0 |
| ADT_A1046 | CD88 | GCCGCATGAGAAACA | S5/1 | BioLegend | 399907 | B342925 | AB_2888692 | C5AR1 | Included in HUC v1.0 |

**Table S3.** Illumina index barcodes used for sequencing TEAseq libraries.

| GEM Well | Library Name | 10x Genomics Index Name | Barcode #1 | Barcode #2 | Barcode #3 | Barcode #4 |
|---|---|---|---|---|---|---|
| 1 | ATAC 1 | SI-NA-A1 | AAACGGCG | CCTACCAT | GGCGTTTC | TTGTAAGA |
| 2 | ATAC 2 | SI-NA-B1 | AGGCTACC | CTAGCTGT | GCCAACAA | TATTGGTG |
| 3 | ATAC 3 | SI-NA-C1 | AGACTTTC | CCGAGGCA | GATGCAGT | TTCTACAG |
| 4 | ATAC 4 | SI-NA-D1 | AGCTGCGT | CAACCATC | GTGGAGCA | TCTATTAG |
| 5 | ATAC 5 | SI-NA-E1 | ACGAAAGC | CGCCCGTA | GTTTGCCT | TAAGTTAG |
| 6 | ATAC 6 | SI-NA-F1 | ATCGCTCC | CCGTACAG | GATAGGTA | TGACTAGT |
| 7 | ATAC 7 | SI-NA-G1 | ATGCGATT | CATATGCG | GGATACGA | TCCGCTAC |
| 8 | ATAC 8 | SI-NA-H1 | AAACTCAT | CGGGAGTA | GTCACAGG | TCTTGTCC |

| GEM Well | Library Name | 10x Genomics Index Name | Index (i7) Barcode | Index 2 (i5) Workflow B Barcode |
|---|---|---|---|---|
| 1 | GEX 1 | SI-TT-A10 | CGTGACATGC | TTTAGACCAT |
| 2 | GEX 2 | SI-TT-B10 | GCCCGATGGA | CTAGACGATT |
| 3 | GEX 3 | SI-TT-C10 | AGAATGGTTT | TCCCACCCTC |
| 4 | GEX 4 | SI-TT-D10 | ATGCGAATGG | CGACACTTGT |
| 5 | GEX 5 | SI-TT-E10 | CACAATCCCA | TTGTGGATAT |
| 6 | GEX 6 | SI-TT-F10 | CCGGCAACTG | TGTTAAACCG |
| 7 | GEX 7 | SI-TT-G10 | ACTTTACGTG | AGGGCGTTCA |
| 8 | GEX 8 | SI-TT-H10 | TTATCTAGGG | TAGAGCCTTT |

| GEM Well | Library Name | 10x Genomics Index Name | Index (i7) Barcode | Index 2 (i5) Workflow B Barcode |
|---|---|---|---|---|
| 1 | ADT 1 | N/A - Custom IDT Oligo | CAAGCAGAAGACGGCATACGAGATATTACTCGGTGACTGGAGTTCCTTGGCACCCGAGAATTCC*A | AATGATACGGCGACCACCGAGATCTACACTCTTTCCCTACACGACGCTC |
| 2 | ADT 2 | N/A - Custom IDT Oligo | CAAGCAGAAGACGGCATACGAGATGGCTCTGAGTGACTGGAGTTCCTTGGCACCCGAGAATTCC*A | AATGATACGGCGACCACCGAGATCTACACTCTTTCCCTACACGACGCTC |
| 3 | ADT 3 | N/A - Custom IDT Oligo | CAAGCAGAAGACGGCATACGAGATTCCGGAGAGTGACTGGAGTTCCTTGGCACCCGAGAATTCC*A | AATGATACGGCGACCACCGAGATCTACACTCTTTCCCTACACGACGCTC |
| 4 | ADT 4 | N/A - Custom IDT Oligo | CAAGCAGAAGACGGCATACGAGATCGCTCATTGTGACTGGAGTTCCTTGGCACCCGAGAATTCC*A | AATGATACGGCGACCACCGAGATCTACACTCTTTCCCTACACGACGCTC |
| 5 | ADT 5 | N/A - Custom IDT Oligo | CAAGCAGAAGACGGCATACGAGATGAGATTCCGTGACTGGAGTTCCTTGGCACCCGAGAATTCC*A | AATGATACGGCGACCACCGAGATCTACACTCTTTCCCTACACGACGCTC |
| 6 | ADT 6 | N/A - Custom IDT Oligo | CAAGCAGAAGACGGCATACGAGATATTCAGAAGTGACTGGAGTTCCTTGGCACCCGAGAATTCC*A | AATGATACGGCGACCACCGAGATCTACACTCTTTCCCTACACGACGCTC |
| 7 | ADT 7 | N/A - Custom IDT Oligo | CAAGCAGAAGACGGCATACGAGATGAATTCGTGTGACTGGAGTTCCTTGGCACCCGAGAATTCC*A | AATGATACGGCGACCACCGAGATCTACACTCTTTCCCTACACGACGCTC |
| 8 | ADT 8 | N/A - Custom IDT Oligo | CAAGCAGAAGACGGCATACGAGATCTGAAGCTGTGACTGGAGTTCCTTGGCACCCGAGAATTCC*A | AATGATACGGCGACCACCGAGATCTACACTCTTTCCCTACACGACGCTC |

| GEM Well | Library Name | 10x Genomics Index Name | Index (i7) Barcode | Index 2 (i5) Workflow B Barcode |
|---|---|---|---|---|
| 1 | HTO 1 | N/A - Custom IDT Oligo | CAAGCAGAAGACGGCATACGAGATCGAGTAATGTGACTGGAGTTCAGACGTGTGC | AATGATACGGCGACCACCGAGATCTACACTCTTTCCCTACACGACGCTC |
| 2 | HTO 2 | N/A - Custom IDT Oligo | CAAGCAGAAGACGGCATACGAGATTAATGCGCGTGACTGGAGTTCAGACGTGTGC | AATGATACGGCGACCACCGAGATCTACACTCTTTCCCTACACGACGCTC |
| 3 | HTO 3 | N/A - Custom IDT Oligo | CAAGCAGAAGACGGCATACGAGATCAGGACGTGTGACTGGAGTTCAGACGTGTGC | AATGATACGGCGACCACCGAGATCTACACTCTTTCCCTACACGACGCTC |
| 4 | HTO 4 | N/A - Custom IDT Oligo | CAAGCAGAAGACGGCATACGAGATTCCGCGAAGTGACTGGAGTTCAGACGTGTGC | AATGATACGGCGACCACCGAGATCTACACTCTTTCCCTACACGACGCTC |
| 5 | HTO 5 | N/A - Custom IDT Oligo | CAAGCAGAAGACGGCATACGAGATTCTCGCGCGTGACTGGAGTTCAGACGTGTGC | AATGATACGGCGACCACCGAGATCTACACTCTTTCCCTACACGACGCTC |
| 6 | HTO 6 | N/A - Custom IDT Oligo | CAAGCAGAAGACGGCATACGAGATAGCGATAGGTGACTGGAGTTCAGACGTGTGC | AATGATACGGCGACCACCGAGATCTACACTCTTTCCCTACACGACGCTC |
| 7 | HTO 7 | N/A - Custom IDT Oligo | CAAGCAGAAGACGGCATACGAGATTATAGCCTGTGACTGGAGTTCAGACGTGTGC | AATGATACGGCGACCACCGAGATCTACACTCTTTCCCTACACGACGCTC |
| 8 | HTO 8 | N/A - Custom IDT Oligo | CAAGCAGAAGACGGCATACGAGATATAGAGGCGTGACTGGAGTTCAGACGTGTGC | AATGATACGGCGACCACCGAGATCTACACTCTTTCCCTACACGACGCTC |

**Table S4.** Materials and reagents used for TEAseq.

| Item | Vendor | Catalog # |
|---|---|---|
| Human TruStain FcX | BioLegend | Cat# 422302, Lot# B362645, RRID# AB_2818986 |
| Chromium X Series Single Cell Analysis System | 10X Genomics | 1000326 |
| Chromium Next GEM Chip J Single Cell | 10X Genomics | 1000230 |
| Chromium Next GEM Single Cell Multiome ATAC/Gene Expression Reagent Bundle | 10X Genomics | 1000285 |
| Dual Index Kit TT Set A | 10X Genomics | 1000215 |
| Single Index Kit N Set A | 10X Genomics | 1000212 |
| Protector RNase Inhibitor | Millipore Sigma | 3335399001 |
| SPRIselect Reagent 5ml | Beckman Coulter | B23317 |
| Dynabeads MyOne Silane 5ml | Thermo Fisher Scientific | 37002D |
| Buffer EB | Qiagen | 19086 |
| Kapa HiFi HotStart Ready Mix | Kapa Biosystems | KM2602 |
| KAPA Library Quantification Kit for Illumina Platforms | Kapa Biosystems | KK4835 |
| Agilent High Sensitivity DNA Kit | Agilent Technologies | 5067-4626 |
| BSA ULTRA PURE (50MG/ML) 50MG | Invitrogen | AM2616 |
| 50% Glycerol 500ml | Teknova | G1800 |
| Digitonin 500mg | Thermo Fisher Scientific | 407565000 |
| Dimethyl sulfoxide, anhydrous, 99.8+%, Thermo Scientific™ 1L | Thermo Fisher Scientific | 43998M1 |
| Magnesium chloride, pure, Thermo Scientific™ 1Kg | Thermo Fisher Scientific | 223210010 |
| Fisher Science Education™ Sodium Chloride, Lab Grade 500g | Thermo Fisher Scientific | S25541 |
| Tris-HCL ph 7.4, a.k.a. Tris(hydroxymethyl)aminomethane hydrochloride 1L | Teknova | T1074 |
| Low TE Buffer (10 mM Tris-HCl, 0.1 mM EDT, pH 8.0) | Teknova | T0221 |
| Falcon Round Bottom Tubes Disposable Polystyrene Corning 5mL | Corning | 352054 |
| Eppendorf Safe-Lock Tubes 1.5 ml PCR clean colorless 500 tubes | Eppendorf Catalog | 022363212 |
| Falcon 5 ml Round Bottom Polystyrene Test Tube, with Cell Strainer Snap Cap | Corning | 352235 |
| Eppendorf twin.tec 96-Well PCR Plate Semi-Skirted | Eppendorf Catalog | 951020303 |
| Eppendorf twin.tec PCR 96-well plate, skirted | Eppendorf Catalog | 951020401 |
| Microseal 'B' Adhesive Seals | BioRad Sciences | MSB-1001 |
| SI-PCR-Oligo, ADT-Rev-AMP, ADT-i7, Additive HTO Primer, and HTO-i7 Primer Oligos | IDT | Custom oligo |

**Table S5.** Spectral flow cytometry antibody panel information.

| Marker Type Legend |
|---|
| Chemokine Receptor |
| Surface Staining |
| Intracellular Staining |
| Viability Stain |
| Fc Receptor Blocking |

| Panel A - Spectral flow sorting for TEAseq input samples | | | | | | | | |
|---|---|---|---|---|---|---|---|---|
| Detector | Fluorochrome | Marker | Manufacturer | Catalog # | Clone | Lot # | RRID # | Dilution |
| YG1 | PE | CD15 | BD Bioscencies | 562371 | W6D3 | 2305420 | AB_11154049 | 150 |
| YG9 | PE-Cy7 | CD4 | Biolegend | 317413 | OKT4 | B357838 | AB_571958 | 800 |
| R6 | Zombie NIR | Viability | Biolegend | 423106 | N/A | B349570 | N/A | 1000 |
| N/A | N/A | FcR Block | BD Biosciences | 564220 | Fc1 | 2244072 | AB_2869554 | 200 |

| Panel B - 30-color spectral flow paired profiling of TEAseq samples | | | | | | | | |
|---|---|---|---|---|---|---|---|---|
| Detector | Fluorochrome | Marker | Manufacturer | Catalog # | Clone | Lot # | RRID # | Dilution |
| UV2 | BUV395 | CD45RA | BD Biosciences | 740298 | HI100 | 1195622 | AB_2740037 | 2000 |
| UV7 | BUV496 | CD14 | BD Biosciences | 741200 | MϕP9 | 2038733 | AB_2870760 | 2500 |
| UV9 | BUV563 | CD25 | BD Biosciences | 612919 | 2A3 | 1141820 | AB_2870204 | 800 |
| UV10 | BUV615 | CCR4 | BD Biosciences | 613000 | 1G1 | 2032652 | AB_2870269 | 200 |
| UV11 | BUV661 | HLA-DR | BD Biosciences | 612980 | G46-6 | 1111954 | AB_2870252 | 1400 |
| UV14 | BUV737 | CD27 | BD Biosciences | 612829 | L128 | 1112895 | AB_2870151 | 800 |
| UV16 | BUV805 | CD3 | BD Biosciences | 612896 | UCHT1 | 2182332 | AB_2870184 | 300 |
| V1 | BV421 | PD-1 | Biolegend | 329920 | EH12.2H7 | B331240 | AB_10960742 | 200 |
| V3 | eFluor 450 | TIGIT | Invitrogen | 48-9500-42 | MBSA43 | 2288222 | AB_2637414 | 400 |
| V5 | BV480 | CD103 | BD Biosciences | 746472 | Ber-ACT8 | 1195624 | AB_2743774 | 800 |
| V7 | V500 | CD19 | BD Biosciences | 561125 | HIB19 | 1242317 | AB_10563208 | 600 |
| V8 | BV570 | CD8A | Biolegend | 301037 | RPA-T8 | B333844 | AB_10933259 | 600 |
| V10 | BV605 | CCR6 | Biolegend | 353419 | G034E3 | B338171 | AB_11124539 | 200 |
| V11 | Qdot 655 | CD38 | Invitrogen | Q22150 | HIT2 | 2303245 | AB_2556506 | 300 |
| V13 | BV711 | GATA3 | BD Biosciences | 565449 | L50-823 | 4004447 | AB_2739242 | 200 |
| V14 | BV750 | CXCR5 | BD Biosciences | 747111 | RF8B2 | 1195625 | AB_2871862 | 300 |
| V15 | BV786 | Ki67 | BD Biosciences | 563756 | B56 | 1169408 | AB_2732007 | 1200 |
| B2 | AF488 | FoxP3 | Invitrogen | 53-4776-42 | PCH101 | 2403272 | AB_11043133 | 300 |
| B3 | Spark Blue 550 | CD4 | Biolegend | 344656 | SK3 | B304843 | AB_2819979 | 700 |
| B9 | BB700 | CCR7 | BD Biosciences | 566437 | 3D12 | 1013583 | AB_2744306 | 300 |
| B10 | PerCP-ef710 | CD69 | Life Technologies | 46-0699-41 | FN50 | 2423801 | AB_2573693 | 600 |
| YG1 | PE | RORgt | BD Biosciences | 563081 | Q21-559 | 3304453 | AB_2686896 | 200 |
| YG3 | PE-CF594 | CD127 | BD Biosciences | 562397 | HIL-7R-M21 | 1060251 | AB_11154212 | 300 |
| YG5 | PE-Cy5 | CXCR3 | BD Biosciences | 561731 | 1C6 | 1116210 | AB_10892799 | 400 |
| YG9 | PE-Cy7 | T-bet | Biolegend | 644824 | 4B10 | B331455 | AB_2561761 | 600 |
| R1 | APC | CD16 | Biolegend | 302012 | 3G8 | B346619 | AB_314212 | 3000 |
| R2 | AF647 | Bcl-6 | BD Biosciences | 561525 | K112-91 | 2313154 | AB_10898007 | 200 |
| R4 | AF700 | CD40L | Biolegend | 310845 | 24-31 | B415906 | AB_2750052 | 400 |
| R6 | Zombie NIR | Viability | Biolegend | 423106 | N/A | B349570 | N/A | 1000 |
| R7 | APC-Fire 750 | ICOS | Biolegend | 313536 | C398.4A | B310666 | AB_2632923 | 500 |
| N/A | N/A | FcR Block | BD Biosciences | 564220 | Fc1 | 2244072 | AB_2869554 | 200 |

| Panel C - 25-color spectral flow staining of Gγ4 and Tfh features | | | | | | | | |
|---|---|---|---|---|---|---|---|---|
| **Detector** | **Fluorochrome** | **Marker** | **Manufacturer** | **Catalog #** | **Clone** | **Lot #** | **RRID #** | **Dilution** |
| UV2 | BUV395 | CD45RA | BD Biosciences | 740298 | HI100 | 1195622 | AB_2740037 | 2000 |
| UV7 | BUV496 | CD8A | BD Biosciences | 612942 | RPA-T8 | 2213972 | AB_2870223 | 800 |
| UV9 | BUV563 | CD25 | BD Biosciences | 612919 | 2A3 | 1141820 | AB_2870204 | 800 |
| UV10 | BUV615 | CCR4 | BD Biosciences | 613000 | 1G1 | 2251445 | AB_2870269 | 200 |
| UV14 | BUV737 | ICOS | BD Biosciences | 749665 | DX29 | 1349072 | AB_2873929 | 600 |
| UV16 | BUV805 | CD3 | BD Biosciences | 612896 | UCHT1 | 3311042 | AB_2870184 | 300 |
| V1 | BV421 | PD-1 | Biolegend | 329920 | EH12.2H7 | B331240 | AB_10960742 | 200 |
| V3 | eFluor 450 | TIGIT | Invitrogen | 48-9500-42 | MBSA43 | 2288222 | AB_2637414 | 400 |
| V5 | BV480 | CD27 | BD Biosciences | 566188 | L128 | 1133418 | AB_2739584 | 1000 |
| V7 | V500 | CD19 | BD Biosciences | 561125 | HIB19 | 1242317 | AB_10563208 | 600 |
| V10 | BV605 | CCR6 | Biolegend | 353419 | G034E3 | B386742 | AB_11124539 | 200 |
| V13 | BV711 | GATA3 | BD Biosciences | 565449 | L50-823 | 4004447 | AB_2739242 | 300 |
| V14 | BV750 | CXCR5 | BD Biosciences | 747111 | RF8B2 | 1195625 | AB_2871862 | 350 |
| B2 | AF488 | RORgt | BD Biosciences | 563621 | Q21-559 | 3268699 | AB_2738325 | 150 |
| B3 | Spark Blue 550 | CD4 | Biolegend | 344656 | SK3 | B413888 | AB_2819979 | 800 |
| B9 | BB700 | CCR7 | BD Biosciences | 566437 | 3D12 | 1013583 | AB_2744306 | 200 |
| B12 | RB744 | FOXP3 | BD Biosciences | 570472 | 259D/C7 | 4179232 | AB_3685765 | 150 |
| YG1 | PE | Gγ4 | NSJ | RQ6130 | Polyclonal | RQ6130-467i15 | AB_3713463 | 4500* |
| YG3 | PE-CF594 | CD127 | BD Biosciences | 562397 | HIL-7R-M21 | 1060251 | AB_11154212 | 300 |
| YG5 | PE-Cy5 | CXCR3 | BD Biosciences | 561731 | 1C6 | 1309502 | AB_10892799 | 400 |
| YG9 | PE-Cy7 | T-bet | Biolegend | 644824 | 4B10 | B331455 | AB_2561761 | 500 |
| R1 | APC | GPR183 | Biolegend | 368907 | SA313E4 | 368907 | AB_2632940 | 250 |
| R2 | AF647 | Bcl-6 | BD Biosciences | 561525 | K112-91 | 2313154 | AB_10898007 | 200 |
| R4 | AF700 | Ki-67 | BD Biosciences | 561277 | B56 | 9315354 | AB_10611571 | 1000 |
| R6 | Zombie NIR | Viability | Biolegend | 423106 | N/A | B349570 | N/A | 1000 |
| N/A | N/A | FcR Block | BD Biosciences | 564220 | Fc1 | 2244072 | AB_2869554 | 200 |

| Panel D - 24-color spectral flow analysis of Gγ4 and Tfh features in stimulated samples | | | | | | | | |
|---|---|---|---|---|---|---|---|---|
| **Detector** | **Fluorochrome** | **Marker** | **Manufacturer** | **Catalog #** | **Clone** | **Lot #** | **RRID #** | **Dilution** |
| UV2 | BUV395 | CD45RA | BD Biosciences | 740298 | HI100 | 1195622 | AB_2740037 | 2000 |
| UV7 | BUV496 | CD8A | BD Biosciences | 612942 | RPA-T8 | 2213972 | AB_2870223 | 1500 |
| UV9 | BUV563 | CD25 | BD Biosciences | 612919 | 2A3 | 1141820 | AB_2870204 | 1200 |
| UV10 | BUV615 | CCR4 | BD Biosciences | 613000 | 1G1 | 2251445 | AB_2870269 | 200 |
| UV14 | BUV737 | ICOS | BD Biosciences | 749665 | DX29 | 2038731 | AB_2873929 | 1000 |
| UV16 | BUV805 | CD3 | BD Biosciences | 612896 | UCHT1 | 3164898 | AB_2870184 | 1000 |
| V1 | BV421 | PD-1 | Biolegend | 329920 | EH12.2H7 | B331240 | AB_10960742 | 200 |
| V3 | eFluor 450 | TIGIT | Invitrogen | 48-9500-42 | MBSA43 | 2288222 | AB_2637414 | 400 |
| V5 | BV480 | CD27 | BD Biosciences | 566139 | L128 | 1263359 | AB_2739537 | 1000 |
| V7 | V500 | CD19 | BD Biosciences | 561125 | HIB19 | 1242317 | AB_10563208 | 600 |
| V10 | BV605 | CCR6 | Biolegend | 353419 | G034E3 | B386742 | AB_11124539 | 200 |
| V13 | BV711 | GATA3 | BD Biosciences | 565449 | L50-823 | 4004447 | AB_2739242 | 400 |
| V14 | BV750 | CXCR5 | BD Biosciences | 747111 | RF8B2 | 1195625 | AB_2871862 | 400 |
| B2 | AF488 | RORgt | BD Biosciences | 563621 | Q21-559 | 3268699 | AB_2738325 | 100 |
| B3 | Spark Blue 550 | CD4 | Biolegend | 344656 | SK3 | B413888 | AB_2819979 | 1500 |
| B12 | RB744 | FOXP3 | BD Biosciences | 570472 | 259D/C7 | 4179232 | AB_3685765 | 200 |
| YG1 | PE | Gγ4 | NSJ | RQ6130 | Polyclonal | RQ6130-467i15 | AB_3713463 | 5000* |
| YG3 | PE-CF594 | CD127 | BD Biosciences | 562397 | HIL-7R-M21 | 1060251 | AB_11154212 | 300 |
| YG5 | PE-Cy5 | CXCR3 | BD Biosciences | 561731 | 1C6 | 1309502 | AB_10892799 | 400 |
| YG9 | PE-Cy7 | T-bet | Biolegend | 644824 | 4B10 | B331455 | AB_2561761 | 600 |
| R1 | APC | GPR183 | Biolegend | 368907 | SA313E4 | 368907 | AB_2632940 | 250 |
| R2 | AF647 | Bcl-6 | BD Biosciences | 561525 | K112-91 | 2313154 | AB_10898007 | 200 |
| R4 | AF700 | Ki-67 | BD Biosciences | 561277 | B56 | 9315354 | AB_10611571 | 1000 |
| R6 | Zombie NIR | Viability | Biolegend | 423106 | N/A | B349570 | N/A | 1000 |
| N/A | N/A | FcR Block | BD Biosciences | 564220 | Fc1 | 3320439 | AB_2869554 | 200 |

** Note* - Listed dilutions of Gγ4 antibody solution refer to Gγ4-PE conjugate (Table S6, Supplementary Methods). After addition of Abcam Lightning-Link conjugation reagent and quencher solutions, the concentration of the input unconjugated Gγ4 antibody was ~

**Table S6.** Reagents used for spectral flow cytometry experiments.

| Miscellanous Flow Cytometry Reagents and Materials | | | |
|---|---|---|---|
| **Item** | **Manufacturer** | **Catalog #** | **Lot #** |
| eBioscience Fixation/Perm Diluent | Invitrogen | 00-5223-56 | 2555848 |
| Fixation/Permeabilization Concentrate | Invitrogen | 00-5123-43 | 2831007 |
| Permeabilization Buffer 10X | Invitrogen | 00-8333-56 | 2674243 |
| PBS pH 7.4 (1X) CMF | Gibco | 10010-023 | 2658922 |
| Brilliant Stain Buffer | BD Horizon | 566349 | 4011018 |
| UltraComp eBeads™ Plus Compensation Beads | Invitrogen | 01-3333-42 | 2897844 |
| Lightning-Link R-PE Conjugation Kit | Abcam | ab102918 | GR3455410-1 |

**Table S7.** Analysis software information.

| General Software | | |
|---|---|---|
| **Program** | **Version** | **Developer** |
| R | 4.4.0 | R Core Team |
| RStudio | 2024.04.0 | Posit Team |
| Prism | 10.2.2 (341) | GraphPad |
| FlowJo | 10.9.0 | Treestar |
| Xenium Explorer | 3.2.0 | 10x Genomics |
| Loupe Browser | 8.0.0 | 10x Genomics |

| RStudio Environment for TEAseq and Xenium Prime Data Analysis | | |
|---|---|---|
| **Package** | **Version** | **GitHub URL** |
| arrow | 18.1.0.1 | https://github.com/apache/arrow |
| AUCell | 1.26.0 | https://github.com/aertslab/AUCell |
| BiocGenerics | 0.50.0 | https://github.com/Bioconductor/BiocGenerics |
| BiocManager | 1.30.23 | https://github.com/Bioconductor/BiocManager |
| biomaRt | 2.60.0 | https://github.com/grimbough/biomaRt |
| BiocParallel | 1.38.0 | https://github.com/Bioconductor/BiocParallel |
| biovizBase | 1.52.0 | https://github.com/Bioconductor/biovizBase |
| BPCells | 0.3.0 | https://github.com/bnprks/BPCells |
| BSgenome.Hsapiens.UCSC.hg38 | 1.4.5 | https://github.com/Bioconductor/BSgenome.Hsapiens.UCSC.hg38 |
| caret | 6.0-94 | https://github.com/topepo/caret |
| cicero | 1.3.9 | https://github.com/cole-trapnell-lab/cicero-release |
| circlize | 0.4.16 | https://github.com/jokergoo/circlize |
| clusterProfiler | 4.12.6 | https://github.com/YuLab-SMU/clusterProfiler |
| ComplexHeatmap | 2.20.0 | https://github.com/jokergoo/ComplexHeatmap |
| data.table | 1.15.4 | https://github.com/Rdatatable/data.table |
| densityClust | 0.3.3 | https://github.com/thomasp85/densityClust |
| doMC | 1.3.8 | https://github.com/doParallel/doMC |
| dplyr | 1.1.4 | https://github.com/tidyverse/dplyr |
| EnhancedVolcano | 1.22.0 | https://github.com/kevinblighe/EnhancedVolcano |
| enrichplot | 1.24.2 | https://github.com/YuLab-SMU/enrichplot |
| EnsDb.Hsapiens.v86 | 2.99.0 | https://github.com/Bioconductor/EnsDb.Hsapiens.v86 |
| fastcluster | 1.2.6 | https://github.com/dmuellner/fastcluster |
| forcats | 1.0.0 | https://github.com/tidyverse/forcats |
| FNN | 1.1.4 | https://github.com/cran/FNN |
| future | 1.33.2 | https://github.com/HenrikBengtsson/future |
| GENIE3 | 1.26.0 | https://github.com/aertslab/GENIE3 |
| GenomeInfoDb | 1.40.0 | https://github.com/Bioconductor/GenomeInfoDb |
| GenomicFeatures | 1.56.0 | https://github.com/Bioconductor/GenomicFeatures |
| GenomicRanges | 1.56.0 | https://github.com/Bioconductor/GenomicRanges |
| ggh4x | 0.3.0 | https://github.com/teunbrand/ggh4x |
| ggnewscale | 0.5.0 | https://github.com/eliocamp/ggnewscale |
| ggplot2 | 3.5.1 | https://github.com/tidyverse/ggplot2 |
| ggraph | 2.2.1 | https://github.com/thomasp85/ggraph |
| ggrepel | 0.9.5 | https://github.com/slowkow/ggrepel |
| ggseqlogo | 0.2 | https://github.com/omarwagih/ggseqlogo |
| ggtern | 3.5.0 | https://github.com/ggtern/ggtern |
| ggVennDiagram | 1.5.2 | https://github.com/gaospecial/ggVennDiagram |
| glmGamPoi | 1.16.0 | https://github.com/const-ae/glmGamPoi |
| harmony | 1.2.0 | https://github.com/immunogenomics/harmony |
| igraph | 2.0.3 | https://github.com/igraph/igraph |
| IRanges | 2.38.0 | https://github.com/Bioconductor/IRanges |
| JASPAR2020 | 0.99.1 | https://github.com/GeorgiKrastev/JASPAR2020 |
| Matrix | 1.7-0 | https://github.com/MatrixOrg/Matrix |
| monocle3 | 1.3.7 | https://github.com/cole-trapnell-lab/monocle3 |
| motifmatchr | 1.26.0 | https://github.com/GreenleafLab/motifmatchr |
| motifDb | 1.46.0 | https://bioconductor.org/packages/MotifDb/ |
| msigdbr | 7.5.1 | https://github.com/igordot/msigdbr |
| Nebulosa | 1.14.0 | https://github.com/powellgenomicslab/Nebulosa |
| openxlsx | 4.2.8 | https://github.com/ycphs/openxlsx |
| org.Hs.eg.db | 3.19.1 | https://github.com/Bioconductor/org.Hs.eg.db |
| paletteer | 1.6.0 | https://github.com/EmilHvitfeldt/paletteer |
| pheatmap | 1.0.12 | https://github.com/raivokolde/pheatmap |
| presto | 1.0.0 | https://github.com/immunogenomics/presto |
| purrr | 1.0.2 | https://github.com/tidyverse/purrr |
| R2HTML | 2.3.4 | https://github.com/cran/R2HTML |
| RcisTarget | 1.23.1 | https://github.com/aertslab/RcisTarget |
| RColorBrewer | 1.1-3 | https://github.com/cran/RColorBrewer |
| readr | 2.1.5 | https://github.com/tidyverse/readr |
| readxl | 1.4.3 | https://github.com/tidyverse/readxl |
| reshape2 | 1.4.4 | https://github.com/hadley/reshape |
| Rfast2 | 0.1.5.4 | https://github.com/RfastOfficial/Rfast2 |
| rlang | 1.1.4 | https://github.com/r-lib/rlang |
| Rsamtools | 2.20.0 | https://github.com/Bioconductor/Rsamtools |
| rtracklayer | 1.64.0 | https://github.com/Bioconductor/rtracklayer |
| Rtsne | 0.17 | https://github.com/jkrijthe/Rtsne |
| S4Vectors | 0.42.0 | https://github.com/Bioconductor/S4Vectors |
| scales | 1.3.0 | https://github.com/r-lib/scales |
| scatterplot3d | 1.2 | https://github.com/cran/scatterplot3d |
| scDblFinder | 1.19.1 | https://github.com/plger/scDblFinder |
| SCENIC | 1.3.1 | https://github.com/aertslab/SCENIC |
| SCpubr | 2.0.2 | https://github.com/enblacar/SCpubr |
| Seurat | 5.1.0 | https://github.com/satijalab/seurat |
| SeuratObject | 5.0.2 | https://github.com/mojaveazure/seurat-object |
| SeuratWrappers | 0.3.5 | https://github.com/satijalab/seurat-wrappers |
| Signac | 1.13.0 | https://github.com/timoast/signac |
| SoupX | 1.6.2 | https://github.com/constantAmateur/SoupX |
| stringr | 1.5.1 | https://github.com/tidyverse/stringr |
| TFBSTools | 1.42.0 | https://github.com/ge11232002/TFBSTools |
| tibble | 3.2.1 | https://github.com/tidyverse/tibble |
| tidyr | 1.3.1 | https://github.com/tidyverse/tidyr |
| tidyselect | 1.2.1 | https://github.com/r-lib/tidyselect |
| tidyverse | 2.0.0 | https://github.com/tidyverse/tidyverse |
| TxDb.Hsapiens.UCSC.hg38.knownGene | 3.18.0 | https://github.com/Bioconductor/TxDb.Hsapiens.UCSC.hg38.knownGene |
| UCell | 2.8.0 | https://github.com/carmonalab/UCell |
| writexl | 1.5.4 | https://github.com/ycphs/writexl |

| RStudio Python Interface Environment for TEAseq Adjusted Mutual Information Data | | |
|---|---|---|
| **Package** | **Version** | **Source URL** |
| reticulate | 1.37.0 | https://github.com/rstudio/reticulate |
| python | 3.9 | https://www.python.org/ |
| scikit-learn | 1.6.1 | https://github.com/scikit-learn/scikit-learn |

| Python Environments for Spatial Transcriptomics Analysis | | |
|---|---|---|
| **Task** | **Task** | **Python Notebook / Script** |
| Identifying CN | neighbor | neighborhood_identification.ipynb |
| Merging GC CN and GC 'hole-filling' | neighbor | neighborhood_merge_GC.ipynb |
| WSI registration | wsireg | wsireg.py |
| Validation of WSI registration | stardist | registration_validation.ipynb |

| Spatial Cellular Neighborhood Analysis Python Environment ('neighbor') | | |
|---|---|---|
| **Code/Packages** | **Version** | **GitHub** |
| Original CN method | N/A | https://github.com/nolanlab/NeighborhoodCoordination/tree/master |
| python | 3.7 | https://www.python.org/ |
| pandas | 2.1.4 | https://github.com/pandas-dev/pandas |
| numpy | 1.26.4 | https://github.com/numpy/numpy |
| scikit-learn | 0.22.1 | https://github.com/scikit-learn/scikit-learn |
| seaborn | 0.9.0 | https://github.com/mwaskom/seaborn |
| matplotlib | 3.5.3 | https://github.com/matplotlib/matplotlib |
| scikit-image | 0.19.3 | https://github.com/scikit-image/scikit-image |
| scipy | 1.7.3 | https://github.com/scipy/scipy |

| Xenium Prime Whole Slide Image Registration Python Environment ('wsireg') | | |
|---|---|---|
| **Package** | **Version** | **GitHub** |
| pandas | 2.2.3 | https://github.com/pandas-dev/pandas |
| python | 3.11 | https://www.python.org/ |
| wsireg | 0.3.10 | https://github.com/NHPatterson/wsireg |

| Whole Slide Image Registration StarDist Validation Python Environment ('stardist') | | |
|---|---|---|
| **Package** | **Version** | **GitHub** |
| stardist | 0.9.1 | https://github.com/stardist/stardist |
| python | 3.1 | https://www.python.org/ |
| numpy | 1.26.4 | https://github.com/numpy/numpy |
| pandas | 2.1.4 | https://github.com/pandas-dev/pandas |
| scipy | 1.10.1 | https://github.com/scipy/scipy |
| tifffile | 2025.5.10 | https://github.com/cgohlke/tifffile |
| rasterio | 1.3.11 | https://github.com/rasterio/rasterio |
| matplotlib | 3.10.7 | https://github.com/matplotlib/matplotlib |

**Table S8.** Literature-based human Th17 gene signature (used in Fig. 2F).

| Gene |
|---|
| *RORA* |
| *RORC* |
| *CCR6* |
| *KLRB1* |
| *IL23R* |
| *IL17A* |
| *IL17F* |
| *IL22* |
| *IL26* |
| *CCL20* |
| *RUNX1* |
| *PALLD* |
| *TNFRSF8* |

**Table S9.** 100-plex custom add-on panel to 5001-plex 10x Genomics Xenium Prime 5K Human Pan Tissue and Pathways Panel.

| 1-25 | 26-50 | 51-75 | 76-100 |
|---|---|---|---|
| *AAK1* | *FOSL2* | *JUND* | *YPEL5* |
| *AHNAK* | *GFI1* | *KCNK5* | *ZFP36* |
| *ANXA1* | *GIMAP1* | *KLF2* | *THAP12* |
| *RNF111* | *GNAS* | *KLF3* | *RNF144A* |
| *ATP2B4* | *GNB1* | *KLF6* | *SCML4* |
| *BCAT1* | *GNB2* | *LINC00861* | *DOCK10* |
| *BHLHE40* | *GNB4* | *MAP3K8* | *TSC22D3* |
| *CADM1* | *GNB5* | *MMP17* | *P2RY8* |
| *CCL5* | *GNG10* | *NFKBIZ* | *PLAC8* |
| *CD48* | ***GNG4*** | *PCNX1* | *CAMK4* |
| *CD69* | *GNG5* | *PDE7A* | *CPQ* |
| *CD81* | *GNG7* | *CREM* | *CCL21* |
| *CHGB* | *GPR132* | *PTPN14* | *ARHGEF28* |
| *CRLF2* | *GPR183* | *RGS1* | *ZBTB7B* |
| *CXXC5* | *HAL* | *RGS3* | *TMSB10* |
| *DEF6* | *HECW2* | *RORA* | *XAF1* |
| *DRAIC* | *ID1* | *SCGB3A1* | *TRIM8* |
| *DUSP4* | *ID2* | *SKI* | *MYO7A* |
| *EGR1* | *ID3* | *SKIL* | *MAFG* |
| *EMP3* | *IL27* | *SMCO4* | *ACOXL* |
| *FABP5* | *IL27RA* | *ST8SIA1* | *CDKL2* |
| *FAM167A* | *IL32* | *TNFRSF4* | *CNIH3* |
| *FAM30A* | *IL7R* | *TRABD2A* | *NKG7* |
| *FAM43A* | *JUN* | *WNK2* | *NR4A2* |
| *FOSB* | *JUNB* | *XXYLT1* | *PREX1* |

**Supplementary Data Files**

**Data file S1. (separate file)**
Differentially expressed features between all TEAseq L1 clusters resolved by 3WNN (Fig. 1B, Fig. S2) and unimodal analyses (Fig. 1G, Fig. S5).

**Data file S2. (separate file)**
Differentially expressed features between all TEAseq L2 3WNN T cell clusters (Fig. 1H, Fig. S6), L2 Tfh core RNA and ATAC features relative to nonTfh (Fig. S7), and ADT expression in L2 Tfh versus nonTfh (Fig. 1I).

**Data file S3. (separate file)**
Differentially expressed features between all TEAseq L3 3WNN Tfh-like clusters (Fig. 2B, Fig. S8, Fig. S9), meta-analysis of L4 Tfh subset GC versus nonGC ATAC GeneActivity and RNA DEG data (Fig. 3B-3D), and ADT expression in *GNG4* RNA$^+$ vs RNA$^-$ tonsillar Tfh (Fig. 3H).

**Data file S4. (separate file)**
Xenium Prime H&E image registration metrics (Fig. S13), DEGs between all L1 clusters (Fig. 4H, Fig. S14), DEGs between all L2 nnCD4 T cell clusters (Fig. 4I, Fig. S15), DEGs between grouped L2 Tfh versus nonTfh clusters (Fig. 4J).

**Data file S5. (separate file)**
Xenium Prime ranking of percent Tfh localized to GC by each feature (Fig. 4L), three-way GC positioning and Tfh lineage identity comparisons (Fig. 4M), DEGs between *GNG4* RNA$^+$ vs RNA$^-$ GC Tfh (Fig. 4P), and DEGs between *FOXP3* RNA$^+$ vs RNA$^-$ GC Tfh (Fig. S18).

**Data file S6. (separate file)**
DEGs between all clusters in reanalysis of 10x Genomics tonsil Visium HD (Fig. S19B) and lymph node Visium v1 (Fig. S19G) data.

**Data file S7. (separate file)**
*GNG4* DAP-to-RNA links in TEAseq L3 Tfh object, DAPs between GC versus nonGC-like L4 Tfh object, DAPs between *GNG4* RNA$^+$ versus RNA$^-$ L4 Tfh, TF motifs enriched in *GNG4* RNA$^+$ Tfh DAPs, L3 Tfh SCENIC regulons containing *GNG4*, Cicero *cis*-coaccessibility networks for the *GNG4* locus in L4 Tfh (Fig. 5A-5D, 5H).

**Data file S8. (separate file)**
DEGs between TfhEff versus grouped nonTfh clusters and TfhEff versus TfhMem in reanalysis of publicly available data from mouse LCMV infection model (Fig. S21).

**Data file S9. (separate file)**
Linkage-disequilibrium analysis of rheumatoid arthritis fine-mapping GWAS variants and eQTL variants from stimulated human peripheral blood nnCD4 T cells (Fig. S23).

**Data file S10. (separate file)**
All other raw data supporting the findings of this study.
